# Population genetics of the coral *Acropora millepora*: Towards a genomic predictor of bleaching

**DOI:** 10.1101/867754

**Authors:** Zachary L. Fuller, Veronique J.L. Mocellin, Luke Morris, Neal Cantin, Jihanne Shepherd, Luke Sarre, Julie Peng, Yi Liao, Joseph Pickrell, Peter Andolfatto, Mikhail Matz, Line K. Bay, Molly Przeworski

**Author notes:** co-senior authors (Mikhail Matz, Line K. Bay, Molly Przeworski).

## Abstract

Although reef-building corals are rapidly declining worldwide, responses to bleaching vary both within and among species. Because these inter-individual differences are partly heritable, they should in principle be predictable from genomic data. Towards that goal, we generated a chromosome-scale genome assembly for the coral *Acropora millepora*. We then obtained whole genome sequences for 237 phenotyped samples collected at 12 reefs distributed along the Great Barrier Reef, among which we inferred very little population structure. Scanning the genome for evidence of local adaptation, we detected signatures of long-term balancing selection in the heat-shock co-chaperone *sacsin*. We further used 213 of the samples to conduct a genome-wide association study of visual bleaching score, incorporating the polygenic score derived from it into a predictive model for bleaching in the wild. These results set the stage for the use of genomics-based approaches in conservation strategies.

## Introduction

Anthropogenic global warming is transforming ecosystems worldwide, pressing species to rapidly respond to changing environments^1–4^. To what extent they will succeed is unclear: some may undergo range shifts or tolerate climate change through phenotypic plasticity. If there is sufficient heritable variation in traits that confer a fitness advantage in the new environment, others may adapt through evolutionary change^5–9^. Particularly devastating has been the impact of increased seawater temperatures on marine ecosystems — notably on coral reefs, which cover less than 1% of the ocean floor but are home to more than a quarter of its total biodiversity^10^. Ecological stress brought on by factors such as changes in temperature or salinity can break down the symbiotic relationship between reef-building corals and their intracellular photosynthetic dinoflagellates (family *Symbiodiniaceae*)^11^ and temperatures only slightly above long term maxima can result in a phenomenon known as mass bleaching, in which corals over vast areas lose their symbionts. Because these symbionts provide the majority of energy required to the coral host, prolonged periods of bleaching can eventually lead to the death of the coral. Increased temperatures can also cause heat damage directly to bleached corals by generating cellular damage and necrosis^12^. Over the last several years, mass bleaching events triggered by extreme seawater temperatures have decimated coral reefs worldwide^13,14^. For example, since 2014, the Great Barrier Reef (GBR) has lost nearly half of its coral cover^15^.

Despite the vulnerability of corals to warming, there is phenotypic variation in the bleaching and heat stress response both within and among coral species^16–18^. In the coral *Acropora millepora*, a common species on reefs across the Indo-Pacific, the variation in heat tolerance among aposymbiotic larvae was shown to be partly heritable^19^. Simulation studies have further suggested the potential for rapid adaptation in *A. millepora* populations^20^, and gene expression analyses have associated genes related to oxidative, extracellular, transport, and mitochondrial functions with elevated heat tolerance^19^. Previous studies remain limited by their reliance on transcriptomics or reduced representation sequencing, however, approaches that cannot distinguish causes from effects of bleaching or provide only a partial view of the genome, and by the lack of a high quality, publicly available reference genome. Thus, we remain far from a genetic understanding of bleaching and thermal tolerance, and from the ability to predict inter-individual variation in the responses.

To take first steps in that direction, we export an approach from agriculture and human genetics, where an increasing number of complex traits can be predicted from the results of genome-wide association studies (GWAS) alongside other predictors^21–23^. We illustrate how a similar approach can be applied to conservation biology, focusing on prediction of bleaching in wild populations of *A. millepora*. To this end, we constructed a highly contiguous chromosome-level *de novo* assembly of the *A. millepora* genome and generated high-coverage, whole-genome resequencing data for samples collected across 12 reefs on the GBR. We analyzed these data to search for signals of adaptation and to construct a reference haplotype panel for imputation of additional low coverage genomes. Using a set of 213 phenotyped and genotyped samples, we then carried out a GWAS for bleaching response in *A. millepora*, demonstrating how a predictive model for bleaching in natural populations can be built from genomic as well as environmental data.

### Inferring population structure and demographic history from genome-wide patterns of variation

We constructed a chromosome-scale *de novo* assembly of the diploid *A. millepora* genome using a combination of PacBio and 10X Chromium barcoded Illumina reads generated from DNA extracted from a single adult colony (Fig 1A, S1-S5, Table S1; SOM). We additionally sequenced two pools of aposymbiotic larvae and mapped these reads to our *de novo* assembly to remove symbiont contigs. In total, we assembled 475 Mb of sequence with chromosome-scale scaffolds for an N50 of 19.8 Mb and predicted 28,186 gene models, which include > 96% of core-eukaryotic single copy orthologs, making it the most complete coral genome constructed to date^24–29^.

**Figure 1:**
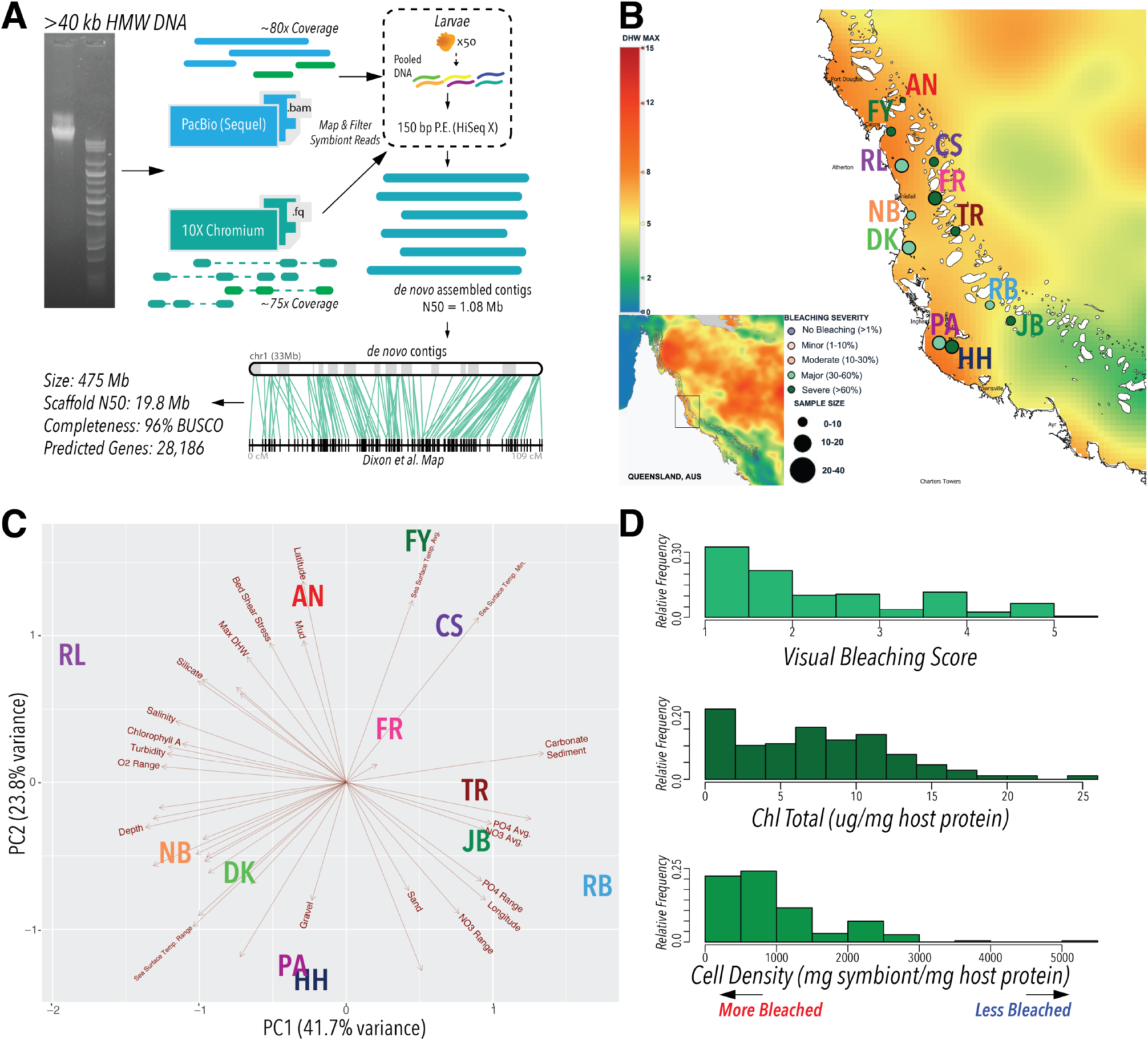
Genome assembly and sample collection for *A. millepora*. (**A**) A *de novo* assembly for the *A. millepora* genome was constructed using a hybrid sequencing approach. High molecular weight (> 40 kb) DNA was extracted from a single adult colony. The alignment of reads generated from two pools of aposymbiotic larvae were used to filter out symbiont contigs (SOM). After removing symbiont contigs, the set of *de novo* assembled host coral contigs were aligned to previously published linkage maps (19, 64) to create a chromosome-scale assembly. (**B**) A total of 253 individuals were collected from across 12 reefs on the GBR in 2017. The size of the circles represent the number of individuals collected at each site and are colored based on the bleaching severity at each reef. The maximum degree heating weeks (DHW) is shown across the region from which samples were collected (and across the GBR in the inset). Each label is colored arbitrarily, but consistent with labels presented in other figures. (**C**) A principal components analysis (PCA) was performed for 40 environmental and spatial variables for each reef, with abbreviations in bold representing their location. The component loads for each environmental variable projected on the first two PCs are depicted with arrows. (**D**) The distribution of visual scores, chlorophyll abundance standardized by host coral protein content, and standardized symbiont cell densities among the collected individuals for which phenotype measurement was possible.

To investigate genetic variation associated with bleaching response, we collected branch fragments from a total of 253 individual *A. millepora* colonies during peak bleaching in March 2017 from 12 reefs (Fig 1B, Table S2) distributed along an approximate 300 km stretch of the Central GBR. For each colony, we recorded the bleaching level visually using the six point coral health chart to the nearest half increment^30^, then later estimated photosynthetic pigment concentration and symbiont cell density. Ecological conditions were measured by 40 environmental and spatial variables that varied among reefs^31^, with the strongest axes of variation being longitude and latitude (Fig 1C). Mean monthly sea surface temperatures varied by more than 0.5°C across the range of sampled reefs, and the depth of colonies from which fragments were collected varied from 1.2 to 7.2 m. Individual colonies sampled presented a range of bleaching phenotypes, both within and between reefs, recorded at the time of collection as a standard visual score and later directly measured as total chlorophyll content standardized by host protein levels and symbiont cell density (Fig 1D).

From this collection of samples, we resequenced 48 whole genomes of diploid individuals (four from each of the 12 reefs) at high coverage (> 20× for all, 36 samples > 100×; Fig S6, Table S3) to examine the extent of population structure, characterize the demographic history, and scan for signals of adaptation. We mapped reads from all individuals to our *de novo* genome assembly and called approximately 6.8 million biallelic single nucleotide polymorphisms (SNPs) after applying stringent quality filters (SOM). We validated these filtered SNP calls by independently amplifying and sequencing 10 intergenic regions in eight samples at extremely high coverage (> 1400×; Fig S8, Table S4), observing an overall genotype concordance of 99.6% between regions sequenced by the two approaches (SOM). After an initial principal components analysis (PCA) and test for outliers using identity-by-state distance, we removed four individuals that were likely the result of sample mis-identification at the time of collection (or possibly were cryptic species), bringing the total number of high-coverage resequenced genomes to 44. These remaining genomes all appear very distantly and approximately equidistantly-related (Fig S7, SOM).

From this set of 44 unrelated individuals, we estimated the decay of pairwise linkage disequilibrium (*r*^2^) as a function of physical distance genome-wide. We observed that, on average, *r*^2^ falls below 0.05 after ~ 15 kb (Fig 2A). We also estimated the nucleotide diversity (as *π*, the mean pairwise difference per base pair) in nonoverlapping intergenic regions of 1 kb, which yielded an average *π* = 0.363% (Fig 2B). Assuming a mutation rate of 4 × 10^−9^ per base pair per generation^20,34^, this value of diversity corresponds to a long-term effective population size (*N_e_*) of around 2.26 × 10^5^. Applying the pairwise sequential Markovian coalescent (PSMC)^33^ to each genome, we inferred how *N_e_* has changed over the last ~ 10^6^ generations (~ 5 million years). All samples display a highly similar demographic trajectory, with the inferred *N_e_* showing a steady decline over the last million generations. Qualitatively similar results were obtained when demographic histories for the same samples were inferred jointly using the multiple sequentially Markovian coalescent (MSMC; Fig S10)^35^. A similar pattern of decline in *N_e_* has been observed in other corals, including a number of *Acropora* and *Orbicella* species^20,26,36^.

**Figure 2:**
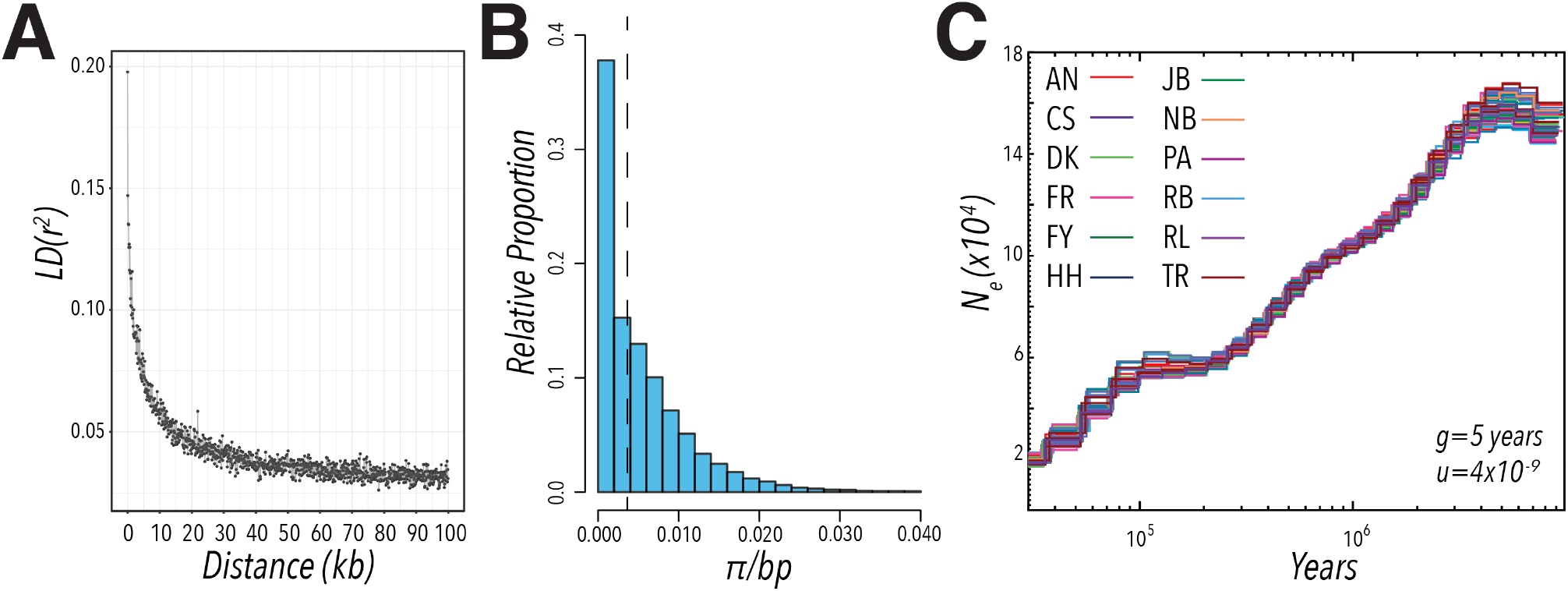
Properties of genetic variation and inferred demographic history in sampled *A. millepora*. (**A**) The decay of linkage disequilibrium (LD), measured as the squared genotypic correlation coefficient (*r*^2^), was estimated for a randomly chosen 1% of SNPs using all sequenced samples. Each point represents the mean *r*^2^ in bins of 100 bp. (**B**) Nucleotide diversity was measured by the average pairwise differences per bp (*π*)^32^ for SNPs in 1 kb windows, using intergenic regions genome-wide. (**C**) Effective population sizes inferred from 44 sequenced *A. millepora* genomes using PSMC^33^, assuming a generation time of 5 years and a mutation rate of 4 × 10^−9^ per base pair per generation^20,34^. Note that this approach provides very little information about the recent past (less than ~ 20 Kya).

The similar long-term demographic histories inferred across all individuals suggested minimal population structure and high connectivity among the 12 sampled reefs. Accordingly, there is no detectable relationship between geographic distance and *F_ST_* between pairs of sampled reefs (Fig 3A). A principal component analysis of SNPs in approximate linkage equilibrium further reveals no obvious clusters (Fig 3B). Moreover, no significant relationships were detected between either of the first two eigenvectors and any of the 40 environmental or spatial variables for each reef, indicating that any population structure that is present is at most very weakly associated with our measured environmental variables. The lack of population structure can be visualized by estimating and plotting relative effective migration rates. Using the program EEMS^37^, we inferred homogenous migration rates and found no obvious barriers to gene flow among sampled reefs (Fig 3C). This pattern of gene flow across large geographic distances is consistent with previous findings in *A. millepora* and is presumably a reflection of its broadcast spawning mode of reproduction^10,17,19,20,38^.

**Figure 3:**
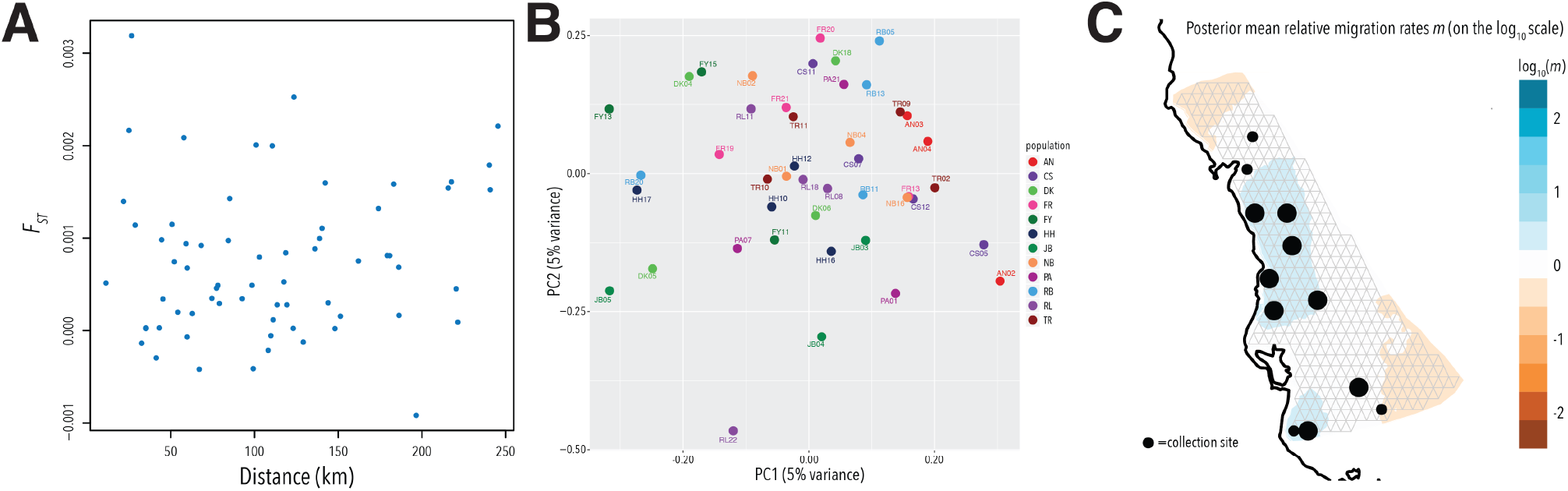
Characterizing population structure and gene flow across 12 reefs. (**A**) There is no discernable relationship between geographic distance and genetic differentiation (measured as *F_ST_*) across pairwise comparisons of sampled reefs. (**B**) A PCA using LD-pruned genome-wide SNPs for the 44 resequenced high-coverage genomes. Each point is color-coded by the reef from which the individual was sampled. (**C**) EEMS^37^ was used to estimate and visualize the relative effective migration surface using LD pruned and common (MAF> 5%) genome-wide SNPs for the 44 resequenced high-coverage genomes. The size of the dots reflect the number of individuals sequenced from each reef.

### A scan for adaptation points to a heat-shock protein co-chaperone

The near absence of genetic differentiation across reefs indicates that there is little population structure over hundreds of kilometers. Yet environmental conditions differ among reefs, notably in terms of thermal regimes (Fig 1B). Since adults only reproduce with those individuals in their vicinity while larvae can disperse over much larger ranges^16,20^, larvae may therefore experience strong selection each generation, as they settle on a heterogeneous reefscape. Models suggest that this scenario of strong, spatially varying selection in the face of continual migration can lead to the maintenance of locally beneficial alleles over long periods of time^39,40^. We therefore searched for signals of adaptation in the 44 resequenced genomes by scanning for genomic regions displaying the high levels of diversity that would result from such long-lived balancing selection. Genome-wide, the most extreme values (*π* > 5.0%, measured in 1 kb windows; Fig 4A) fell over a region containing a single gene, which is orthologous to *sacsin* in *A. digitifera*. The same genic region also has a highly elevated value of the *h*_12_ statistic^41^, which measures the frequencies of the two most common haplotypes (Fig 4B). Between the two most common haplotypes there were 20 fixed synonymous differences and 29 fixed nonsynonymous differences (Fig 4A). Confirming that this pattern of relatedness is unusual, a PCA for the region stands out by comparison to the rest of the genome (as assessed with *lostruct*^42^, see SOM; Fig S11). These summary statistics point to an unusually deep genealogy with long internal branches at *sacsin*: indeed, the divergence of two haplotypes even predates the species split with *A. tenuis* and *A. digitifera* (Fig 4C). To ensure that these findings were not artifactual, we confirmed the presence of the two diverged haplotypes by independently sequencing and assembling overlapping 2 kb amplicons spanning the entire gene for a subset of four samples (Fig S13-14, Table S6-7; SOM). Moreover, there was no significant elevation in coverage surrounding *sacsin* for any sample to suggest the presence of a paralog or pseudogene (Fig S12).

**Figure 4:**
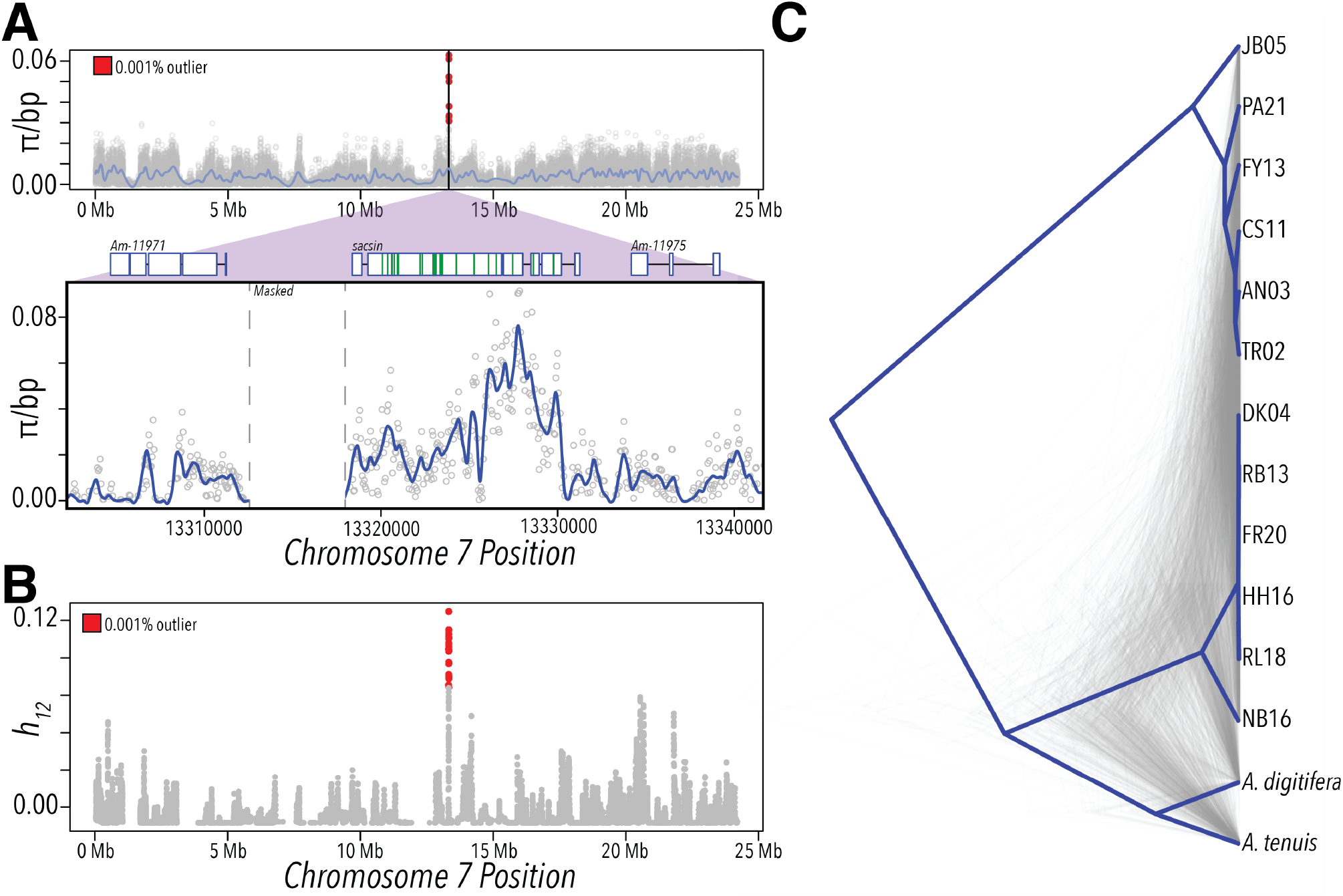
Genomic scans for local adaptation detect a signal at *sacsin*. (**A**) In the top panel, values of pairwise nucleotide diversity (*π*) estimated in sliding windows of 1 kb and a step-size of 500 bp across chromosome 7, with points colored in red representing regions in the top 0.001% of values genome-wide. A loess-smoothed trend line of *π* across windows is indicated in purple. The peak with the most extreme values falls in the *sacsin* gene region. In the bottom panel, a close-up view of *π* estimated across a region of chromosome 7 surrounding the *sacsin* gene, which includes all of the outlier windows denoted with red in the top panel. *π* per bp was calculated in sliding windows of 100 bp (shown in gray dots); the loess-smoothed line is shown in blue. The predicted gene structure of *sacsin* is indicated above, as well as the flanking upstream and downstream genes. The 29 non-synonymous differences fixed between samples of the two most common haplotypes in *sacsin* are denoted with green lines. Gray vertical dashed lines delimit a region that was masked from variant calling because of predicted repetitive elements. (**B**) The *h*_12_ summary statistic^41^, which measures the frequency of the two most common haplotypes, is plotted across chromosome 7. Red dots represent *h*_12_ values in the top 0.001% genome-wide. The peak with the most extreme values falls in the *sacsin* gene region. (**C**) A gene tree for the central 1 kb region in *sacsin* is shown in dark blue; it was constructed for one randomly chosen haplotype from a randomly selected individual from each of the 12 sampled reefs. The gene tree is rooted to aligned sequences from the reference genomes for *A. digitifera*^24^ and *A. tenuis* (draft assembly from www.reefgenomics.org). Shown in gray are gene trees for 1000 randomly sampled 1 kb regions from across the genome for the same individuals.

*Sacsin* is a co-chaperone for the heat-shock protein *Hsp70* and contains multiple regions of homology to *Hsp90*^43,44^. Our analyses reveal that variation in *sacsin* has been maintained for an atypically long time (*i.e*., tens of millions of years), indicating that the gene is a target of balancing selection and hence of functional importance in the wild. The source of balancing selection is unclear, as there is no discernable association of the haplotype frequencies with genetic or current environmental PCs (*P* > 0.15, Table S5). However, *sacsin* has been shown to be significantly upregulated when exposed to elevated temperatures in the laboratory, in multiple coral species from the genus *Acropora* and *Pocillipora*^45–48^, indicating that variation in the gene is plausibly associated with heat response in natural populations.

### A genome-wide association study for bleaching

Using the 44 high-coverage genomes, we used read-backed phasing^49^ to construct a reference haplotype panel and impute genotypes in low-pass sequences of additional samples. We generated a mean of ~ 1.5× coverage whole-genome data^50^ for 193 individuals after applying filters based on quality, read depth and the proportion of missing data (Fig S16). Of these samples, 34 had also been sequenced at high coverage, allowing us to assess the accuracy of our imputation approach. In the 34 samples, the overall correlation between sequenced and imputed genotypes was > 94% for SNPs with minor allele frequency > 0.05 and genotype probability ≥ 0.95 (Fig S17), an imputation accuracy similar to what is observed in human data using reference panels of similar size^51,52^.

Combining the low coverage genomes with the 44 high coverage ones, we obtained genotype calls at ~ 6.8 million sites in a total of 237 total samples. After excluding outliers that were likely misidentifications^53^, individuals with high levels of relatedness, and those with missing phenotype data (Fig S18-19; see SOM), our final sample size was 213. Consistent with our previous results, in the larger sample we again observed minimal population structure, high gene flow, and a peak of elevated genetic diversity at *sacsin* (Fig S20). While the low population structure is helpful for GWAS, it makes precise estimates of SNP heritability unattainable without substantially larger sample sizes (see SOM;^54,55^).

The sequencing also yielded data from the symbiont *Symbiodiniaceae*, of which multiple species are known to associate with *A. millepora*^56,57^. Since intracellular symbiont DNA was extracted and sequenced simultaneously with the coral host, we used our genomic sequencing approach to characterize the symbiont species present in each sample^11,53^. The abundance of symbiont reads in each sample was significantly different between quartiles of visual scores (Fig 5A, *P* = 2.6 × 10^−7^ by a Kruskal-Wallis test), as well as highly correlated with total chlorophyll content or symbiont cell density (Fig S22), suggesting that the relative abundance of reads from the symbiont can serve as a quantitative measure of bleaching. We further determined the composition of symbiont types for each sample by calculating the relative proportion of reads mapping to available draft genomes. In our samples, individuals dominated by *Durusdinium* had significantly higher visual scores on average (Fig 5B, *P* = 2.46 × 10^−7^ by a Mann-Whitney *U* test), consistent with previous findings that *Durusdinium* (formerly Symbiodinium Clade D) is associated with substantially higher thermal tolerance (but often lower growth rates) across diverse coral genera in the wild^11,58–60^.

**Figure 5:**
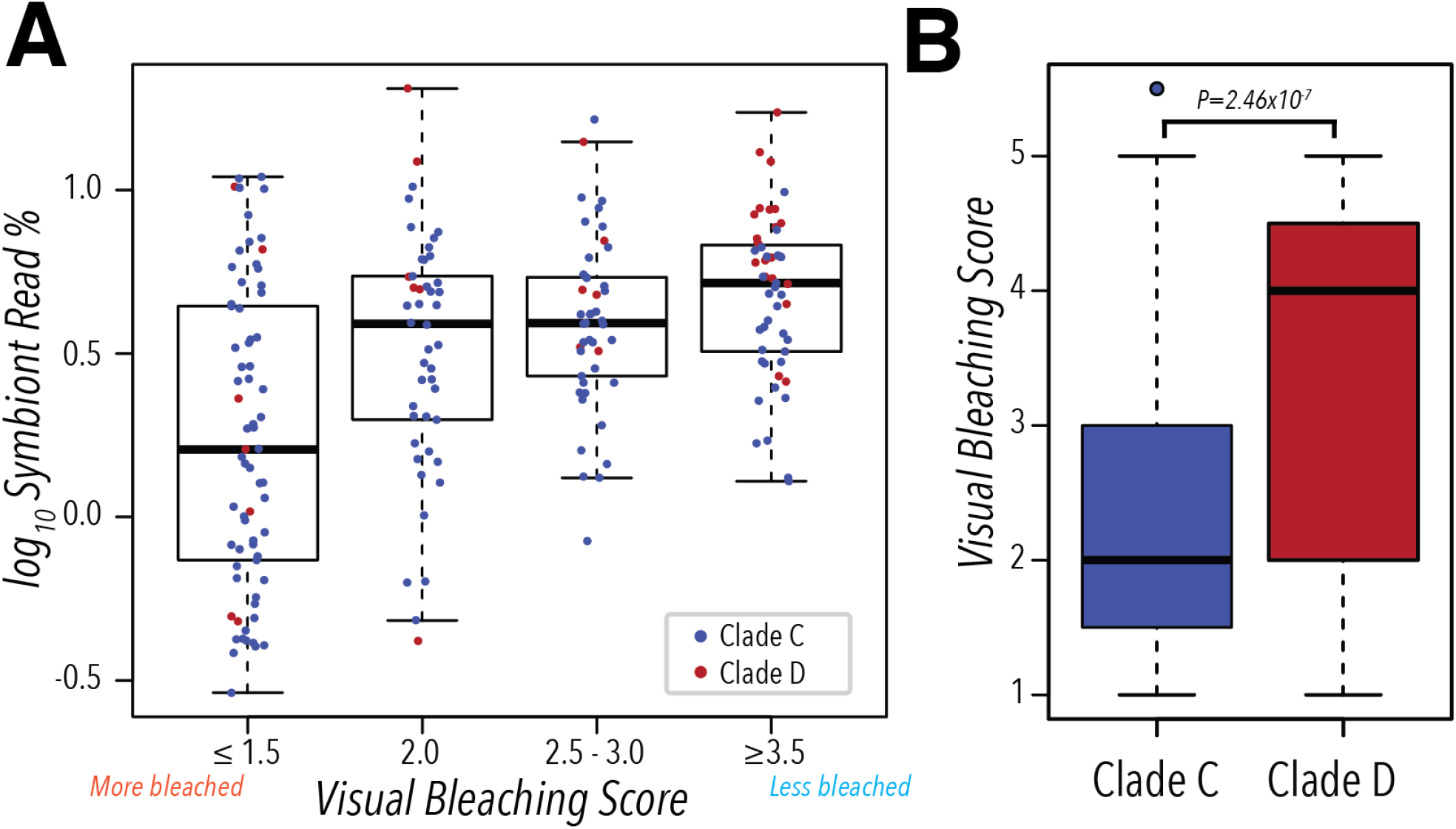
Relationship between symbiont species and bleaching response revealed by high-throughput sequencing. (**A**) The total proportion of reads mapping to draft symbiont genomes was measured for each sample. Bins for the visual scores were chosen such that approximately equal numbers of samples were included in each. Points are color-coded according to the dominant symbiont genus found in each sample. The y-axis is shown on the log_10_ scale. (**B**) Samples dominated by *Durusdinium* (formerly *Symbiodinium* Clade D) have a significantly higher (*P* = 2.46 × 10^−7^ by a two-tailed Mann-Whitney *U* test) visual score compared to samples dominated by *Cladocopium* (formerly *Symbiodinium* Clade C).

Given these data from corals and their symbionts, we used a linear mixed model (LMM) to test for additive effects of SNPs (with minor allele frequencies > 5%) on the quantile-normalized visual score^61,62^, including as covariates the top four PCs of environmental variables (which together account for > 85% of the total environmental variance), the first two genetic PCs, batch effects (the collection date and sequencing batch), the collection depth^63^, and the proportion of *Durusdinium* reads relative to all symbiont reads (see SOM). We also conducted similar GWAS for total chlorophyll content and symbiont cell density (Fig S23). The three measures of bleaching are highly correlated (Spearman’s correlation coefficient among pairs of phenotypes > 0.6; Fig S21, see SOM); because we are missing phenotype data for chlorophyll content (such that *n* = 190) and symbiont cell density (*n* = 172), we focused on the standard visual score (*n* = 213).

We determined a genome-wide significance threshold of *P* = 4.3 × 10^−8^ for the quantile-normalized visual score by considering the distribution of *P*-values obtained from 10^5^ permutation tests (Fig S29; see SOM). By this approach, no single SNP in our GWAS is genome-wide significant, indicating the absence of common, major effect loci associated with variation in bleaching. In particular, the minimum *P*-value in the *sacsin* gene is high (0.01). For related phenotypes of total chlorophyll content and symbiont cell density, however, SNPs in *sacsin* are among the top signals (minimal *P* values are 5.05 × 10^−5^ and 2.38 × 10^−5^, respectively), suggesting that the lack of signal in the visual score GWAS may reflect lack of power. For visual score, the top peaks were a region encompassing two genes on chromosome 13 and one with three genes on chromosome 14 (Figure 6A; also detected using the software BIMBAM^64^, see Fig S28). These associations remain to be confirmed, but it is worth noting that among the three genes on chromosome 14 is *malate synthase*, a key component of the glyoxylate cycle, a pathway not found in metazoans outside of cnidarians^65^. *Malate synthase* is up-regulated in response to thermal stress in *A. palmata* and implicated in the coral stress response^66,67^.

**Figure 6:**
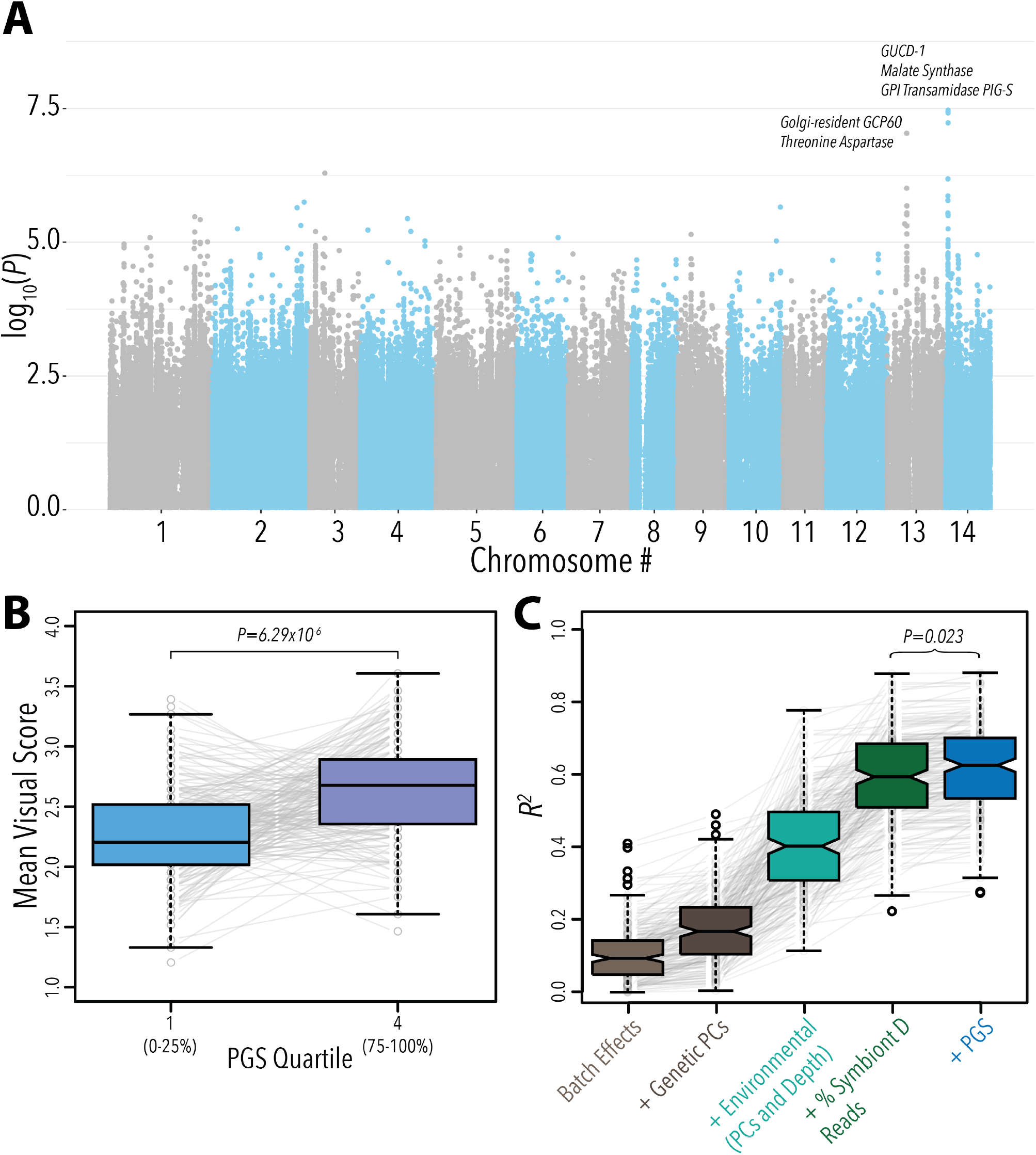
A genome wide association study and prediction accuracy for bleaching response. (**A**) A Manhattan plot of *P*-values obtained from testing for an association of each SNP with variation in visual score (see SOM). Shown are the names of genes in the two top peaks, on chromosome 13 and 14. (**B**) Individuals in the highest quartile of polygenic scores had a significantly higher visual score compared with individuals in the lowest quartile. Shown are the average visual scores for samples in the lowest and highest quartiles from 100 jackknife cross-validation partitions. Gray lines connecting individual points represent the difference in visual scores for each jackknife partition. (**C**) The increase in prediction accuracy (measured as *R*^2^) for linear models with different sets of predictors. Each boxplot shows the distribution of *R*^2^ obtained for 100 jackknife cross-validation partitions, with a random 15% of the total sample withheld as a test set in each. Batch effects include the collection date and the sequencing batch (see SOM). Each boxplot shows the *R*^2^ for sets of predictors in a linear model. Mann-Whitney *U* tests were performed to test for differences in the distribution of *R*^2^ between models.

### Predicting individual responses in natural populations

The GWAS results suggest that variation in bleaching is not due to common loci of large effect, such that our sample of 213 diploid individuals is under-powered to identify single loci that are genome-wide significant. Nonetheless, when combined, the estimated effects of variants across the genome may be predictive of phenotype^22^. We therefore constructed a polygenic score (PGS; also called breeding value^68^) using LD-based clumping (for a *P*-value threshold of 10^−5^; see SOM, Fig S30) and the effect sizes estimated in the GWAS^69^. Because we lacked a true out-of-sample validation set, we assessed the prediction accuracy of the PGS using a jackknife cross-validation procedure with an 85/15% training-test split in each partition; similar results were also obtained for 80/20% and 90/10% training-test splits (Fig S31).

A useful application of a PGS for bleaching in informing conservation strategies may be in identifying those individuals that are most tolerant, *i.e*., lie in the high end of the PGS distribution. We therefore examined if, using our cross-validation procedure, individuals in the top quartile of PGS had higher visual scores on average than those in the lowest quartile. The two quartiles differed by a mean visual score difference of more than 0.4 (Fig 6B; *P* = 6.29 × 10^−6^ by a one-tailed Wilcoxon signed rank test).

Given that both environmental and genetic factors contribute to variation in bleaching, we combined our PGS with other predictors, including environmental effects and the dominant symbiont species type. Using our cross-validation procedure, we assessed the improvement in prediction accuracy gained from including these predictors in a linear model (Figure 6C). Compared to a model including only batch effects and genetic PCs, we observed a large increase in prediction accuracy using four PCs for the environmental variables, as well as a substantial improvement when we included data obtained from our genomic sequencing, namely the proportion of *Durusdinium* reads out of all symbiont reads and the PGS (Figure 6C). Including the PGS alone provided a small but significant increase in prediction accuracy (*P* = 0.023; mean incremental *R*^2^ = 0.026) and the same was true in models using the top six or eight environmental PCs instead (which in total explain 99% of the variance in environmental variables, see SOM, Fig S32; *P* < 0.029). In total, the model explains an estimated ~ 62% of the variance in quantile-normalized visual score in our sample. As sample sizes increase, so will the prediction accuracy of the PGS, and hence of the model; moreover, it will become possible to test for interactions between the coral genome and environmental factors as well as symbiont type.

### Implications

Understanding how species will respond to increasing temperatures is key to ensuring adequate protection by conventional management approaches and to supporting novel interventions aimed at restoration or facilitating adaptation to prevent future losses. In this work, we focused on bleaching in the coral *A. millepora*, a trait of central ecological importance, demonstrating the feasibility of predicting inter-individual response from genomic data as well as environmental variables. Moving forward, a predictive model of bleaching can help distinguish those individuals more likely to be tolerant on reefs across different shelf positions, latitudes, and environmental conditions, informing current spatial protection strategies and predictions about reef futures^70,71^. The approach may also help to identify the most tolerant individuals to use for managed translocations and selective breeding for assisted evolution-based conservation strategies^72^. Alternatively, such predictions could be used to increase the abundance of such corals in local populations, rather than moving them through assisted gene flow, which carries risks and ethical concerns.

### Data Availability

The reference genome assembly is available for download at www.przeworskilab.com/data/. All paired-end reads are deposited on the SRA under accession numbers SAMN13447100-13447355 and under BioProject PRJNA593014.

## Supporting information

SOM

## Acknowledgments

We thank members of the Australian Institute of Marine Science’s coral bleaching group for assistance with sample collections; Richard Durbin, Arbel Harpak, Molly Schumer and the Andolfatto, Przeworski, and Sella labs for helpful discussions. We are particularly grateful to John Sorkness from the Austin Reef Club for providing a live *A. millepora* fragment as a source of high-quality DNA for genome sequencing. This project was funded by the Australian Institute of Marine Science, the Australian Government’s National Environmental Science Program, the Agouron Institute and by Columbia University. We acknowledge computing resources from Columbia University’s Shared Research Computing Facility project, which is supported by NIH Research Facility Improvement Grant 1G20RR030893-01, and associated funds from the New York State Empire State Development, Division of Science Technology and Innovation (NYSTAR) contract C090171. The coral collections were conducted under Great Barrier Reef Marine Park Authority collection permit G16/38488.1.

## References

[1] C. Bellard, C. Bertelsmeier, P. Leadley, W. Thuiller, and F. Courchamp, Impacts of climate change on the future of biodiversity, Ecology Letters 365–377 (2018), ISSN 1461-0248.

[2] C. Parmesan and G. Yohe, A globally coherent fingerprint of climate change impacts across natural systems, Nature 421, 37 (2003), ISSN 1476-4687.

[3] S. A. Henson, C. Beaulieu, T. Ilyina, J. G. John, M. Long, R. Séférian, J. Tjiputra, and J. L. Sarmiento, Rapid emergence of climate change in environmental drivers of marine ecosystems, Nature Communications 8, 14682 (2017), ISSN 2041-1723.

[4] M. Pacifici, W. B. Foden, P. Visconti, J. E. M. Watson, S. H. M. Butchart, K. M. Kovacs, B. R. Scheffers, D. G. Hole, T. G. Martin, H. R. Akçakaya, et al., Assessing species vulnerability to climate change, Nature Climate Change 5, 215 (2015), ISSN 1758-6798.

[5] A. A. Hoffmann and C. M. Sgrò, Climate change and evolutionary adaptation, Nature 470, 479 (2011), ISSN 1476-4687.

[6] I.-C. Chen, J. K. Hill, R. Ohlemüller, D. B. Roy, and C. D. Thomas, Rapid Range Shifts of Species Associated with High Levels of Climate Warming, Science 333, 1024 (2011), ISSN 0036-8075, 1095-9203.

[7] G.-R. Walther, E. Post, P. Convey, A. Menzel, C. Parmesan, T. J. C. Beebee, J.-M. Fromentin, O. Hoegh-Guldberg, and F. Bairlein, Ecological responses to recent climate change, Nature 416, 389 (2002), ISSN 1476-4687.

[8] J. Merilä and A. P. Hendry, Climate change, adaptation, and phenotypic plasticity: the problem and the evidence, Evolutionary Applications 7, 1 (2014), ISSN 1752-4571.

[9] Kelly Morgan, Adaptation to climate change through genetic accommodation and assimilation of plastic phenotypes, Philosophical Transactions of the Royal Society B: Biological Sciences 374, 20180176 (2019).

[10] D. J. Ayre and T. P. Hughes, Climate change, genotypic diversity and gene flow in reefbuilding corals, Ecology Letters 7, 273 (2004), ISSN 1461-0248.

[11] T. C. LaJeunesse, J. E. Parkinson, P. W. Gabrielson, H. J. Jeong, J. D. Reimer, C. R. Voolstra, and S. R. Santos, Systematic Revision of Symbiodiniaceae Highlights the Antiquity and Diversity of Coral Endosymbionts, Current biology: CB 28, 2570 (2018), ISSN 1879-0445.

[12] N. Traylor-Knowles, Heat stress compromises epithelial integrity in the coral, Acropora hyacinthus, Peer J 7, e6510 (2019), ISSN 2167-8359.

[13] T. P. Hughes, K. D. Anderson, S. R. Connolly, S. F. Heron, J. T. Kerry, J. M. Lough, A. H. Baird, J. K. Baum, M. L. Berumen, T. C. Bridge, et al., Spatial and temporal patterns of mass bleaching of corals in the Anthropocene, Science 359, 80 (2018), ISSN 0036-8075, 1095-9203.

[14] T. P. Hughes, J. T. Kerry, M. Alvarez Noriega, J. G. Alvarez Romero, K. D. Anderson, A. H. Baird, R. C. Babcock, M. Beger, D. R. Bellwood, R. Berkelmans, et al., Global warming and recurrent mass bleaching of corals, Nature 543, 373 (2017), ISSN 0028-0836.

[15] T. P. Hughes, J. T. Kerry, A. H. Baird, S. R. Connolly, A. Dietzel, C. M. Eakin, S. F. Heron, A. S. Hoey, M. O. Hoogenboom, G. Liu, et al., Global warming transforms coral reef assemblages, Nature 556, 492 (2018), ISSN 1476-4687.

[16] D. J. Ayre and T. P. Hughes, Genotypic diversity and gene flow in brooding and spawning corals along the Great Barrier Reef, Australia, Evolution; International Journal of Organic Evolution 54, 1590 (2000), ISSN 0014-3820.

[17] M. J. H. Van Oppen, L. M. Peplow, S. Kininmonth, and R. Berkelmans, Historical and contemporary factors shape the population genetic structure of the broadcast spawning coral, Acropora millepora, on the Great Barrier Reef, Molecular Ecology 20, 4899 (2011), ISSN 1365-294X.

[18] J. R. Guest, J. Low, K. Tun, B. Wilson, C. Ng, D. Raingeard, K. E. Ulstrup, J. T. I. Tanzil, P. A. Todd, T. C. Toh, et al., Coral community response to bleaching on a highly disturbed reef, Scientific Reports 6, srep20717 (2016), ISSN 2045-2322.

[19] G. B. Dixon, S. W. Davies, G. V. Aglyamova, E. Meyer, L. K. Bay, and M. V. Matz, Genomic determinants of coral heat tolerance across latitudes, Science 348, 1460 (2015), ISSN 0036-8075, 1095-9203.

[20] M. V. Matz, E. A. Treml, G. V. Aglyamova, and L. K. Bay, Potential and limits for rapid genetic adaptation to warming in a Great Barrier Reef coral, PLOS Genetics 14, e1007220 (2018), ISSN 1553-7404.

[21] N. R. Wray, K. E. Kemper, B. J. Hayes, M. E. Goddard, and P. M. Visscher, Complex Trait Prediction from Genome Data: Contrasting EBV in Livestock to PRS in Humans: Genomic Prediction, Genetics 211, 1131 (2019), ISSN 1943-2631.

[22] E. S. Buckler, J. B. Holland, P. J. Bradbury, C. B. Acharya, P. J. Brown, C. Browne, E. Ersoz, S. Flint-Garcia, A. Garcia, J. C. Glaubitz, et al., The Genetic Architecture of Maize Flowering Time, Science 325, 714 (2009), ISSN 0036-8075, 1095-9203.

[23] A. V. Khera, M. Chaffin, K. G. Aragam, M. E. Haas, C. Roselli, S. H. Choi, P. Natarajan, E. S. Lander, S. A. Lubitz, P. T. Ellinor, et al., Genome-wide polygenic scores for common diseases identify individuals with risk equivalent to monogenic mutations, Nature Genetics 50, 1219 (2018), ISSN 1061-4036, 1546-1718.

[24] C. Shinzato, E. Shoguchi, T. Kawashima, M. Hamada, K. Hisata, M. Tanaka, M. Fujie, M. Fujiwara, R. Koyanagi, T. Ikuta, et al., Using the Acropora digitifera genome to understand coral responses to environmental change, Nature 476, 320 (2011), ISSN 0028-0836.

[25] M. Helmkampf, M. R. Bellinger, S. M. Geib, S. B. Sim, and M. Takabayashi, Draft Genome of the Rice Coral Montipora capitata Obtained from Linked-Read Sequencing, Genome Biology and Evolution 11, 2045 (2019).

[26] Y. Mao, E. P. Economo, and N. Satoh, The Roles of Introgression and Climate Change in the Rise to Dominance of Acropora Corals, Current biology: CB 28, 3373 (2018), ISSN 1879-0445.

[27] H. Ying, D. C. Hayward, I. Cooke, W. Wang, A. Moya, K. R. Siemering, S. Sprungala, E. E. Ball, S. Forêt, and D. J. Miller, The Whole-Genome Sequence of the Coral Acropora millepora, Genome Biology and Evolution 11, 1374 (2019).

[28] C. R. Voolstra, Y. Li, Y. J. Liew, S. Baumgarten, D. Zoccola, J.-F. Flot, S. Tambutté, D. Allemand, and M. Aranda, Comparative analysis of the genomes of Stylophora pistillata and Acropora digitifera provides evidence for extensive differences between species of corals, Scientific Reports 7, 1 (2017), ISSN 2045-2322.

[29] R. Cunning, R. A. Bay, P. Gillette, A. C. Baker, and N. Traylor-Knowles, Comparative analysis of the Pocillopora damicornis genome highlights role of immune system in coral evolution, Scientific Reports 8, 16134 (2018), ISSN 2045-2322.

[30] U. E. Siebeck, N. J. Marshall, A. Klüter, and O. Hoegh-Guldberg, Monitoring coral bleaching using a colour reference card, Coral Reefs 25, 453 (2006), ISSN 0722-4028, 1432-0975.

[31] S. A. Matthews, C. Mellin, A. MacNeil, S. F. Heron, W. Skirving, M. Puotinen, M. J. Devlin, and M. Pratchett, High-resolution characterization of the abiotic environment and disturbance regimes on the Great Barrier Reef, 1985–2017, Ecology 100, e02574 (2019), ISSN 1939-9170.

[32] M. Nei and W. H. Li, Mathematical model for studying genetic variation in terms of restriction endonucleases, Proceedings of the National Academy of Sciences 76, 5269 (1979), ISSN 0027-8424, 1091-6490.

[33] H. Li and R. Durbin, Inference of human population history from individual whole-genome sequences, Nature 475, 493 (2011), ISSN 0028-0836, 1476-4687.

[34] Z. T. Richards, D. J. Miller, and C. C. Wallace, Molecular phylogenetics of geographically restricted Acropora species: implications for threatened species conservation, Molecular Phylogenetics and Evolution 69, 837 (2013), ISSN 1095-9513.

[35] S. Schiffels and R. Durbin, Inferring human population size and separation history from multiple genome sequences, Nature genetics 46, 919 (2014), ISSN 1061-4036.

[36] C. Prada, B. Hanna, A. F. Budd, C. M. Woodley, J. Schmutz, J. Grimwood, R. Iglesias-Prieto, J. M. Pandolfi, D. Levitan, K. G. Johnson, et al., Empty Niches after Extinctions Increase Population Sizes of Modern Corals, Current Biology 26, 3190 (2016), ISSN 0960-9822.

[37] D. Petkova, J. Novembre, and M. Stephens, Visualizing spatial population structure with estimated effective migration surfaces, Nature Genetics 48, 94 (2016), ISSN 1546-1718.

[38] P. Souter, B. L. Willis, L. K. Bay, M. J. Caley, A. Muirhead, and M. J. H. v. Oppen, Location and disturbance affect population genetic structure in four coral species of the genus Acropora on the Great Barrier Reef, Marine Ecology Progress Series 416, 35 (2010), ISSN 0171-8630, 1616-1599.

[39] H. Levene, Genetic Equilibrium When More Than One Ecological Niche is Available, The American Naturalist 87, 331 (1953), ISSN 0003-0147.

[40] S. Yeaman and S. P. Otto, Establishment and Maintenance of Adaptive Genetic Divergence Under Migration, Selection, and Drift, Evolution 65, 2123 (2011), ISSN 1558-5646.

[41] N. R. Garud, P. W. Messer, E. O. Buzbas, and D. A. Petrov, Recent Selective Sweeps in North American Drosophila melanogaster Show Signatures of Soft Sweeps, PLOS Genetics 11, e1005004 (2015), ISSN 1553-7404.

[42] H. Li and P. Ralph, Local PCA Shows How the Effect of Population Structure Differs Along the Genome, Genetics 211, 289 (2019), ISSN 0016-6731, 1943-2631.

[43] M. Ménade, G. Kozlov, J.-F. Trempe, H. Pande, S. Shenker, S. Wickremasinghe, X. Li, H. Hojjat, M.-J. Dicaire, B. Brais, et al., Structures of Ubl and Hsp90-like domains of sacsin provide insight into pathological mutations, Journal of Biological Chemistry jbc.RA118.003939 (2018), ISSN 0021-9258, 1083-351X.

[44] D. A. Parfitt, G. J. Michael, E. G. M. Vermeulen, N. V. Prodromou, T. R. Webb, J.-M. Gallo, M. E. Cheetham, W. S. Nicoll, G. L. Blatch, and J. P. Chapple, The ataxia protein sacsin is a functional co-chaperone that protects against polyglutamine-expanded ataxin-1, Human Molecular Genetics 18, 1556 (2009), ISSN 0964-6906.

[45] E. M. Hemond, S. T. Kaluziak, and S. V. Vollmer, The genetics of colony form and function in Caribbean Acropora corals, BMC Genomics 15 (2014), ISSN 1471-2164.

[46] A. B. Mayfield, Y.-J. Chen, C.-Y. Lu, and C.-S. Chen, The proteomic response of the reef coral Pocillopora acuta to experimentally elevated temperatures, PLOS ONE 13, e0192001 (2018), ISSN 1932-6203.

[47] R. Cunning and A. C. Baker, Not just who, but how many: the importance of partner abundance in reef coral symbioses, Frontiers in Microbiology 5 (2014), ISSN 1664-302X.

[48] A. B. Mayfield, Y.-B. Wang, C.-S. Chen, S.-H. Chen, and C.-Y. Lin, Dual-compartmental transcriptomic + proteomic analysis of a marine endosymbiosis exposed to environmental change, Molecular Ecology 25, 5944 (2016), ISSN 1365-294X.

[49] O. Delaneau, B. Howie, A. J. Cox, J.-F. Zagury, and J. Marchini, Haplotype estimation using sequencing reads, American Journal of Human Genetics 93, 687 (2013), ISSN 1537-6605.

[50] S. Picelli, K. Björklund, B. Reinius, S. Sagasser, G. Winberg, and R. Sandberg, Tn5 transposase and tagmentation procedures for massively scaled sequencing projects, Genome Research 24, 2033 (2014), ISSN 1088-9051.

[51] L. Huang, Y. Li, A. B. Singleton, J. A. Hardy, G. Abecasis, N. A. Rosenberg, and P. Scheet, Genotype-imputation accuracy across worldwide human populations, American Journal of Human Genetics 84, 235 (2009), ISSN 1537-6605.

[52] B. L. Browning, Y. Zhou, and S. R. Browning, A One-Penny Imputed Genome from Next-Generation Reference Panels, American Journal of Human Genetics 103, 338 (2018), ISSN 1537-6605.

[53] D. P. Manzello, M. V. Matz, I. C. Enochs, L. Valentino, R. D. Carlton, G. Kolodziej, X. Serrano, E. K. Towle, and M. Jankulak, Role of host genetics and heat-tolerant algal symbionts in sustaining populations of the endangered coral Orbicella faveolata in the Florida Keys with ocean warming, Global Change Biology 25, 1016 (2019), ISSN 1365-2486.

[54] P. M. Visscher, G. Hemani, A. A. E. Vinkhuyzen, G.-B. Chen, S. H. Lee, N. R. Wray, M. E. Goddard, and J. Yang, Statistical Power to Detect Genetic (Co)Variance of Complex Traits Using SNP Data in Unrelated Samples, PLOS Genetics 10, e1004269 (2014), ISSN 1553-7404.

[55] P. M. Visscher, S. E. Medland, M. A. R. Ferreira, K. I. Morley, G. Zhu, B. K. Cornes, G. W. Montgomery, and N. G. Martin, Assumption-Free Estimation of Heritability from Genome-Wide Identity-by-Descent Sharing between Full Siblings, PLoS Genetics 2 (2006), ISSN 1553-7390.

[56] J. E. Parkinson and I. B. Baums, The extended phenotypes of marine symbioses: ecological and evolutionary consequences of intraspecific genetic diversity in coral–algal associations, Frontiers in Microbiology 5 (2014), ISSN 1664-302X.

[57] T. F. Cooper, R. Berkelmans, K. E. Ulstrup, S. Weeks, B. Radford, A. M. Jones, J. Doyle, M. Canto, R. A. O’Leary, and M. J. H. v. Oppen, Environmental Factors Controlling the Distribution of Symbiodinium Harboured by the Coral Acropora millepora on the Great Barrier Reef, PLOS ONE 6, e25536 (2011), ISSN 1932-6203.

[58] M. Stat and R. D. Gates, Clade D Symbiodinium in Scleractinian Corals: A “Nugget” of Hope, a Selfish Opportunist, an Ominous Sign, or All of the Above? (2011), https://www.hindawi.com/journals/jmb/2011/730715/.

[59] M. K. Morikawa and S. R. Palumbi, Using naturally occurring climate resilient corals to construct bleaching-resistant nurseries, Proceedings of the National Academy of Sciences 116, 10586 (2019), ISSN 0027-8424, 1091-6490.

[60] A. Klueter, J. Trapani, F. I. Archer, S. E. McIlroy, and M. A. Coffroth, Comparative growth rates of cultured marine dinoflagellates in the genus Symbiodinium and the effects of temperature and light, PLOS ONE 12, e0187707 (2017), ISSN 1932-6203.

[61] X. Zhou and M. Stephens, Genome-wide efficient mixed-model analysis for association studies, Nature Genetics 44, 821 (2012), ISSN 1546-1718.

[62] X. Zhou, A unified framework for variance componenet estimation with summary statistics in genome-wide association studies, The Annals of Applied Statistics 11, 2027 (2017), ISSN 1932-6157.

[63] P. R. Muir, P. A. Marshall, A. Abdulla, and J. D. Aguirre, Species identity and depth predict bleaching severity in reef-building corals: shall the deep inherit the reef?, Proceedings of the Royal Society B: Biological Sciences 284, 20171551 (2017).

[64] B. Servin and M. Stephens, Imputation-Based Analysis of Association Studies: Candidate Regions and Quantitative Traits, PLOS Genetics 3, e114 (2007), ISSN 1553-7404.

[65] F. A. Kondrashov, E. V. Koonin, I. G. Morgunov, T. V. Finogenova, and M. N. Kondrashova, Evolution of glyoxylate cycle enzymes in Metazoa: evidence of multiple horizontal transfer events and pseudogene formation, Biology Direct 1, 31 (2006), ISSN 1745-6150.

[66] N. R. Polato, N. S. Altman, and I. B. Baums, Variation in the transcriptional response of threatened coral larvae to elevated temperatures, Molecular Ecology 22, 1366 (2013), ISSN 1365-294X.

[67] R. M. Wright, G. V. Aglyamova, E. Meyer, and M. V. Matz, Gene expression associated with white syndromes in a reef building coral, Acropora hyacinthus, BMC genomics 16, 371 (2015), ISSN 1471-2164.

[68] D. S. Falconer and T. F. C. Mackay, Introduction to Quantitative Genetics (Pearson, Harlow, 1996), 4th ed., ISBN 978-0-582-24302-6.

[69] S. Purcell, B. Neale, K. Todd-Brown, L. Thomas, M. A. R. Ferreira, D. Bender, J. Maller, P. Sklar, P. I. W. de Bakker, M. J. Daly, et al., PLINK: A Tool Set for WholeGenome Association and Population-Based Linkage Analyses, The American Journal of Human Genetics 81, 559 (2007), ISSN 0002-9297.

[70] I. B. Baums, A restoration genetics guide for coral reef conservation, Molecular Ecology 17, 2796 (2008), ISSN 1365-294X.

[71] J. E. Parkinson, A. C. Baker, I. B. Baums, S. W. Davies, A. G. Grottoli, S. A. Kitchen, M. V. Matz, M. W. Miller, A. A. Shantz, and C. D. Kenkel, Molecular tools for coral reef restoration: Beyond biomarker discovery, Conservation Letters n/a, e12687 (2019), ISSN 1755-263X.

[72] M. J. H. v. Oppen, J. K. Oliver, H. M. Putnam, and R. D. Gates, Building coral reef resilience through assisted evolution, Proceedings of the National Academy of Sciences 112, 2307 (2015), ISSN 0027-8424, 1091-6490.

