## Supplementary material for "Population genetics of the coral *Acropora millepora*: Towards a genomic predictor of bleaching": SOM

### Materials and Methods for *"Population genetics of the coral Acropora millepora: Towards a genomic predictor of bleaching"*

Zachary Fuller*, Veronique Mocellin, Luke Morris, Neal Cantin, Jihanne Shepherd, Luke Sarre, Julie Peng, Yi Liao, Joseph Pickrell, Peter Andolfatto, Mikhail Matz, Line Bay, Molly Przeworski

**8 Supplemental Figures** **35**

**9 Supplemental Tables** **67**

**10 References** **75**

### 1 Genome Assembly

#### 1.1 Overview

The *de novo* genome assembly of the coral *Acropora millepora* was constructed using a combination of PacBio reads and Illumina paired-end reads with 10X Chromium barcodes. High-molecular weight DNA was extracted from sample JS-1 (donated graciously by the Austin Reef Club) and sequencing was performed by the New York Genome Center (10X-HiSeqX) and Cold Spring Harbor Laboratory (PacBio Sequel). A general overview of the assembly process is shown in Figure S1.

#### 1.2 Initial PacBio Assembly

Two initial PacBio-only assemblies were constructed, one using the software Canu[1] and the other using Falcon[2]. The Falcon assembly was later scaffolded using 10X Chromium barcodes and is referred to as the "consensus" assembly. All genome assembly steps were performed on Columbia University's High Performance Computing (HPC) Habanero cluster. Below are the commands and software used for each stage of the assembly.

##### 1.2.1 Canu Assembly

**Preprocessing**
First, a .xml settings file was created containing information pointing to the raw PacBio reads. Then, raw .bam files were converted to a .fasta file of subreads:

```
$ /smrtlink5/smrtcmds/bin/dataset create --type SubreadSet \
       subreads.bam.fofn subreads.xml
$ /smrtlink5/smrtcmds/bin/bam2fasta -o JS1.subreads.raw.fasta subreads.xml
```

**Assembly**
The initial assembly parameters are as follows:

```
genomeSize=420m
useGrid=True
gnuplotTested=true
--gridOptions=-A mfplab
ovsMemory=120
ovsThreads=24
ovsMethod=parallel
ovbMemory=120
ovbThreads=8
batMemory=200
merylMemory=64g
```

The genome size estimate was based on *A. digitifera*[3]. Using the subreads and above parameters in the configuration file, Canu was run to construct an initial PacBio-only assembly:

```
$ /canu -p AM -pacbio-raw JS1.subreads.raw.fasta
```

### of Contigs: 9140
Assembly N50: 220897
Assembly Length (bp): 877872071

##### 1.2.2 Falcon Assembly

**Initial Falcon assembly** The first assembly from running Falcon was constructed using only PacBio sequences. The config file (*fc_run.cfg*) for Falcon was set as:

```
[General]
use_tmpdir = /rigel/mfplab/users/zlf2101/coral_genomes/AM/pacbio_reads/tmp_files
job_type = slurm
input_fofn = inputs.fofn
input_type = raw
length_cutoff = -1
length_cutoff_pr = 1000
genome_size = 420000000
seed_coverage = 30
job_queue = mfplab2
jobqueue = mfplab
```

```
sge_option_da = -A %(jobqueue)s -c 8 -t 6:00:00 --mem=64gb
sge_option_la = -A %(jobqueue)s -c 2 -t 6:00:00 --mem=32gb
sge_option_pda = -A %(jobqueue)s -c 8 -t 6:00:00 --mem=64gb
sge_option_pla = -A %(jobqueue)s -c 8 -t 6:00:00 --mem=32gb
sge_option_fc = -A %(jobqueue)s -c 24 -t 6:00:00 --mem=124gb
sge_option_cns = -A %(jobqueue)s -c 8 -t 6:00:00 --mem=64gb
pa_concurrent_jobs = 32
ovlp_concurrent_jobs = 32
pa_HPCdaligner_option =  -v -B24 -t16 -e.70 -l1000 -s1000
ovlp_HPCdaligner_option = -v -B24 -t32 -h70 -e.95 -l1000 -s500 -k25 -w5
pa_DBsplit_option = -x500 -s100
ovlp_DBsplit_option = -x500 -s100
falcon_sense_option = --output_multi --min_idt 0.70 \\
        --min_cov 4 --max_n_read 200 --n_core 12
overlap_filtering_setting = --max_diff 100 --max_cov 100 \\
        --min_cov 5 --bestn 10 --n_core 24
```

**scaff10x (Round 1)** The initial PacBio Falcon assembly was then scaffolded using single-molecule information provided from 10X Chromium barcoded linked reads. Two iterations of scaff10x (https://github.com/wtsi-hpag/Scaff10X) were run.

```
#Debarcode 1-reads:
INPUT=$1
BC_NAME="${INPUT/R1/RC1}"
OUT_NAME="${INPUT/R1/BC}"
NAME="${OUT_NAME/fastq/name}"
SCAFF_PATH="/rigel/mfplab/users/zlf2101/software/scaff10x/"
$SCAFF_PATH/scaff_BC-reads-1 $INPUT $BC_NAME $NAME > try.out
#Debarcode 2-reads:
INPUT=$1
BC_NAME="${INPUT/R2/RC2}"
OUT_NAME="${INPUT/R2/BC}"
NAME="${OUT_NAME/fastq/name}"
$SCAFF_PATH/scaff_BC-reads-2 $NAME $INPUT $BC_NAME > try.out
PACBIO_ASMBLY=$1
OUTPUT=$2
READ1_10X=$3
READ2_10X=$4
$SCAFF_PATH/scaff10x -nodes 24 -matrix 2000 -reads 10 -score 20 -pacbio 0 \\
        -edge 50000 -link 8 -block 10000 -align bwa \\
        $PACBIO_ASMBLY $READ1_10X $READ2_10X $OUTPUT > try.out
```

The output .fasta from the first iteration of scaff10x was then set as PACBIO_ASMBLY, and a second iteration was performed using the same parameters.
### of Scaffolds: 2293
Assembly N50: 603371
Assembly Length (bp): 647980563
Using the short insert reads with barcodes removed, the $k$-mer distribution was analyzed to estimate the genome size, heterozygosity (*i.e.,* nucleotide diversity $\pi$), and repeat content. Using a $k$-mer size of $k = 51$, this distribution was analyzed using the GenomeScope web server (http://qb.cshl.edu/genomescope/) (Figure S2). This optimal $k$-mer size was determined using KmerGenie (http://kmergenie.bx.psu.edu).

**ARCS/LINKS scaffolding** The resulting assembly after two rounds of running scaff10x was further scaffolded using the ARCS/LINKS[4,5] pipeline with the 10X Chromium barcodes and paired-end reads as input. Longranger software (https://support.10xgenomics.com) was first used to create an interleaved file of barcoded paired-end reads:

```
longranger-2.1.6/longranger-cs/2.1.6/bin/basic --id=JS-1 \\
       --fastqs=/10X_reads/raw_fastqs/ \\
       --sample=BGSK639-JS-1-CTAAACGG,BGSK639-JS-1-AACCGTAA, \\
                BGSK639-JS-1-GGTTTACT,BGSK639-JS-1-TCGGCGTC
gunzip -c Longranger barcoded.fastq.gz | \\
        perl -ne 'chomp;
        $ct++;
               $ct=1 if($ct>4);
                       if($ct==1){if(/(@\S+)\sBX:Z:(\S{16})/)
                       {$flag=1;$head=$1."_".$2;print "$head\n";}
               else{$flag=0;}}else{print "$\n" if($flag);}' > CHROMIUM_interleaved.fastq
```

The reads were then aligned to the assembly with `bwa-mem` and the contigs were renumbered to avoid formatting issues with ARCS software:

```
cat ../$SCAFF10x_1/pacbio.scaff10x.2.fasta |
        perl -ne 'chomp;
        if(/^>/){$ct++;print ">$ct\n";}
        else{print "$_\n";}' > pacbio.scaff10x.2-renamed.fasta
bwa index pacbio.scaff10x.2-renamed.fasta
bwa mem -t32 pacbio.scaff10x.2-renamed.fasta -p CHROMIUM_interleaved.fastq \\
        > bwa_aligned_chromium.sam
samtools view -Sb bwa_aligned_chromium.sam \\
        >bwa_aligned_chromium.bam
samtools sort -n bwa_aligned_chromium.bam -@ 16 -O BAM \\
        -o bwa_aligned_chromium_sorted.bam
```

A Graphviz Dot file (.gv) was then generated using ARCS:

```
DRAFT=$1
ALIGNS=$2
echo $DRAFT
echo $ALIGNS
$ARCS_PATH/arcs -f $DRAFT -a $ALIGNS -s 98 -c 5 -l 0 -d 0 -r 0.05 \\
        -e 30000 -m 20-10000 -v &> ARCSlog.txt
python $ARCS_PATH/arcs/Examples/makeTSVfile.py <original.gv> <tigpair_checkpoint.tsv> $DRAFT
```

LINKS (v 1.8.5) was then used to join nodes in the graph produced by ARCS:

```
./LINKS -f pacbio.scaff10x.2-renamed.fasta -s empty.fof \\
        -m -d 4000 -k 20 -e 0.1 -l 5 -a 0.9 \\
        -t 2 -o 0 -z 500 -b links_c5r0.05e30000-l5-a0.9 \\
        -r -p 0.001 -x 1
```

### of Scaffolds: 2171
Assembly N50: 645672
Assembly Length (bp): 647981783

**SSPACE-longread scaffolding** The output assembly after running ARCS/LINKS was then further scaffolded using PacBio longread sequences with SSPACE-LongRead[6].

```
SSPACE_PATH=/rigel/mfplab/users/zlf2101/software/SSPACE-LongRead_v1-1
CONTIGS=../arcs_scaffolds/links_c5r0.05e30000-l5-a0.9.scaffolds.fa
PACBIO=../pacbio_reads/JS1.subreads.raw.fasta
perl $SSPACE_PATH/SSPACE-LongRead.pl -c $CONTIGS -p $PACBIO \\
        -b SSPACE_scaffolds/SSPACE_results -t 32
```

A final two iterations of scaff10x were performed on the resulting assembly: # of Scaffolds: 1553
Assembly N50: 938379
Assembly Length (bp): 650267226

**PBJelly gapfilling** Up to this step, the initial PacBio assembly was scaffolded using linked reads with 10X Chromium barcodes and raw PacBio reads. Gaps in this assembly were then filled using PacBio subreads with PBJelly software[7]. The PBJelly pipeline is a set of five steps (setup, support, mapping, extraction, and output) that is controlled by a single configuration file:

```
<jellyProtocol>
    <reference>sspace_scaff10x.2.fasta.renamed.fasta</reference>
    <outputDir>/rigel/mfplab/users/zlf2101/coral_genomes/AM/pbjelly2</outputDir>
    <blasr>-minMatch 8 -minPctIdentity 70 -bestn 1 \\
           -nCandidates 20 -maxScore -500 -nproc 16 -noSplitSubreads</blasr>
    <input baseDir="/rigel/mfplab/users/zlf2101/coral_genomes/AM/pbjelly2">
        <job>JS1.subreads.fasta</job>
    </input>
</jellyProtocol>
```

To avoid excessive memory consumption during the extraction step, only PacBio reads with evidence for gap support were read into memory (as opposed to the entire 36G JS1.subreads.fasta file). Prior to the extraction step, the read names contained in the .gml file in support/ were written as a list to a separate file and *faSomeRecords* (`https://github.com/ENCODE-DCC/kentUtils`) was used to create filtered-JS1.subreads.fasta, which contained only those PacBio reads to be used in the remainder of the pipeline. The configuration file was updated prior to running extraction, to locate this filtered set of PacBio subreads.

### of Scaffolds: 1539
Assembly N50: 952761
Assembly Length (bp): 654818319

**Arrow error correction** Error correction using raw PacBio data was then performed using the Arrow algorithm available from PacBio (`https://github.com/PacificBiosciences/GenomicConsensus`). Arrow is a consensus calling model based on a more straightforward hidden Markov model than the conditional random field approach implemented in the legacy *quiver* algorithm[8]. Reads were first aligned to the output from PBJelly using pbalign:

```
./pbalign --tmpDir ./arrow_ec/tmpfiles $READS \\
        ../pbjelly2/jelly.scaff10x.2.fasta $OUTPUT
```

Arrow was then performed on the resulting .bam alignment and .fasta output:

```
smrtlink5/smrtcmds/bin/arrow pb.raw.mapped.jelly2.bam -j 24 \\
        -r ../pbjelly2/jelly.scaff10x.2.fasta \\
        -o variants.gff -o consensus.fasta -o consensus.fastq
```

### of Scaffolds: 1532
Assembly N50: 954497
Assembly Length (bp): 657785445

**Pilon error correction** Final error correction and polishing for the Falcon assembly was performed using paired-end reads. 10X Chromium barcodes were removed from the paired-end reads used in pilon[9]. The debarcoded reads were then mapped to the consensus.fasta assembly generated from Arrow with `bwa-mem`[10]:

```
bwa index ../arrow_ec/consensus.fasta
bwa mem -t32 $REF $READS1 $READS2 > bwa_aligned_chromium.sam
samtools view -Sb bwa_aligned_chromium.sam > bwa_aligned_chromium.bam
```

```
samtools sort -n bwa_aligned_chromium.bam -@ 16 \\
        -O BAM -o bwa_aligned_chromium_sorted.bam
```

To parallelize pilon, error correction was performed for each scaffold in the assembly separately. First, a list of contig names was created and used as a parameter in the SLURM array batch job. On the Columbia University HPC Habanero cluster, there is an array size maximum of 1000, so two list files were created:

```
cat ../arrow_ec/consensus.fasta|grep ">"| \\
        tr -d ">"|awk '{printf("%s %s\n", NR, $0)}' > numbered_contigs.txt
split -l 1000 numbered_contigs.txt
mv xaa contigs.1.txt
mv xab contigs.2.txt
```

pilon was then performed on each scaffold in the assembly :

```
java -Xmx24G -jar ../../../software/pilon-1.22.jar \\
        --bam bwa_aligned_chromium_sorted.bam \\
        --genome ../arrow_ec/consensus.fasta \\
        --targets $CONTIG --threads 4 \\
        --output ${CONTIG}.corrected
```

#### 1.3 Mapping Larval Reads, Removing Symbiont Sequence & Final Assembly

##### 1.3.1 Mapping of Larval Reads

The symbiont can reach densities of up to 1 million cells per square centimeter of tissue in the adult polyps [11] from which the high molecular weight DNA of JS-1 sequenced for the assembly was extracted. As a result, we expect up to approximately 10% of the assembled contigs from JS-1 to be *Symbiodinium spp.* sequences. To remove symbiont contigs and sequences of other contaminants from the adult sample, we used an approach of sequencing larvae, in which symbiont densities are expected to be negligible (because they have not yet acquired them from the environment *i.e.,* they are aposymbiotic). Two pools of > 50 larval samples (lables: ”e” - Orpheus Island, ”g” - Princess Charlotte Bay) each were sequenced on an Illumina HiSeq platform, at 2x150bp PE configuration. Approximately 70Gb of raw sequence data was produced from each sample. First, raw sequencing reads were trimmed and quality control performed:

```
$SEQTK_PATH/seqtk trimfq $READ1 > $TRIM1
$SEQTK_PATH/seqtk trimfq $READ2 > $TRIM2
java -jar $TRIM_PATH/trimmomatic-0.38.jar PE -phred33 $TRIM1 $TRIM2 $TRIM1.a1.fq \\
        $TRIM1.a1.U.fq $TRIM2.a2.fq $TRIM2.a2.U.fq \\
        ILLUMINACLIP:polyA.fa:2:3:3 MINLEN:36 SLIDINGWINDOW:4:25
java -jar $TRIM_PATH/trimmomatic-0.38.jar PE -phred33 $TRIM1.a1.fq $TRIM2.a2.fq \\
        $TRIM1.b1.fq $TRIM1.b1.U.fq $TRIM2.b2.fq \\
        $TRIM2.b2.U.fq ILLUMINACLIP:adapter.fa:2:3:3 MINLEN:36 SLIDINGWINDOW:4:25
```

Reads were then mapped to *consensus.fa* and *canu.fa* using `bwa-mem`. Each assembly was broken at runs of more than 10 consecutive ”N”s to mitigate the effects of potential mis-scaffolded sequences:

```
bwa mem -t32 $REF $READ1 $READ2 > $PREFIX.bwa.sam
samtools view -Sb $PREFIX.bwa.sam > $PREFIX.bwa.sam.bam
samtools sort $PREFIX.bwa.sam.bam -@ 16 -O BAM -o $PREFIX.bwa.sam.bam.sorted.bam
```

##### 1.3.2 Endosymbiont Removal

We undertook an unsupervised clustering approach to remove assembled contigs that showed evidence of being derived from endosymbiont genomic material and other external contaminants. To create features for the unsupervised clustering, we first quantified the sequence composition by calculating frequencies for 1,2,3, and 4-mers in each contig (in both forward and reverse-compliment) using a custom Python program *kmerCount.py* (`https://github.com/zfuller5280/CoralGenomes`). Next, using the .bam alignments of the larval read mappings, we generated coverage histograms for each contig, as well as the proportion of each with 0-coverage:

```
bedtools genomecov -ibam $INPUTBAM > $SAMPLE.hist
```

For each contig, the coverage was summarized in bins of size depth=10 from 1x-1000x. The coverage histograms were included as features, along with the 1,2,3, and 4-*mer* frequencies, proportion of 0-coverage, and %GC content for the non-linear dimensionality reduction method t-distributed stochastic neighbor embedding (t-SNE)[12] (Figure S3). Cluster assignment was then performed from this dimensionality reduction with Hierarchical Density-Based Spatial Clustering of Applications with Noise (HBDSCAN). As a smaller number of assembled contigs derived from the symbiont are expected in the total assembly, the cluster with fewer contigs was then manually interrogated using *blastn* to confirm that the sequences were derived from endosymbiont genomic material if they significantly aligned to available *Symbiodinium spp.* draft genomes (availale from `http://refuge2020.reefgenomics.org`). In addition, *blastn* was performed for all sequences from the cluster with the largest number of contigs (assumed to be *A. millepora*); none matched any available *Symbiodinium spp.* genomes. Contigs belonging to the cluster with no evidence for being derived from coral and with evidence for being derived from endosymbiont or other external contaminants were then removed from each assembly:

```
#R
//For Canu
tsne<-Rtsne(merged_kmers,dims=2,perplexity=500,verbose=T,max_iter=2000)
cl <- hdbscan(tsne$Y, minPts = 50)
write.table(rownames(contig_features[cl$cluster==1,]),"
        canu_symbiont.contigs.txt",sep="\n",quote=F,row.names = F, col.names = F)
//For consensus
tsne<-Rtsne(merged_kmers,dims=2,perplexity=100,verbose=T,max_iter=2000)
cl <- hdbscan(tsne$Y, minPts = 50)
write.table(rownames(contig_features[cl$cluster==1,]),"
        cons_symbiont.contigs.txt",sep="\n",quote=F,row.names = F, col.names = F)
#/bin/bash
./faSomeRecords -exclude $INPUTRAWCONTIGS $NAMESLIST $OUTPUT
$ cat symbiont_contigs.txt |wc -l
>>   168
$ cat canu_symbiont.contigs.txt |wc -l
>>   891
```

##### 1.3.3 Merged Assembly & Polishing

The *quickmerge* program was used to merge the filtered *canu.fa* and *consensus.fa* assemblies:

```
srun -A mfplab /software/quickmerge/merge_wrapper.py \\
        filtered_consensus.fa filtered_canu.fa
```

*finishingTool* was then run using the PacBio subreads to polish the merged assembly generated from *quickmerge*:

```
//Get rid of special characters in fasta headers
perl -pe 's/>[^\$]*$/">Seg" . ++$n ."\n"/ge' \\
        ../filtered_quickmerge/quickmerge.out.fa > contigs.fasta
perl -pe 's/>[^\$]*$/">Seg" . ++$n ."\n"/ge' \\
        ../../AM/pacbio_reads/JS1.subreads.fasta > reads.fasta
python finishingTool/finisherSC.py -f True -l True -par 20 ../destinedFolder $MUMMER_PATH
```

A final round of *arrow* error correction and *pilon* polishing was performed on the resulting assembly.
sum = 682459408, n = 1194, ave = 571574.04, largest = 7711905
N50 = 1089919, n = 189

N60 = 884050, n = 258
N70 = 676857, n = 346
N80 = 504880, n = 463
N90 = 291973, n = 638
N100 = 2417, n = 1194
N_count = 10668
Gaps = 120

##### 1.3.4 HaploMerger2

Using the output of the raw assembly, the software HaploMerger2[13] was used to merge contigs/scaffolds that were assembled as individual diploid haplotypes into a single haploid genome assembly. First, as required by HaploMerger2, the assembly *Amil.asm.fa* was compressed using gzip and copied into the project_example1/ directory of the HaploMerger2 installation path. In the .sh files for each step of the HaploMerger2 pipeline, the parameter setting the number of processors was manually changed to 12.

```
sh ./hm.batchB1.initiation_and_all_lastz Amil
sh ./hm.batchB2.chainNet_and_netToMaf Amil
sh ./hm.batchB3.haplomerger Amil
sh ./hm.batchB4.refine_unpaired_sequences Amil
sh ./hm.batchB5.merge_paired_and_unpaired_sequences Amil
```

The final raw assembly statistics are:
sum = 525541835, n = 1065, ave = 493466.51, largest = 7982607
N50 = 1236380, n = 118
N60 = 1000970, n = 165
N70 = 755178, n = 225
N80 = 530310, n = 308
N90 = 285480, n = 439
N100 = 798, n = 1065
N_count = 9837
Gaps = 109

##### 1.3.5 Final Masking

To conservatively mask regions that may be the result of chimeric misassemblies of endosymbiont (or other external contaminant) sequences, we first located intervals with 0-coverage in both pooled larvae samples.

```
$/bedtools genomecov -bga -i g_ref.bwa.sam.sorted.bam|grep -w 0$ > g.zeroCov
$/bedtools genomecov -bga -i g_ref.bwa.sam.sorted.bam|grep -w 0$ > g.zeroCov
```

The mean size of such intervals was 72.38 bp in sample "e" and 72.45 bp in sample "g". Using these intervals, we then defined regions where neither pooled sample had any reads mapping (*i.e.,* both had 0-coverage).

```
#R
e_zero<-read.table("e.zeroCov")
g_zero<-read.table("g.zeroCov")
outersect <- function(x, y, ...) {
  big.vec <- c(x, y, ...)
  duplicates <- big.vec[duplicated(big.vec)]
  setdiff(big.vec, unique(duplicates))
}

library(GenomicRanges)
e_zero<-subset(e_zero, !(e_zero$V1 %in% outersect(e_zero$V1,g_zero$V1)))
g_zero<-subset(g_zero, !(g_zero$V1 %in% outersect(e_zero$V1,g_zero$V1)))
gr0 = with(e_zero, GRanges(V1, IRanges(start=V2, end=V3)))
```

```
gr1 = with(g_zero, GRanges(V1, IRanges(start=V2, end=V3)))
queryHits = findOverlaps(gr0, gr1)
subjectHits = findOverlaps(gr1, gr0)
hits = findOverlaps(gr0, gr1)
gr2<-pintersect(gr0[queryHits(hits)],gr1[subjectHits(hits)])
in_widths<-(width(ranges(gr2)))
df <- data.frame(seqnames=seqnames(gr2),
                 starts=start(gr2),
                 ends=end(gr2),
                 names=c(rep(".", length(gr2))),
                 scores=c(rep(".", length(gr2))),
                 strands=strand(gr2))
write.table(df, file="zero_cov.bed", quote=F, sep="\t", row.names=F, col.names=F)
```

In total, 25706057 bp of the raw assembly (4.89%) was not covered by reads in either pooled larval sample, consistent with rough *a priori* expectation. We therefore masked these intervals, converting the identity of nucleotides contained within the regions to 'N'.

```
bedtools maskfasta -fi Amil.asm.fa -fo Amil.coverageMasked.fa -bed zero_cov.bed
```

Total assembly N_count = 25715894

##### 1.3.6 Removal of Redundant Contigs

Preliminary analyses of the *v1.01* assembly indicated that even after a first run of HaploMerger2, misassembled haplotypes (*i.e.,* haplotigs) were likely still present. For example, in approximately 10,200 gene pairs (see the next section for annotation steps), the fraction of synonymous differences *Ks* was lower than the mean diversity level genome-wide. To test for further evidence of haplotigs present in the assembly, an all-vs-all alignment was performed:

***(1) Evaluating for potential redundant contigs:***

```
makeblastdb -in all.maker.transcripts.fasta -dbtype nucl -parse_seqids
blastn -query all.maker.transcripts.fasta -db all.maker.transcripts.fasta \\
          -evalue 0.000001 -out AmiGenSel.blast -outfmt 6 -num_threads 68
perl Gene_cluster_blastnfmt6.pl AmiGenSel.blast
awk '{if ($1 == $2){} else { print $0}}' AmiGenSel.blast.out > AmiGenSel.blast.out.out
perl GetUniqpairs.pl AmiGenSel.blast.out.out > AmiGenSel.blast.out.out.out
#The above commands will produce a total of 13,000 potential paralog pairs
#The Ks value for each pair was caluculated by KaKs_calculator:
#https://code.google.com/archive/p/kaks-calculator/
#All custom perl scripts can be found at:
#https://github.com/yiliao1022/Acropora_millepora_genomics
```

***(2) all-vs-all alignment with Lastz:***

```
lastz Amil.fasta[multi] Amil.fasta[multi] --strand=both --ambiguous=n \\
           --inner=2000 --gappedthresh=6000 --gapped --identity=90 --format=axt > all.axt
axtChain --linearGap=medium all.axt Amil.fasta Amil.fasta all.axt.chain
chainMergeSort all.axt.chain > all.chain
cat all.chain | grep chain | head -n 10000 > chain.out
perl GetPotentialAlle.pl chain.out > temp.txt
```

On the basis of the all-vs-all alignment, 54 contigs accounting for approximately 6 Mb of sequence were removed from the assembly before proceeding to chromosome scaffolding.

##### 1.3.7 Chromosome Scaffolding

*A. millepora*, like the majority of corals in the *Acropora* genus, is diploid with *2n*=28 chromosomes[14]. We used two previously published linkage maps inferred from SNP/RAD-seq markers to linearly arrange the scaffolds from the initial draft assembly into 14 linkage groups. The Wang *et al.* (2009)[15] map had 429 markers and examined 80 meioses, and the Dixon *et al.* (2015)[16] map relied on 1448 markers and 329 meioses. We first used the software Chromonomer (`http://catchenlab.life.illinois.edu/chromonomer/`) to integrate the genome assembly with these previously published genetic maps. Chromonomer requires three input files in addition to the .fasta file containing the assembly: i) a .tsv containing the genetic location of markers, ii) a SAM/BAM file containing the alignments of the genetic markers to the assembly, and iii) an AGP file describing the location of contigs within the assembly. The .tsv files are available from the original publications and were concatenated together.

**Genetic marker mapping:** The .fasta file containing sequences of the genetic markers was mapped to the assembly to produce a SAM file.

```
bwa mem Amil.asm.fa all_links_marks.fasta > all_links.sam
```

**AGP file conversion:** The Perl script *fa2agp.pl* was used (https://www.hmpdacc.org/hmp/doc/fasta2apg.pl)

```
perl ~/fa2agp.pl -i Amil_ref.fa -o . -n Amil_ref
```

**Chromonomer run:** Parameters were set as default and the assembly .fasta file was used as an additional input, resulting in a .fasta output of chromosome-level scaffolds.

```
~/chromonomer -p allMap.markers.tsv -o 20180919 -s all_links.sam \\
        -a Amil_ref.agp --data_version 20180919
```

For a final orientation of the Chromonomer arranged scaffolds, ALLMAPS[17] was used. For ALLMAPS, the two linkage map marker sets were aligned separately to the assembly using `bwa-mem` and converted to a .tsv BED-style file containing the locations of alignments.

```
bwa mem Amil.chromonomer.chrs.fa dixon_markers.fasta|\\
            samtools view -b -|bedtools bamtobed -i stdin > dixon_map.bed
bwa mem Amil.chromonomer.chrs.fa wang_markers.fasta|\\
            samtools view -b -|bedtools bamtobed -i stdin > wang_map.bed
```

Using a script included in the ALLMAPS installation, the BED-style files were then merged. The merged BED file was then used as an input to ALLMAPS.

```
python -m jcvi.assembly.allmaps mergebed meyer_map.bed dixon_map.bed
python -m jcvi.assembly.allmaps path out.bed Amil.chromonomer.chrs.fa
```

Gaps of size 50bp were inserted in the chromosome scaffolds between contigs. The final, chromosome-level scaffolded assembly statistics are:
sum = 475381253, n = 854, ave = 556652.52, largest = 39361238
N50 = 19840543, n = 9
N60 = 19500522, n = 11
N70 = 16847141, n = 14
N80 = 904615, n = 45
N90 = 296754, n = 142
N100 = 798, n = 854
N_count = 37012
Gaps = 380

#### 1.4 Gene Annotation

##### 1.4.1 Running MAKER

Protein coding gene models of the *Acropora millepora* genome were annotated using the software MAKER-P v.2.31.10[18] on the TACC (Texas Advanced Coputing Center) cluster stampede2. An in-depth description of running MAKER for genome annotation is available from `https://gist.github.com/darencard/`. The following transcript and protein data were used for input to MAKER and set in the configuration files.

Transcript data used: We used previously published RNA-seq data from *A. millepora*, including NCBI accessions from Moya *et al.* (2012)[19] and Bhattacharya et al . (2016)[20]. Additionally, we used transcriptome data from *A. tenuis* (available at `https://matzlab.weebly.com/`) as alternative evidence to support the predicted gene models.

Protein data used: We used the published peptide sequences available from the *A. digitifera* reference genome assembly[3].

A first round of MAKER was run to predict gene models directly from transcript and protein evidence.

```
$/Software/maker/bin/maker -base Rnd1 maker_opts.ctl maker_bopts.ctl maker_exe.ctl
```

This initial run generated 34,478 putative gene models.

##### 1.4.2 Round 2 - SNAP and AUGUSTUS

Training parameters for *ab initio* gene-predictors (SNAP and AUGUSTUS) were set according to the instructions available at: `https://pods.iplantcollaborative.org/wiki/display/TUT/Training+ab+initio+Gene+Predictors+for+MAKER+genome+annotation`

AUGUSTUS was trained using Benchmarking Universal Single-Copy Orthologs (BUSCO v3)[21].

```
##Extracting the GFF files for EST,protein,repeat mapping result from the first MAKER round:
$Software/maker/bin/gff3_merge \\
      -n -s -d all_rnd1_master_datastore_index.log > all_rnd1.all.maker.noseq.gff
awk '{ if ($2 == "est2genome") print $0 }' \\
      all_rnd1.all.maker.noseq.gff > all_rnd1.all.maker.est2genome.gff
awk '{ if ($2 == "protein2genome") print $0 }'\\
      all_rnd1.all.maker.noseq.gff > all_rnd1.all.maker.protein2genome.gff
awk '{ if ($2 ~ "repeat") print $0 }'\\
      all_rnd1.all.maker.noseq.gff > all_rnd1.all.maker.repeats.gff
```

This run produced 30,990 gene models. Afterwards, another round of training for SNAP/AUGUSTUS and MAKER was performed using the GFF files generated from AUGUSTUS as input, yielding 29,683 predicted gene models. From this set, 66 genes with exons mapping across 1 or more scaffolds were removed from the annotation, resulting in 29,617 predicted gene models. For the chromosome-level assembly, we conducted two additional rounds of MAKER using the same settings listed above, without new parameter training steps (*i.e.,* we used the parameters for SNAP and AUGUSTUS from previous rounds). In the end, the MAKER pipeline predicted putative 28,186 gene models for the chromosome-scale assembly.

##### 1.4.3 BUSCO Analysis

We assessed annotation completeness using BUSCO v3. First, a BUSCO analysis was performed using all core eukaryotic transcripts.
Results:
C:87.7%[S:81.8%,D:5.9%],F:8.3%,M:4.0%,n:303
266 Complete BUSCOs (C)
248 Complete and single-copy BUSCOs (S)

18 Complete and duplicated BUSCOs (D)
25 Fragmented BUSCOs (F)
12 Missing BUSCOs (M)
303 Total BUSCO groups searched

An additional BUSCO analysis was performed using all core metazoan transcripts.
Results:
C:86 %[S:79.7%,D:6.3%],F:5.8%,M:8.2%,n:978
841 Complete BUSCOs (C)
779 Complete and single-copy BUSCOs (S)
62 Complete and duplicated BUSCOs (D)
57 Fragmented BUSCOs (F)
80 Missing BUSCOs (M)
978 Total BUSCO groups searched
Of the 978 near-universal single-copy orthologs (metazoan set) tested, 841 were completely present and 57 were fragmented, accounting for 91.8% of the total BUSCO group searched. For the eukaryote ortholog set, 96% of the core genes were present in the final annotation.

##### 1.4.4 Functional Annotation

Functional annotation of the predicted genes was performed using the Trinotate pipeline (`https://trinotate.github.io/`) and eggNOG-mapper (`https://github.com/jhcepas/eggnog-mapper/wiki`). The Trinotate pipeline leverages the results from different functional annotation strategies, including homology search (BLAST+/SwissProt), protein domain detection (HMMER/PFAM), protein signal peptide and transmembrane domain prediction (signalP/tmHMM), and published annotation database search (eggNOG/GO/Kegg).

```
$ software/Trinotate/admin/Build_Trinotate_Boilerplate_SQLite_db.pl \\
      Trinotate
makeblastdb -in uniprot_sprot.pep -dbtype prot
gunzip Pfam-A.hmm.gz
$ software/hmmer-3.1b2-linux-intel-x86_64/binaries/hmmpress Pfam-A.hmm
echo "blastx -query Amillepora.transcripts.fasta -db uniprot_sprot.pep \\
      -num_threads 64 -max_target_seqs 3 -outfmt 6 > blastx.outfmt6" > blast
echo "blastp -query Amillepora.proteins.fasta -db uniprot_sprot.pep \\
      -num_threads 64 -max_target_seqs 3 -outfmt 6 \\
      > blastp.outfmt6" >> blast \\
       launcher_creator.py -n blast -j blast -a tagmap -N 2 \\
      -w 1 -t 12:00:00 -q normal -e
sbatch blast.slurm

echo "/software/tmhmm-2.0c/bin/tmhmm \\
      --short Amillepora.proteins.fasta > tmhmm.out" >> Run
echo "/software/signalp-4.1/signalp \\
      -f short -n signalp.out Amippepora.proteins.fasta" >> Run
echo "/software/hmmer-3.1b2-linux-intel-x86_64/binaries/hmmscan \\
      --cpu 64 --domtblout TrinotatePFAM.out Pfam-A.hmm Amillepora.proteins.fasta > pfam.log" >> Run
echo "/software/Trinotate/util/rnammer_support/RnammerTranscriptome.pl \\
      --transcriptome Amillepora.transcripts.fasta \\
      --path_to_rnammer /software/rnammer-1.2/rnammer" >> Run
launcher_creator.py -n Run -j Run -a tagmap -t 12:00:00 -N 4 -w 1 -q normal
sbatch Run.slurm

/software/Trinotate/Trinotate Trinotate.sqlite init \\
      --gene_trans_map Trinity.fasta.gene_trans_map \\
      --transcript_fasta Mcavernosa.transcripts.fa \\
```

```
      --transdecoder_pep Mcavernosa.proteins.fasta.pep
/software/Trinotate/Trinotate Trinotate.sqlite LOAD_swissprot_blastx \\
       blastx.outfmt6
/software/Trinotate/Trinotate Trinotate.sqlite LOAD_swissprot_blastp \\
       blastp.outfmt6
/software/Trinotate/Trinotate Trinotate.sqlite LOAD_pfam \\
       TrinotatePFAM.out
/software/Trinotate/Trinotate Trinotate.sqlite LOAD_tmhmm \\
       tmhmm.out
/software/Trinotate/Trinotate Trinotate.sqlite LOAD_signalp \\
       signalp.out
/software/Trinotate/Trinotate Trinotate.sqlite report \\
      > trinotate_annotation_report.xls
```

Trinotate retrieved 15,191 (54% of the total predicted set), 13,193 (47%), 12,490 (44%) GO, KEGG, and eggNog annotations, respectively.

For a final functional annotation step using orthology assignment, EggNog-mapper (`ttps://github.com/jhcepas/eggnog-mapper/`) was run on the full set of predicted gene models. This tool uses precomputed orthology groups and phylogenies available from the eggNOG database and has been demonstrated to have lower false positive rates compared to other homology-based approaches, such as BLAST and InterProScan[22].

```
python /software/eggnog-mapper/emapper.py \\
            -i ../Amil.proteins.fasta \\
            --output Amil_euk -d euk --usemem --cpu 60
```

eggNOG-mapper retrieved 9,182 (32.5% of the total predicted set), 8,536 (30%) and 15,717 (56%) GO, KEGG, and eggNog annotations, respectively.

##### 1.5 Comparison to the Ying *et al.* Genome Assembly

During the writing of this manuscript and after the initial release of our draft genome (August 1, 2018; `www.przeworskilab.com/data`), we became aware of an *A. millepora* assembly published by Ying *et al.* (2019)[23]. This assembly was generated from paired-end and mate-pair reads —DNA was extracted from sperm isolated from a single colony collected at Magnetic Island, Queensland. Descriptive statistics comparing our genome assembly and annotation with the Ying *at al.* (2019) release are given in Table S1. Notably, our assembly has greater contiguity, as indicated for example by comparing the scaffold N50 (19.8 Mb vs. 494 Kb). Diagrams showing the alignment of scaffolds from the Ying *at al.* assembly to the chromosome-scale scaffolds presented here are shown in Figure S4 and S5.

### 2 Samples

##### 2.1 Collection

Coral samples were collected from 12 reefs in the central region of the Great Barrier Reef at the height of mass bleaching, between 24 March and 1st April 2017 (Table S2). Samples were collected from up to 25 haphazardly selected colonies of *Acropora millepora* by SCUBA divers during one to two dives per site. For each coral colony, the bleaching score was estimated visually using the Coral Colour Reference Card[24] to the nearest colour including half increments and the depth and time of collection were also recorded. One to three single branches were collected per coral, in accordance with Great Barrier Reef Marine Park Permit G16/38488.1. During the dives, coral samples were sent to the surface every 5 minutes and immediately fixed in Liquid Nitrogen for further analysis.

##### 2.2 DNA Extraction

A small subsample was cut from the base of each frozen coral sample using bonecutters and stored in 100% ethanol at room temperature until DNA extraction. DNA extraction was carried out using the E.Z.N.A.

Tissue DNA Kit (Omega Bio-tek, USA) following the manufacturer's tissue protocol with slight modifications. Approximately 20 mg of coral surface tissue/skeleton was cut from each sample using a scalpel, weighed and combined with TL buffer, OB Protease and 0.5 mm sterile glass beads in a 1.5 ml centrifuge tube. The tubes containing the coral tissue samples were then homogenized three times at a speed of 4 m.s-1 for 30 s (FastPrep-24 5G, MP Biomedicals, USA) and incubated overnight in a microarray oven (30 rpm, ~16 hrs, 55 °C; Model 777, SciGene, USA). The remainder of the DNA extraction was carried out using the standard protocol and DNA was eluted in 50 $\mu$l. DNA quantity and quality were assessed using a spectrophotometer (NanoDrop 2000, Thermo Scientific, USA) and gel electrophoresis. DNA was stored at -20 °C before shipping to USA in unskirted PCR plates.

#### 2.3 Tissue Blasting

Coral samples were removed from the -80 °C freezer and placed on ice. Care was taken to work in the dark and to keep the samples cold during tissue blasting. Coral holobiont tissues were removed from their skeletons with high-pressure air into approximately 10 ml of pre-chilled 0.04 $\mu$M ultra-filtered seawater (FSW). Coral blastate was transferred to a 15 ml centrifuge tube and the volume recorded. The coral blastate was then homogenized for approximately 30 s using a post-mounted laboratory homogenizer (Bio-Gen PRO200, PRO Scientific, USA). A 1 ml aliquot of the coral homogenate was then taken, centrifuged (1,500 × g, 3 min, 4 °C), the supernatant discarded, and the resulting algal symbiont pellet was stored at -80°C until chlorophyll analysis. The remaining coral homogenate was centrifuged (1,500 × g, 3 min, 4 °C), and 500 $\mu$l of the resulting coral host supernatant was aliquoted in triplicate into deep-well plates and stored at -80 °C until coral host protein analysis.

#### 2.4 Chlorophyll Analysis

Algal symbiont samples were defrosted on ice; care was taken to keep these samples cold and dark during processing. After defrosting, any remaining supernatant was removed and 700 $\mu$l of pre-chilled 95% ethanol was added to each symbiont pellet. The symbiont pellet was then resuspended using a vortex and a pipette when required. The samples were then sonicated for 3 min in a custom-built ice bath attached to an ultrasonic generator (Sonic Power MU-600, Mirae Ultrasonic, South Korea), resuspended again and incubated on ice for 20 min. The samples were finally centrifuged (10,000 × g, 5 min, 4 °C) and 200 $\mu$l of the resulting supernatant was aliquoted in triplicate into clear 96-well plates. The absorbance was read on a microplate reader (Synergy H4, BioTek Instruments, USA) at 665, 649 and 632 nm at 25 °C. The values were then corrected against 95% ethanol blanks and chlorophylls a, c and total chlorophyll were calculated using the following equations[25]:

$$
\begin{aligned}
Chl_a &= (-0.9394 \times Abs_{632nm}) + (-4.2774 \times Abs_{649nm}) + (13.3914 \times Abs_{665nm}) \\
Chl_c &= (28.5073 \times Abs_{632nm}) + (-9.9940 \times Abs_{649nm}) + (-1.9749 \times Abs_{665nm}) \\
Chl_{total} &= (23.4742 \times Abs_{632nm}) + (11.4096 \times Abs_{649nm}) + (4.0735 \times Abs_{665nm})
\end{aligned}
$$

#### 2.5 Protein Analysis

Coral host samples were defrosted at room temperature and care was taken to keep the samples dark during processing. NaOH was added to the samples to reach a final concentration of 0.5 M and mixed using a pipette. The samples were sonicated for 5 min in a custom-built water bath attached to an ultrasonic generator (Sonic Power MU-600, Mirae Ultrasonic, South Korea) and incubated in a laboratory oven (1 hr, 90 °C) before centrifugation (1,500 x g, 10 min, 25 °C). Total coral host protein was then measured using the DC Protein Assay (Bio-Rad Laboratories, USA) following the manufacturer's microplate protocol and measuring absorbance on a microplate reader (Synergy H4, BioTek Instruments, USA). The measurements of coral host protein were used to normalize algal symbiont chlorophyll measurements and used as an approximate symbiont-to-host ratio or photosynthetic capacity of the symbiosis in order to characterize coral bleaching[26].

### 3 High Coverage Alignment & Variant Calling

#### 3.1 Samples & Sequencing

A total of 48 samples were selected for high coverage sequencing (Table S3). To this end, we chose the four samples with the highest concentration of extracted DNA (from each of the 12 reef locations). DNA was sent on dry ice to the New York Genome Center for library preparation using the Illumina Truseq PCR-free protocol, where they were sequenced on an Illumina HiSeqX platform in two separate batches. The first batch contained 12 samples and was sequenced across two lanes, while the second batch contained 36 samples and was sequenced across eight lanes. Due to an error at the facility, samples in the second batch were sequenced at substantially higher coverage per individual than those in the first (Figure S6; see Table S2 for coverage statistics after alignment).

#### 3.2 Read Alignment

Raw 150 bp paired-end reads were delivered as demultiplexed FASTQ files separated by lane for each sample. Low quality forward and reverse reads were first removed with *seqtk* (`https://github.com/lh3/seqtk`) using default parameters. Illumina adapters and poly-A sequences were then trimmed from the reads using Trimmomatic (v0.38)[27]. The trimmed reads were then mapped to the reference genome with `bwa` (v0.7.17) using the *mem* algorithm and default parameters[10]. `samtools` (v1.6-8)[28] was used to sort and index the binary alignments and concatenate *.bam* files across lanes for each sample.

Read group information was added to each of the sample level *.bam* files using *picard* (`https://github.com/broadinstitute/picard`) (v2.18.20). Although the libraries were generated with a PCR-free preparation, *picard* was used to mark and remove possible optical duplicates in each alignment. Following GATK "best practices" (`https://software.broadinstitute.org/gatk/documentation/tooldocs/3.8-0/org_broadinstitute_gatk_tools_walkers_indels_IndelRealigner.php`), no local indel realignment was performed prior to variant calling.

```
$SEQTK_PATH/seqtk trimfq $READ1 > $TRIM1
$SEQTK_PATH/seqtk trimfq $READ2 > $TRIM2

java -jar $TRIM_PATH/trimmomatic-0.38.jar PE -phred33 \\
        $TRIM1 $TRIM2 $TRIM1.a1.fq $TRIM1.a1.U.fq $TRIM2.a2.fq \\
        $TRIM2.a2.U.fq ILLUMINACLIP:polyA.fa:2:3:3 MINLEN:36 SLIDINGWINDOW:4:25
java -jar $TRIM_PATH/trimmomatic-0.38.jar PE -phred33 \\
        $TRIM1.a1.fq $TRIM2.a2.fq $TRIM1.b1.fq $TRIM1.b1.U.fq $TRIM2.b2.fq \\
        $TRIM2.b2.U.fq ILLUMINACLIP:adapter.fa:2:3:3 MINLEN:36 SLIDINGWINDOW:4:25
bwa mem -t24 $REF $TRIM1.b1.fq.gz $TRIM2.b2.fq.gz|\\
        samtools view -bS - > ${OUTPUT}.bwa.sam.bam
samtools sort ${OUTPUT}.bwa.sam.bam -@ 16 -O BAM -o ${OUTPUT}.bwa.sam.bam.sorted.bam
java -Xmx4G -jar $PICARD_PATH/picard.jar AddOrReplaceReadGroups \\
        I=$INPUT O=${NAME}.rg.bam RGID=$NUMB RGLB=lib1 \\
        RGPU=unit1 RGPL=illumina RGSM=$NAME
java -Xmx6G -jar $PICARD_PATH/picard.jar MarkDuplicates \\
        VALIDATION_STRINGENCY=LENIENT TMP_DIR=${ALIGNS}/tmp_dir/ \\
        I=${ALIGNS}/${IN}.rg.bam O=${ALIGNS}/${IN}.marked_duplicates.bam \\
        M=${ALIGNS}/${IN}.marked_dup_metrics.txt
```

#### 3.3 Variant Calling

GATK (v3.8)[29] was used to call variants in each sample using the HaplotypeCaller algorithm in GVCF mode. First, variants were called for each scaffold in parallel and then concatenated using GATK CatVariants to produce sample-level *.gvcf* files. GATK GenotypeGVCFs was then used to call genotypes across samples and generate a single multisample *.vcf* file. An initial principal components analysis (PCA) on the genome-wide SNPs and visual examination of the first two PC's revealed that four samples were extreme outliers (Figure S7A & Figure S7B), likely as a result of sample mis-identification or possibly the presence of cryptic species. They were further confirmed as outliers using the identity-by-state sharing test in PLINK [30], with each having a Z-score falling below the recommended threshold of $< 4$. (These four samples were removed prior to filtering and all subsequent analyses.)

Because we lacked an *A. millepora* high-quality variant set, we did not perform variant recalibration with GATK and instead applied hard-filtering thresholds, using recommendations from GATK (`https://software.broadinstitute.org/gatk/documentation/article.php?id=3225`). We excluded sites with low quality score (GQ$<$ 20); low quality by depth score (QD$<$ 10); high Fisher strand score (FS$>$ 10); low mapping quality (MQ$<$ 40); low read position rank sum score (ReadPosRankSum$< -8$); high strand odds ratio (SOR $> 4$); and low mapping quality rank sum score (MQRankSum $< -12.5$). Additionally, we masked variant calls $\pm$ 5 bp around indels and those with less than 10$\times$ coverage or greater than 2 standard deviations of the genome-wide mean coverage per sample. Any sites with more than 10% of individuals with missing genotypes were masked. We further removed all multi-nucleotide polymorphisms (accounting for 4.54% of SNPs) to create a high-quality set of biallelic 6,386,121 SNPs.

```
java -Xmx8g -jar GenomeAnalysisTK-3.8-1-0-gf15c1c3ef/GenomeAnalysisTK.jar \\
        -T HaplotypeCaller -R ${ALIGNS}/Amil.v2.00.chrs.fasta -I ${ALIGNS}/$IN \\
        -o ${IN}.${REG}.g.vcf.gz --emitRefConfidence GVCF -rf BadCigar -L $REG
java -Xmx8g -cp GenomeAnalysisTK-3.8-1-0-gf15c1c3ef/GenomeAnalysisTK.jar \\
        org.broadinstitute.gatk.tools.CatVariants -R ${ALIGNS}/Amil.v2.00.chrs.fasta \\
        $(for i in ${IN}.*.g.vcf.gz;do echo "-V ${i}";done) -out ${IN}.cat.bam.vcf \\
        -assumeSorted
java -Xmx64g -jar GenomeAnalysisTK-3.8-1-0-gf15c1c3ef/GenomeAnalysisTK.jar \\
        -T GenotypeGVCFs -R ${ALIGNS}/Amil.v2.01.chrs.fasta \\
        $(while read line;do echo "--variant ${line}";done < sorted_gVCFs.txt) \\
        --max_alternate_alleles 4 --standard_min_confidence_threshold_for_calling 30 \\
        -o Amil.merged.vcf -nt 8
bcftools -g 5 --threads 6 \\
        --output Amil.merged.44inds.indel5bp.vcf.gz Amil.merged.44inds.vcf.gz
vcffilter -i Amil.merged.44inds.indel5bp.vcf \\
        -o Amil.DPfiltered.indel5bp.biallelic.snps.vcf --snps-only \\
        --max-alleles 2 --javascript sample_filter.js --all-samples --no-gzip
vcffilter -i Amil.DPfiltered.indel5bp.biallelic.snps.vcf \\
        --all-samples -o Amil.DPfiltered.indel5bp.infoFiltered.biallelic.snps.vcf \\
        --no-gzip --keep-expr "INFO.QD > 10 && INFO.MQ > 40 && \\
        INFO.FS < 10 && INFO.SOR < 4 && INFO.ReadPosRankSum > -8 &&\\
        INFO.MQRankSum > -12.5" --min-quality 20 \\
        --exclude-bed Amil.v2.00.masked_regions.bed --max-ambiguity-ratio 0.1
```

#### 3.4 SNP Call Validation

Because we did not have access to a previous set gold-standard *A. millepora* variants, we sought to estimate error rates for our hard-filtered SNP calls. We randomly selected 10 non-repetitive 2 kb regions across the genome and designed primers (see Table S4) for each using Primer3 (`http://primer3.ut.ee`). These regions were then amplified individually in eight, randomly selected samples by PCR using high fidelity DNA polymerase (Figure S8).

##### 3.4.1 Amplicon Preparation

Each reaction was assembled as: 7.5$\mu l$ Q5 Hot Start High Fidelity 2$\times$ Master mix (NEB M0494S), 0.75$\mu l$ 10$\mu M$ forward primer, 0.75$\mu l$ 10$\mu M$ reverse primer, and 15$ng$ genomic DNA and dH2O to bring the reaction to a final volume of 15$\mu l$. PCR amplification was achieved through the following program in a BioRad thermocycler: 98°C 30 sec, [98°C 10sec 62-68°C 15sec and 72°C 1 min] $\times$ 35 cycles, 72°C 2min and 4°C hold. The exact annealing temperature ranged between 62-68°C depending on the specific set of primers. Amplicon size was validated by 1.5% Gel electrophoresis of 2$\mu l$ sample, and the rest of the reaction was further purified with Agencourt AMPure XP beads (Beckman Coulter A63881) at 0.8:1 (beads: DNA) to remove PCR reaction buffer, primers and dimmers. Purified DNA was eluted in 15$\mu l$ EB solution.

##### 3.4.2 Amplicon Library Preparation & Sequencing

Mosaic End adapter A and B (Tn5ME-A: TCGTCGGCAGCGTCAGATGTGTATAAGAGACAG; Tn5ME-B: GTCTCGTGGGCTCGGAGATGTGTATAAGAGACAG) were annealed respectively with Rev (Tn5ME-Rev: /5Phos/CTGTCTCTTATACACATCT) by mixing 10$\mu l$ (100$\mu M$) of each oligonucleotides with 80$\mu l$ of reassociation buffer (10mM Tris pH 8.0, 50mM NaCl, 1mM EDTA) in a BioRad thermocycler using the following program: 95°C for 10 min, 90°C for 1 min, followed with a decrease in temperature by 1°C/cycle for 60 cycles, held for 1 min at each.

Pre-charge of Tn5 with adapters was carried out in solution by mixing 22.5$\mu l$ of 100 $ng/\mu l$ Tn5 (Tn5 protein was produced following the protocol described by Picelli *et al.* 2014[31]), 76.5$\mu l$ reassociation buffer/glycerol (1:1), and 4.5$\mu l$ of equal molar of annealed adapter 1 (A-Rev) and annealed adapter 2 (B-Rev). The reaction was then incubated at 37°C for 30 min. The annealed adapters bind to Tn5 transposase to form the transposome complex.

Then, 20$ng$ of each amplicon was tagmented by mixing with 1$\mu l$ of above assembled Tn5 transposome, 4$\mu l$ of 5 × TAPS buffer (50mM TAPS-NaOH pH 8.5 (Alfa aesar J63268), 25mM MgCl2, 50% v/dimethylformamide (ThermoFisher 20673), pH 8.5 at 25°C) and water to a total volume of 20$\mu l$ and incubated at 55°C for 7 min. The transposome fragments and attaches adapters to amplicon. The reaction was completed by adding 5$\mu l$ of 0.2% SDS (Promega, V6551) to each reaction and incubated at 55°C for 7 min to inactivate and release Tn5.

To enrich the DNA fragments that have adapter molecules on both ends and add index to the library, 2$\mu l$ of the stopped tagmentation, 1$\mu l$ index i5 primer 1$\mu M$, 1$\mu l$ index i7 primer 1$\mu M$, 10$\mu l$ of OneTaq HS Quick-Load 2× master mix (NEB, M0486L) and 6$\mu l$ of water were combined to make a 20$\mu l$ final reaction. The reaction was heated at 68°C for 3min and 95°C for 30 sec, then thermocycled 16 times at 95°C for 10 sec, 55°C for 30 sec, and 68°C for 30 sec, followed by a final extension of 5 min at 68°C.

For multiplexing, 5$\mu l$ individual amplicon libraries from each reaction were pooled together, then cleaned and size selected using Agencourt AMPure XP beads (Beckman Coulter, A63881) at 0.8:1 (beads: DNA) ratio. The final library was quantified with a Qubit high-sensitivity DNA kit (Invitrogen Q32854) and examined on an Agilent 2100 Bioanalyzer high-sensitivity DNA chip (Agilent p/n 2938-85004) for library size distribution. The library was then sequenced on a single Illumina MiSeq lane to generate dual indexed, 150 bp paired-end reads at the Genomics Core Facility at the Lewis-Sigler Institute for Integrative Genomics at Princeton University. Each amplicon was sequenced to a mean coverage of 1454×.

##### 3.4.3 Estimation of Error Rates

The raw 150 bp paired-end reads were first demultiplexed and parsed by barcode. Low quality forward and reverse reads were then removed with *seqtk* (`https://github.com/lh3/seqtk`) using default parameters. Illumina adapters and poly-A sequences were trimmed from the reads using Trimmomatic (v0.38)[27]. Trimmed reads for each successful amplicon were then mapped with `bwa` (v0.7.17), using the *mem* algorithm and default parameters[10], to the corresponding target region extracted from the reference genome. Using `bcftools call` under default parameters, we called genotypes at every aligned site independently for each individual sample and region. We considered genotypes with fewer than 100 supporting reads as missing, and further excluded heterozygous variant calls with a $p$-value $< 10^{-3}$ for a binomial test of equal allelic balance. In total, we called 110,560 genotypes across samples and regions.

We observed an overall genotype concordance of 99.59% between the MiSeq validation and our original sequences. To assess the concordance, we conditioned on variant sites (2998 total) and assumed the high-coverage MiSeq calls to be the truth. Following the definitions of Wall *et al.* (2014)[32], we observed a false positive rate of 3.86% (116 sites) and a false negative rate of 0.33% (10 sites). All genotype errors were at heterozygous sites, *i.e.,* in no case did we observe an original homozygous non-reference call that was called homozygous reference from the MiSeq reads. Considering only variant sites across the eight individuals, $r^2$ is 0.977 between allele frequencies in validation and original variant calls and across MiSeq validation regions, indicating that even in the presence of occasional errors at the level of individual genotypes, the allele frequencies are reliably estimated.

```
#Call variant sites in amplicons
samtools mpileup -f region_fastas/${REG}.fasta -v ${SAMP}_${REG}.sorted.bam |\\
        bcftools call -c - > ${SAMP}_${REG}.vcf
```

Error rates tend to be higher for rare variants, in particular singletons[33]. In the original variant calls, 16 sites were singletons, seven of which were homozygous reference calls in the MiSeq reads; thus the error rates for singletons is significantly higher error rate than for more common variants ($p = 8.8 \times 10^{-13}$ by $\chi^2$). Scripts used to calculate concordance rates (vcf_concordance.py) and correlation across multiple samples (vcf_multisample_concordance.py) are found at the GitHub repository `https://github.com/zfuller5280/CoralGenomes`.

#### 3.5 Kinship Inference

To test for the presence of clones, siblings, or other close familial relationships among the sequenced individuals, we estimated pairwise kinship coefficients; the proportion of SNPs with 0 identity-by-state (IBS0) were estimated using KING[34]. By these approaches, no clones were identified. The maximum IBS0 estimated between any pair of samples was 0.0715 and for no comparison was a relationship higher than $3^{rd}$-degree relatedness (as defined in the KING manual) detected. Thus, all 44 samples seem to be distantly related (Figure S9).

```
king -b ${INPUT}.bed --kinship
```

#### 3.6 Characterizing Linkage Disequilibirum and Diversity, and Inferences about Demographic History

For the 44 samples, we estimated the decay of pairwise linkage disequilbrium (LD) as a function of physical distance. Pairwise LD was measured as the squared genotypic correlation coefficient ($r^2$) using PLINK[30], for a randomly chosen 1% of genome-wide SNPs. Pairwise distance comparisons were limited to 100 kb.

```
./plink --allow-extra-chr --ld-window 999999 --ld-window-kb 100 --ld-window-r2 0 \\
        --out ld_decay --r2 --thin 0.01 --vcf ${INPUT_VCF}
```

The average $r^2$ was then calculated for distances in bins of 100 bp in R:

```
library(dplyr)
library(stringr)
dfr<-read.delim("ld_decay.ld",sep="",check.names=F,stringsAsFactors=F)
dfr$distc <- cut(dfr$dist,breaks=seq(from=min(dfr$dist)-1,to=max(dfr$dist)+1,by=100))
dfr1 <- dfr %>% group_by(distc) %>% summarise(mean=mean(rsq),median=median(rsq))
```

We estimated nucleotide diversity ($\pi$, *i.e.,* the average number of single nucleotide differences between a pair of chromosomes) in non-overlapping 1 kb intergenic regions. First, we used bedtools to generate the coordinates of 1 kb windows that were entirely located between annotated gene features extracted from the GFF3 file in our assembly. We then removed windows that intersected any region that was masked or contained 'Ns' in the reference assembly. This resulted in a total of 40,137 such windows over 40.14 Mb unmasked intergenic sequence. To estimate $\pi$, we used `--site-pi` from `vcftools` in a custom Python wrapper found at `https://github.com/zfuller5280/CoralGenomes/calc_pi.py`. To handle missing data, we subtracted the number of uncallable sites from the total number of invariant sites in each window.

We applied the Pairwise Sequential Markovian Coalescent (PSMC)[35] to each genome to infer how effective population sizes ($N_e$) have changed over evolutionary time. For this analysis, we considered only scaffolds > 100 kb in size. Default parameters were used for all commands, except that we increased the stringency of the quality filter to 30 and used a bin size of 10 bp (instead of the default 100 bp), to adjust for the higher density of heterozygous sites in *A. millepora* compared to humans. Very similar results were obtained for bin sizes ranging from 5 - 50 bp. The following commands were used to generate the required input files and perform the PSMC analysis for each genome:

```
samtools mpileup -Q 30 -q 30 -u -v -f ../Amil.long_scaffolds.repeatmasked.fa \\
                 ${IN}|bcftools call -c| \\
                 vcfutils.pl vcf2fq -d 5 -D 100 -Q 30 > ${IN}.long_scaffolds.fq
fq2psmcfa -s10 ${IN}.marked_duplicates.bam.fq.long_scaffs.fq \\
                 > ${IN}.10bp.long_scaffolds.psmcfa
psmc -p "4+25*2+4+6" -o ${IN}.10bp.long_scaffolds.psmc ${IN}.10bp.long_scaffolds.psmcfa
```

We also used the Multiple Sequentially Markovian Coalescent (MSMC)[36] to infer changes in $N_e$ across samples jointly. Because of the computational demands of MSMC, we ran the analysis on a total of 12 genomes (one from each reef). The results from the MSMC analysis are qualitatively similar to the demographic histories inferred from PSMC (Figure S10). In both cases, *i.e.,* whether the analysis is performed jointly or on each genome individually, these results indicate similar coalescent histories for all individuals,and provide no evidence for population structure.

```
MEAN_COV=$(samtools depth -r ${REG} ${ALIGNS}/${NAME}.bam|\\
          awk '{sum += $3} END {print sum / NR}')
samtools mpileup -C50 -u -r ${REG} -f Amil_ref.fa ${ALIGNS}/${NAME}.s.bam|\\
          bcftools call -c -V indels|msmc-tools/bamCaller.py \\
          ${MEAN_COV} ${NAME}_${REG}_mask.bed.gz\\|
          gzip -c > ${NAME}.${REG}.msmc.vcf.gz
msmc2/msmc-tools/generate_multihetsep.py --mask=${NAME}_${REG}_mask.bed.gz \\
          --negative_mask=../Amil.exclude.regions.chrs \\
          ${NAME}.${REG}.msmc.vcf.gz > ${NAME}.${REG}.input-msmc
msmc2 -p="4+25*2+4+6" -o ${IN}.msmc.cov.res $(for i in ${IN}*input-msmc;do echo $i;done)
```

#### 3.7 Assessing Population Structure & Visualizing Effective Migration Rates

To assess population structure, we first used linkage disequilibrium (LD)-pruning to generate a set of common SNPs (MAF > 0.05) in approximate linkage equilibrium. We used the following command in PLINK:

```
./plink --allow-extra-chr --geno 0.10 --indep-pariwise 200 20 0.2 --maf 0.05\\
              --out pruned --set-missing-var-ids @:#[Amil] --vcf ${INPUT}
```

We then used the following command to perform a principal components analysis (PCA) for the 44 genomes:

```
./plink --allow-extra-chr --extract pruned.prune.in --out pruned_pca \\
             --set-missing-var-ids @:#[Amil] --vcf ${INPUT} --pca
```

We calculated the geographic distance between reefs as the haversine distance between GPS coordinates. Between each pair of sampled reef locations, we estimated the Weir-Cockerham weighted estimator of $F_{ST}$ in PLINK using the following command:

```
./plink --vcf ${INPUT}$ --extract pruned.prune.in --weir-fst-pop ${POP1} \\
           --weir-fst-pop ${POP2} --out ${POP1}_${POP2}_pairwise
```

EEMS[37] was used to visualize effective migration rates across the 12 sampled reefs using the same set of LD pruned SNPs. EEMS infers relative effective migration rates (*i.e.,* per generation migration rates scaled by $N_e$) under an equilibrium model, and under the assumption that migration occurs symmetrically between evenly spaced demes. As such, caution should be used in the interpretation of absolute values inferred by EEMS; instead these values are useful for visualizing patterns of genetic differentiation relative to geographic distance[38]. The geographic locations for each reef were taken from their GPS coordinates. PLINK was first used to convert the VCF file into binary biallelic genotype tables (*bed* format). Next, the following commands were used to convert the required input files and run EEMS software:

```
./plink --allow-extra-chr --make-bed --out Amil.hq --vcf pruned.vcf
./bed2diffs_v1 --bfile Amil.hq --nthreads 4
./runeems_snps --params params-eems_run.ini --seed 123
```

In the *.ini* parameter file, the number of demes included in the model was set to 400 and the number of MCMC iterations was set to $5\times10^6$, with a burn-in of $1\times10^6$.

#### 3.8 Characterizing Population Structure Along the Genome with *lostruct*

The method *lostruct* was used to identify localized genomic regions in which patterns of relatedness differ from what is typical for the genome. *lostruct* performs PCA in windows of a user-defined size to summarize the pattern of relatedness in each genomic region. The dissimilarity of each window to other windows is then visualized using multi-dimensional scaling (MDS). For the 44 high coverage genomes, the window size was set to 1 kb and the analysis was run using the script run_lostruct.R available at `https://github.com/petrelharp/local_pca/tree/master/templated`.

```
./Rscript run_lostruct.R -i ${INPUT_DIR} -t bp -s 1000 \
        -I genomes44/sample_info.tsv -o lostruct44_results/
```

The *sacsin* gene region showed unusual structure and was a clear outlier on the second MDS coordinate (Figure S11). The same region was also a clear outlier on the second MDS coordinate for window sizes of 2 and 5 kb.

There were two top peaks for the first MDS coordinate from *lostruct*, one on chromosome 10 and the other on chromosome 3. The top peak on chromosome 10 contained two genes, annotated as Amillepora17817 and Amillepora17828 in our assembly. For Amillepora17817, a `blastp` search revealed no significant hits to a gene with a known function, while Amillepora17828 showed homology to *cyclin-I* in *A. digitiferra*. The peak on chromosome 3 intersected the coding sequence of four genes: Amillepora23130, Amillepora23137, Amillepora23136 and Amillepora23139. However, the only significant `blastp` hits were against uncharacterized loci in *A. millepora* and *A. digitiferra*.

#### 3.9 Construction of Reference Haplotype Panel for Imputation

Using the set of filtered, high-coverage biallelic SNPs (described in **3.3**), we constructed a reference panel of haplotypes with the read-aware phasing approach implemented in SHAPEIT2[39]. First, phase informative reads (those spanning at least two heterozygous sites) were extracted using the extractPIRs tool available from SHAPEIT2. The *.vcf* files for each scaffold were then phased with SHAPEIT2 using both the standard MCMC iterative procedure and the read-aware phasing module. The raw haplotype files produced as output were then converted to phased *.vcf* files.

```
extractPIRs.v1.r68.x86_64/extractPIRs --bam $INPUT \\
        --vcf scaffold_vcfs/${NAME}.vcf --out $NAME.PIRs.list
shapeit.v2.904.3.10.0-693.11.6.el7.x86_64/bin/shapeit -assemble \\
        --input-vcf contig_vcfs/${NAME}.vcf --input-pir ${NAME}.PIRs.list \\
        -O ${NAME}.HaplotypeData --thread 4 --force
shapeit.v2.904.3.10.0-693.11.6.el7.x86_64/bin/shapeit -convert \\
        --input-haps $INPUT --output-vcf $INPUT.vcf
```

#### 3.10 $h_{12}$ Summary of Haplotype Frequencies

Using the phased haplotypes generated for the creation of a reference panel, we looked for unusually high frequencies of the two most common haplotypes in localized genomic regions with the $h_{12}$ statistic[40]. $h_{12}$ is a summary statistic that considers the frequencies of the first and second most common haplotypes in a sample in a single measure, as

$$h_{12} = (p_1 + p_2)^2 + \sum_{i>2}^{n} p_i^2$$

where $p_i$ is the frequency of the $i^{th}$ most common haplotype in the sample, and there are $n$ distinct haplotypes. $h_{12}$ was estimated with a custom Python script available at (`https://github.com/zfuller5280/CoralGenomes/hapStats.py`) using a window size of 100 SNPs centered around each focal site. We searched for loci with highly unusual values of $h_{12}$ by considering those values in the top 0.001%. genome-wide. A peak of unusual $h_{12}$ values was detected on chromosome 7, falling over the *sacsin* gene (Figure S11). These outlier SNPs were located in a region between coordinates 13323256-13327878, which falls entirely within the coding sequence of *sacsin*. This signal is consistent with the presence of two haplotypes at high frequency in the region.

The frequencies of the two most common haplotypes did not show a significant association (see Table S5) with either of the top two genetic PCs or the top eight PCs constructed from environmental variables (see section **7.1.1**).

##### 3.11 Visualizing Gene Trees for *sacsin* Versus Random Genomic Regions

To infer the gene tree for *sacsin*, we first randomly selected one phased high-coverage genome from each of the 12 sampled reefs. Then, for each individual, we randomly extracted one haplotype for a central 1 kb region in the gene (13326340-13327340) in a .vcf file. We next converted the variant calls in the .vcf to a .fasta file spanning this region. To obtain alignments with *A. digitifera* and *A. tenuis*, we extracted this same region from our reference *A. millepora* assembly for use as a BLASTN query and took the top hit against both reference genomes. The sequences for the 12 *A. millepora* samples and the *A. digitifera* and *A. tenuis* genomes were then aligned using MUSCLE. The script to extract the region of interest from the high-coverage haplotypes and outgroup species and convert all sequences to a .fasta file is found at (`https://github.com/zfuller5280/CoralGenomes/makeOutgroupPhasedFastas.py`). A maximum likelihood phylogeny was then constructed from the alignment using the program *dnaml*. A modified version of the SNPhylo pipeline was used to perform the local alignment and phylogenetic inference, available at (`https://github.com/zfuller5280/CoralGenomes/snphylo.sh`).

We then randomly selected 1000 autosomal 1 kb loci (from un-masked regions in the referenec) and used the same workflow as above to construct gene trees for each. We plotted this set of 1000 random 1 kb regions and the *sacsin* gene tree using DensiTree. This visualization is shown in Figure 4C in the main text.

```
#Sample and extract random 1kb regions
bedtools random -l 1000 -n 1000 -seed 16 -g Amil.genome > random_regions
bedtools shuffle -i random_regions -incl gene_regions.bed -g Amil.genome \\
        -excl Amil.v2.01.masked_regions.bed > random_1kb_gene_regions
COUNT=1;
while read line;
        do CHROM=$(echo $line|cut -f1 -d" ");
        START=$(echo $line|cut -f2 -d" ");
        STOP=$(echo $line|cut -f3 -d" ");
        bcftools view -r ${CHROM}:${START}-${STOP} \\
        -s DRR108008,AN02,CS11,DK04,FR20,FY13,HH16,JB05,NB16,PA21,RB13,RL18,TR02 \\
        phased.vcf.gz > ${COUNT}.random.gene_tree.vcf;((COUNT+=1));
done < random_1kb_gene_regions
```

### 4 Validation of *sacsin*

The gene *sacsin* has unusually high diversity, reflective of unusually deep coalescence times among lineages. These signals are consistent with our expectations for a gene under long-term balancing selection. In our *A. millepora* genome assembly, the coding region is located on Chromosome 7 spanning the coordinates 13318370 to 13331252 and is annotated as Amillepora11972. It was identified as showing homology to *sacsin* in *A. digitifera* after running blastn and `blastp` on the nucleotide and amino acid sequence, respectively. We first interrogated the coverage distribution in *sacsin* to test for the presence of a collapsed paralog or split allele in our assembly. We would expect two collapsed paralogs to show increased read depth relative to the mean coverage, while we would expect a split allele to show decreased relative read depth[41]. We did not observe a significant change in read depth over the gene relative to the average coverage for any of the 44 high-coverage resequenced genomes (Figure S12).

##### 4.1 Targeted Resequencing of *sacsin*

To further validate that the diversity patterns at sacsin were not a bioinformatic or assembly artifact, we selected four samples (DK06, NB16, PA07, and RL08) for targeted amplification and resequencing of *sacsin*.

The choice of individuals was made on the basis of available quantity of genomic DNA remaining after the previous whole-genome sequencing, while ensuring that at most one individual was selected per sampled reef, and that at least one predicted heterozygote and one of each of the homozygote haplotypes would be sequenced.

To conduct the validation, we designed 22 primer pairs across the *sacsin* region using Primer3 (see Table S6 for sequences, Figure S13 for locations over the region). Each primer pair produced overlapping amplicons of between 1819-2231 bp. These amplicons were then digested with Tn5 to produce short fragments tagmented with adaptor sequences for subsequent enrichment. During enrichment barcodes were added to allow demultiplexing after sequencing.

Reaction mixtures for PCR of these amplicons were composed of 0.25$\mu l$ Phusion High-Fidelity DNA Polymerase (NEB E0553L), 5$\mu l$ 5X Phusion HF Buffer, 0.5$\mu l$ 10mM dNTPs, 0.75$\mu l$ 10$\mu M$ forward primer, 0.75$\mu l$ 10$\mu M$ reverse primer, 56ng, 50ng, 50ng, and 38.8ng genomic DNA were used respectively for samples DK06, NB16, PA07, and RL08, and nuclease-free water to bring the reaction to a final volume of 25$\mu l$. PCR conditions used were: 98°C 30 sec, [98°C 10sec 65°C 30sec and 72°C 1 min]×30 cycles, 72°C 10min and 4°C hold. To prepare Tn5 for tagmentation it must be combined with annealed adapters to form a transposome complex[31]. Annealed adapters were obtained for each of Mosaic End adapters A and B (Tn5ME-A: TCGTCGGCAGCGTCAGATGTGTATAAGAGACAG; Tn5ME-B: GTCTCGTGGGCTCG-GAGATGTGTATAAGAGACAG) by mixing 8$\mu l$ of the respective adapter (100$\mu M$) with 8$\mu l$ of the Rev adapter (100$\mu M$, Tn5ME-Rev: CTGTCTCTTATACACATCT) and 4$\mu l$ of Reassociation Buffer (10mM Tris pH 8.0, 50mM NaCl, 1mM EDTA). Annealing conditions were 95°C 3 min, 90°C 30 sec, reduce temperature by 0.5°C every 30 sec 70 times, and 4°C hold. 2.7 $\mu l$ of Tn5 was mixed with 1.65 $\mu l$ of each annealed adapter. This was incubated at 23°C for 1 hour and diluted 5 fold to produce a diluted transposome complex.

For each sample, the 22 amplicons were tagmented by mixing 2 $\mu l$ diluted transposome complex, 1 $\mu l$ PCR product, 4 $\mu l$ TAPS buffer, and nuclease-free water to bring the reaction to a final volume of 20 $\mu l$, and incubating for 7 min at 55°C. To prevent excessive digestion, the reaction was then killed by the addition of 2.5 $\mu l$ 0.2% SDS to each reaction and incubating for 7 min at 55°C. Tagmented amplicons were then enriched for fragments containing the adapter sequences by a subsequent PCR. Forward and reverse primers used for enrichment were i5 and i7 indexes, allowing barcoding for subsequent pooling and demultiplexing of samples. Reaction mixtures contained 4$\mu l$ of killed tagmentation reaction, 10 $\mu l$ OneTaq HS Quick-Load 2×master mix (NEB, M0486L), 2 $\mu l$ of each primer (2.5 $\mu M$ for amplicons derived from RL08 and 10 $\mu M$ for amplicons derived from DK06, NB16, and PA07), and nuclease-free water for a final volume of 20 $\mu l$. PCR conditions used were: 72°C 3 min, 94°C 30 sec, [94°C 10 sec 62°C 15 sec and 68°C 30 sec]×17 cycles, 68°C 5min and 4°C hold.

1$\mu l$ of each barcoded tagmentation reaction was pooled to form a library. This library was then cleaned and size selected over three washes using Agencourt AMPure XP beads (Beckman Coulter, A63881). First, a left-sided size selection (bead: DNA of 0.6) was used to remove large fragments, followed by two right-sided size selections, the first with a bead: DNA of 0.2 and the second with a bead: DNA of 0.8. The final library was quantified with a Qubit high-sensitivity DNA kit (Invitrogen Q32854) and examined on an Agilent 2100 Bioanalyzer high-sensitivity DNA chip (Agilent p/n 2938-85004) for library size distribution. The library was then sequenced on a single Illumina MiSeq lane to generate dual indexed, 250bp paired-end reads at the Genomics Core Facility at the Lewis-Sigler Institute for Integrative Genomics at Princeton University.

#### 4.2 Sequence and Variant Call Validation

We took two approaches to verify the sequence of *sacsin* in our reference genome and to validate SNPs called in the region. First, we pooled reads from amplicons for each sample and performed a *de novo* assembly in a reference-free approach. Second, we mapped the reads from each amplicon to our reference genome and then aligned the assembled amplicon sequences.

##### 4.2.1 *De novo* Assembly

For each sample, we constructed *de novo* assemblies with the paired-end reads generated from the sequenced amplicons. We first used a custom Python script (`https://github.com/JaneliaSciComp/msg/blob/master/barcode_splitter.py`) to parse the raw reads by i5 and i7 indices. Using commands listed under **2.2**, low quality forward and reverse reads were then removed with *seqtk* (`https://github.com/lh3/`

`seqtk`) using default parameters. Illumina adapters and poly-A sequences were trimmed from the reads using Trimmomatic (v0.38)[27]. For each sample, the trimmed reads were then pooled across amplicons. Reads were then assembled with spades[42] under default parameters and using 16 threads. Iterative $k$-mer lengths of 21, 33, 55, 77, 99 and 127 were automatically detected based on read length.

```
spades.py -o ./spades_${SAMPLE} -1 ${SAMPLE}.r1.fq.gz -2 ${SAMPLE}.r2.fq.gz
```

The longest continuous assembled sequences for each sample were:

- DK06 - 12995 bp
- NB16 - 2170 bp
- PA07 - 4845 bp
- RL08 - 4897 bp

We aligned the assembled scaffolds for each sample to the *sacsin* region from the reference assembly using ClustalW with default parameters. From the original genomic variant calls and phased haplotypes, DK06 and NB16 were both predicted to carry homozygous haplotypes over *sacsin* that were similar to the reference sequence. The longest scaffold for DK06 aligned over the entire coding sequence starting at 11 bp before the start codon and extending 102 bp downstream, while the longest scaffold for NB16 began aligning 2007 bp into the coding sequencing. RL08 was predicted to have the alternate homozygous haplotype for *sacsin*, and the longest scaffold started aligning 4382 bp into the coding sequence. PA07 was predicted to be heterozygous for both haplotypes; we aligned assembled scaffolds for the two haplotypes that overlapped in the coding sequence, confirming their presence. The longest scaffold, more similar to the reference sequence, began aligning to *sacsin* 3858 bp downstream from the start codon. The next longest assembled scaffold (3749 bp) was more similar to the alternate haplotype and started aligning 4111 bp downstream from the start codon. Assembled scaffolds for each sample are deposited and available for download at `https://github.com/zfuller5280/CoralGenomes/data/`. These results validate the *sacsin* gene sequence in the reference assembly and confirm the presence of diverged haplotypes in the region for the samples sequenced.

##### 4.2.2 Amplicon Mapping

In addition to a reference free *de novo* assembly approach, we also mapped the reads from each amplicon to our *A. millepora* genome to i) confirm their expected order, ii) have a second method to validate the reference sequence over the *sacsin* gene and iii) estimate the error rate in the original SNP calls in the gene. For this analysis, we used the sample from each predicted homozygous haplotype that had the greatest coverage (DK06 and RL08). We first aligned the demultiplexed and trimmed reads from each amplicon to the *sacsin* region in the reference assembly using `bwa-mem` with default parameters. We then sorted and marked duplicates in each alignment. For calling variants, we used `bcftools` to create pileup files and the `vcf2fq` tool to call consensus sequences for each amplicon. We considered genotypes with less than 50× coverage as missing.

```
bwa mem sacsinregion.fa $READS1 $READS2| \\
        samtools sort - -@ 4 -O BAM -o ${OUTPUT}.sorted.bam
java -Xmx6G -jar picard.jar MarkDuplicates VALIDATION_STRINGENCY=LENIENT \\
        TMP_DIR=tmp_dir/ I=${IN}.sorted.bam \\
        O=${IN}.marked_duplicates.bam M=${IN}.marked_dup_metrics.txt
bcftools mpileup -f sacsinregion.fa ${IN}.bam |\\
        bcftools call -c - | bcftools/misc/vcfutils.pl vcf2fq > ${IN}_consensus.fa
```

We aligned the resulting FASTA sequences for each amplicon to the reference *sacsin* gene using ApE (`https://jorgensen.biology.utah.edu/wayned/ape/`) and manually inspected their order based on the overlapping primer design. SnapGene (`www.snapgene.com`) was used to visualize the alignments of amplicons (Figure S14). For both DK06 and RL08, the order of aligned amplicon sequences matched their expected position based on successfully amplified primer pairs. This result further confirmed the presence of both alternative divergent haplotypes in the region and validated the reference sequence for *sacsin*. In both samples, we observed small deletions relative to the reference sequence approximately 2 kb downstream of the *sacsin* coding region (see Figure S14).

##### 4.2.3 Estimation of Error Rates

To compare variant calls in the sequenced amplicons to the original SNP calls, we first concatenated the sorted BAM files for alignments of each sample. We then generated VCF files using `bcftools` tools and applied the following filters:

```
bcftools filter -e 'DP<50 | QUAL<20 | PV4[0]<0.01| MQ<40 | \\
            PV4[1]<0.01 | PV4[2]<0.01 | PV4[3]<0.01 | HWE<0.001'
```

The PV4 of VCF files generated with `bcftools` field contains $p$-values for strand bias, baseQ bias, mapQ bias and tail distance bias. Genotypes failing any of the filters above were converted to missing *i.e.,* 'N'. A VCF file containing the called genotypes for *sacsin* in RL08 and DK06 can be found at `https://github.com/zfuller5280/CoralGenomes/data/combined.sacsin.calls.vcf`. In each sample, we first considered each SNP in the original genotype calls. Of the 454 original SNP calls, a total of 18 and 20 were missing (*i.e.,* N) in the MiSeq amplicon validation calls for RL08 and DK06, respectively. For all other non-missing variants, if we assume the MiSeq validation calls to be the truth, we confirmed each called SNP in *sacsin* for the original sequences of DK06 and RL08 (Table S7). (We note that no variants for either sample were singletons in the original sequences, likely contributing to the lower false positive rate observed in *sacsin* than in the randomly selected validation regions in **3.4**). Again assuming the MiSeq calls to be the truth, we observed false negative rates of 1.43% and 0.43% for RL08 and DK06, respectively.

### 5 Genotype Imputation

#### 5.1 Algorithm

The algorithm we use for genotype imputation in low-coverage sequencing data is based on the copying model of Li and Stephens (2003)[43] and implemented in the software *loimpute* developed by Gencove, Inc. These methods are originally described in Wasik *et al.* (2019) and for completeness are reproduced here. In brief, assume we have a set of $N$ bi-allelic variants that have been genotyped on a set of $M$ phased haplotypes, such that the allele at variant $i$ on haplotype $j$ is $h_{ij}$, and coded as a 0 if the allele matches the reference genome and as a 1 if the allele matches the alternate allele at the site. At each of the $N$ sites, we additionally have a set of $L$ sequencing reads from the individual whose genotypes we would like to impute, such that the allele at variants $i$ at sequencing read $k$ is $r_{ik}$. We now wish to impute the genotype of the (diploid) target individual at each of the variants.

Now let $X_{i,1}$ and $X_{i,2}$ be the identity of the haplotypes being copied at variant $i$ by the target individual. Similarly to Li and Stephens (2003)[43], the transition probabilities from $X_{i,1}$ to $X_{i,2}$ can be modeled as:

$$P(X_{i+1,1} = x'|X_{i,1} = x) = \begin{cases} \exp(-\rho d_j/M) + (1 - exp(-\rho d_j/M))(1/M) & \text{if } x' = x \\ (1 - exp), & \text{otherwise} \end{cases}$$

where $\rho$ is the population-scaled recombination rate per base and $d_j$ is the physical distance between the variants at position $i$ and $i+1$. The setup for the second haplotype of the individual is identical.

Now let $Y_i = X_{i,1} + X_{i,2}$. The emission probabilities of each read given $Y_i$ can be modeled as:

$$P(r_{ik} = r|Y_i = y) = \begin{cases} \epsilon & r = 0, y = 2 \\ 1-\epsilon & r = 0, y = 0 \\ 0.5 & y = 1 \\ 1-\epsilon & r = 1, y = 2 \\ \epsilon & r = 1, y = 0 \end{cases}$$

where $\epsilon$ is the sequencing error rate. The posterior probability of each genotype in the target individual can be estimated using standard methods for hidden Markov models. Note that this algorithm has time complexity $O(NM^2)$.

#### 5.2 Expected Imputation Accuracy

Prior to sequencing the remainder of the samples at low coverage, we assessed the accuracy of our genotype imputation procedure implemented in *loimpute* with a cross-validation approach. We first randomly selected four samples (CS07, DK06, RB13, TR10) to use as a validation set, and repeated **2.5** to construct a reference haplotype panel with these individuals removed. Using `samtools`, we then randomly subsampled reads from the alignments for each of the validation samples on Chromosome 1 to mimic low-pass sequencing at various mean target coverages. We compared the accuracy of our imputation method against the performance of BEAGLE5[44] using the squared linear correlation coefficient ($r^2$) between the true (*i.e.,* high-coverage) genotypes and most probable imputed genotypes (Figure S15A) for each set of subsampled sites. $r^2$ was averaged across sites and samples as in Huang *et al.* 2009[45] to obtain a single summary for each mean target coverage. Additionally, for each of the validation samples, we estimated the non-reference discrepancy for a range of MAF cutoffs (Figure S15B):

$$\frac{\#\ of\ discordant\ sites}{\#\ of\ total\ sites\ \text{-}\ \#\ concordant\ reference\ sites}$$

*loimpute* had significantly greater accuracy and lower non-reference discrepancy than BEAGLE5 across various minor allele frequency (MAF) thresholds and each mean target coverage examined (Figure S15A and Figure S15B). We achieved an accuracy ($r^2$) $> 95\%$ and non-reference discrepancy $< 5\%$ for variants with MAF $> 0.05$ in the reference haplotype panel at a target coverage of $3\times$ and greater. For variants with MAF $> 0.05$, our imputation approach outperformed previous studies in humans with small reference panels[44–46] and other non-model organisms such as salmon[47] and cattle[48]. The success of genotype imputation here is presumably in large part due to the favorable ratio of LD and diversity in *A. millepora.*

```
samtools view -bs ${RNUM}.${DOWNSAMPLE} ${IN}.marked_duplicates.bam chr1|\\
        samtools mpileup - > ${IN}.downsampled_${COVERAGE}x.pileup
samtools view -bs ${RNUM}.${DOWNSAMPLE} ${IN}.marked_duplicates.bam chr1|\\
        samtools mpileup -f ${REF} -v - > ${IN}.downsampled_${COVERAGE}x.pileup.vcf
gzip ${IN}.downsampled_${COVERAGE}x.pileup
REF_PANEL="chr1.HaplotypeData.vcf.gz"
loimpute -i ${IN}.downsampled_${COVERAGE}x.pileup.gz -h ${REF_PANEL}\\
        -o ${IN}.${COVERAGE}_imputed -k 80 -id ${IN}
gunzip ${IN}.${COVERAGE}_imputed.vcf.gz
bcftools call -c ${IN}.downsampled_${COVERAGE}x.pileup.vcf > ${IN}.${COVERAGE}x_sites.vcf
beagle.28Sep18.793.jar impute=true gt=${IN}.${COVERAGE}x_sites.vcf\\
        ref=chr1.HaplotypeData.vcf.gz.bref3 out=${IN}.${COVERAGE}x.beagle.imputed
gunzip ${IN}.${COVERAGE}x.beagle.imputed.vcf.gz
```

### 6 Low Coverage Alignment & Variant Calling

#### 6.1 Samples & Sequencing

Libraries for DNA extracted from the remaining 209 samples were prepared for multiplexed shotgun genotyping (MSG) with Tn5 transposase and tagmentation[31].

##### 6.1.1 Library Preparation

First, Mosaic End adapters A and B (Tn5ME-A: TCGTCGGCAGCGTCAGATGTGTAT AAGAGACAG; Tn5ME-B: GTCTCGTGGGCTCGGAGATGTGTATAAGAGACAG) were annealed respectively with Rev (Tn5ME-Rev: /5Phos/CTGTCTCTTATACACATCT) by mixing 10$\mu l$ (100$\mu M$) of each oligonucleotide solution with 80$\mu l$ of reassociation buffer (10mM Tris pH 8.0, 50mM NaCl, 1mM EDTA) in a BioRad thermocycler using the following program: 95°C for 10 min, 90°C for 1 min, followed with a decrease by 1°C/cycle for 60 cycles, held for 1 min at each temperature. Pre-charge of Tn5 with adapters was carried out in solution by mixing 22.5$\mu l$ of 100 $ng/\mu l$ Tn5 (Tn5 protein was produced following the protocol described by (Picelli *et al.* 2014), 76.5$\mu l$ reassociation buffer/glycerol (1:1), and 4.5$\mu l$ of equal molar of annealed adapter 1 (A-Rev) and annealed adapter 2 (B-Rev). The reaction was then incubated at 37°C for 30 min. The annealed adapters bind to Tn5 transposase to form the transposome complex.

$20ng$ of gDNA was then tagmented by mixing with $1\mu l$ of the above assembled Tn5 transposome, $4\mu l$ of 5× TAPS buffer (50mM TAPS-NaOH pH 8.5 (Alfa aesar J63268) , 25mM $MgCl_2$, 50% v/dimethylformamide (ThermoFisher 20673) , pH 8.5 at 25°C) and water to a total volume of $20\mu l$ and incubated at 55°C for 7 min. The transposome fragments and attaches adapters to amplicon. The reaction was completed by adding $5\mu l$ of 0.2% SDS (Promega, V6551) to each reaction and incubated at 55°C for 7 min to inactivate and release Tn5.

To enrich the DNA fragments that have adapter molecules on both ends and add indices to the library, $2\mu l$ of the stopped tagmentation, $1\mu l$ index i5 primer $1\mu M$, $1\mu l$ index i7 primer $1\mu M$, $10\mu l$ of OneTaq HS Quick-Load 2× master mix (NEB, M0486L) and $6\mu l$ of water were combined to make a $20\mu l$ final reaction volume. The reaction was heated at 68°C for 3min and 95°C for 30 sec, then thermocycled 16 times at 95°C for 10 sec, 55°C for 30 sec, and 68°C for 30 sec, followed by a final extension of 5 min at 68°C.

For multiplexing, $5\mu l$ individual libraries from each reaction were pooled together, and purified using Agencourt AMPure XP beads (Beckman Coulter, A63881) at 0.8:1 (beads: DNA) ratio. Following the protocol described above, two separate pooled libraries were constructed. The first library was size-selected to be between 350-750bp, whereas the second library did not include a size selection step. Both libraries were quantified with a Qubit high-sensitivity DNA kit (Invitrogen Q32854) and examined on an Agilent 2100 Bioanalyzer high-sensitivity DNA chip (Agilent p/n 2938-85004) for library size distribution. Each library was then sequenced on an Illumina HiSeq4000 platform to generate dual indexed, 150bp paired-end reads at Genewiz. The size-selected library was sequenced across three lanes, while the non-size-selected library was sequenced across seven lanes.

#### 6.2 Read Alignment

Following Schumer *et al.* (2018)[49], raw reads were first parsed by index, using a custom Python script `https://github.com/JaneliaSciComp/msg/blob/master/barcode_splitter.py`. Using commands listed under **2.2**, low quality forward and reverse reads were then removed with *seqtk* (`https://github.com/lh3/seqtk`) using default parameters. Illumina adapters and poly-A sequences were trimmed from the reads using Trimmomatic (v0.38)[27]. Trimmed and demultiplexed reads for each sample were then mapped to our *A. millepora* reference genome assembly with `bwa` (v0.7.17) using the mem algorithm and default parameters[10]. Read group information was added to each of the sample and library level *.bam* files using `picard` (`https://github.com/broadinstitute/picard`) (v2.18.20). `samtools` (v1.6-8)[28] was used to sort and index the binary alignments and concatenate *.bam* files across lanes and libraries for each sample. `picard` "MarkDuplicates" was used to mark PCR and optical duplicates in each concatenated alignment file. First, we assessed the alignment metrics for each library using "CollectInsertSizeMetrics" and "CollectWgsMetricsWithNonZeroCoverage" tools from `picard`:

```
picard.jar CollectInsertSizeMetrics I=${INPUT} O=${NAME}.inserts.metrics \\
             R=Amil.v2.01.chrs.fasta HISTOGRAM_FILE=${NAME}.reads.hist \\
             VALIDATION_STRINGENCY=SILENT
picard.jar CollectWgsMetricsWithNonZeroCoverage I=${INPUT} O=${NAME}.nz.metrics \\
            R=Amil.v2.01.chrs.fasta CHART_OUTPUT=${NAME}.nz.pdf\\
            VALIDATION_STRINGENCY=SILENT
```

We realized that a large fraction of aligned bases were excluded in both the size selected and non-size selected libraries due to reads marked as duplicates, overlapping reads, and read removal because of low mapping quality (MAPQ $< 20$) (Figure S16). The non-size selected library had a larger proportion of aligned bases excluded because of overlapping reads than the size selected library. Meanwhile, the size selected library had a larger proportion of aligned bases removed because of reads marked as duplicates than the non-size selected library. Thus, we combined reads from all sequenced libraries and lanes for alignment in the final dataset. For these concatenated reads, we then removed samples with an average coverage $< 0.5\times$ after alignment. After this initial filtering step, 193 samples remained. The mean coverage for these 193 samples was 1.43× (median = 1.40), with a max coverage of approximately 3.5×.

#### 6.3 Genotype Imputation & Variant Calling

We then imputed genotypes in the 193 samples using *loimpute* and the reference haplotype panel constructed in **2.5**. First, we generated pileup files for each alignment using `samtools` (v1.6-8)[28], which were then used as input for *loimpute*. We considered genotype probability thresholds of 0.90, 0.95, and 0.99 for imputed genotype calls to be considered as non-missing for hard calls.

```
samtools mpileup -r ${REG} /merged_bams/${IN}.marked_duplicates.bam \\
            > ${IN}.${REG}.pileup
gzip ${IN}.${REG}.pileup
loimpute -i ${IN}.${REG}.pileup.gz -h ref_vcfs/${REG}.vcf -k 86 \\
            -id ${IN} -gpf ${GP_CUTOFF} -o ${IN}.${REG}.95.loimpute;
```

In addition to the 193 samples in the Tn5 libraries were 34 individuals originally sequenced at high-coverage, described in **2**. Therefore, we assessed the accuracy of our imputation approach by comparing the high-coverage calls with the imputed genotypes at each of the genotype probability thresholds (Figure S17A). Consistent with our down-sampling simulation approach described in **2.6**, we saw an overall $r^2 > 90\%$ for all samples $> 0.5\times$ coverage at any of the genotype probability thresholds we considered. In fact, we observed an improvement from our original simulations: $r^2 = 94\%$ for an average coverage of $1.5\times$, for genotypes with a probability $\geq 0.95$. While the overall mean accuracy of imputed genotypes genome-wide improved if we increased the stringency of the genotype probability threshold to 0.99, on average each sample had more than 10% of sites removed as a result (Figure S17B). As a compromise, we used a genotype probability threshold of 0.95 for analyses requiring hard calls.

#### 6.4 Removing Close Relatives and Outliers

From the initial set of individuals that passed our filtering and the original high-coverage genomes, we had a set of 237 samples with genome-wide resequencing. We tested for the presence of clones or other highly related individuals by estimating kinship coefficients and the proportion of pairwise IBS0 using KING (see section **2.5**). While the average kinship coefficient between pairs of individuals was -0.08, indicating very distant relationships, three pairs had a kinship coefficient $> 0.4$, pointing to clones. No other comparison had an inferred relationship greater than $3^{rd}$-degree relatives (*i.e.,* kinship coefficient $> 0.0884$). One individual from each highly related pair was selected randomly and removed from further analysis. These individuals were: NB08, HH30, and HH15.

We further used IBS sharing to detect potential outliers using PLINK[30]. Pairwise IBS distance was calculated using all SNPs and five nearest neighbors were identified for each individual. The IBS distance to each of the five nearest neighbors was then transformed into a Z-score. Individuals with a minimum Z score among the five nearest neighbors less than -4 (as recommended by PLINK documentation) were excluded from analysis as population outliers. These individuals were: HH23, JB14, FR14, CS06, JB08, JB09, JB11, JB12.

```
./plink --file ${INPUT} --cluster --neighbour 1 5
```

We also performed a second, independent analysis of IBS distance to identify clones and samples likely collected from the wrong species. Here, we used the single-read sampling approach implemented in ANGSD[50] to estimate pairwise distances among all sequenced individuals. Importantly, because ANGSD uses single read sampling, this method is robust to variation in sequencing coverage across individual samples[50] (see Manzello *et al.* 2019 for a similar analysis using 2bRAD data in the coral *Orbicella faveolata*). A hierarchical clustering was performed on the pairwise IBS distances using the `hclust()` function in `R`. A dendrogram of this clustering is shown in Figure S18. This additional analysis identified the same individuals as outliers and clonal pairs as detected using PLINK. The outlier individuals were further visually confirmed in a PCA (Figure S19). After removing individuals from pairs of identified clones and outliers (likely the result of sample mis-identification[51]), the sample size of sequenced individuals was 226.

#### 6.5 Analysis of Population Structure, Inference of Effective Migration Rates, and Characterization of Genetic Diversity in the Full Dataset

To investigate if the signatures of limited population structure, high gene flow, and the same peaks of elevated genetic diversity were present in the full dataset of all 226 sequenced individuals, we re-ran a PCA, EEMS, and genome-wide scan of $\pi$ for all samples. Similar to the results using 44 genomes, we found little evidence of population structure or clustering in a PCA (Figure S20A). The results from modeling relative migration rates with EEMS were qualitatively similar and no strong barriers to gene flow were inferred between any of the sampled reefs (Figure S20B). These results indicate that there is minimal population structure among the individuals included in our study. The strongest signal genome-wide of elevated diversity measured by $\pi$ in 1kb windows was again detected at *sacsin* (Figure S20C).

### 7 GWAS

We performed genome-wide association studies (GWAS) for bleaching measured as either a visual score, a total chlorophyll content and a symbiont cell density. For each GWAS, we removed samples with missing phenotype data, bringing the total sample size down to 213, 190 and 172, respectively. The rank-ordering among pairs of phenotype values for the three measures of bleaching are highly correlated, with a Spearman's correlation coefficient $> 0.6$ for each (Figure S21).

#### 7.1 Covariates

The final set of covariates included in all GWAS models were: the top four PCs of environmental and spatial variables, the first two genetic PCs, the collection date, the sequencing batch (see section **3.1**), collection depth and the relative proportion of *Symbiodinium* Clade D reads (out of all symbiont reads). We refer to the collection date and sequencing batch as batch effects.

##### 7.1.1 Environmental PCs

For each reef sampling site, we used the environmental and spatial variables from Matthews *et al.* (2019) as well as GPS coordinates for latitude and longitude. This set of variables are: ‘CRS_NO3_AV’, ‘CRS_NO3_SR ‘CRS_PO4_AV ‘CRS_PO4_SR,’ ‘CRS_O2_AV’, ‘CRS_O2_SR’, ‘CRS_S_AV’, ‘CRS_S_SR’, ‘CRS_T_AV’, ‘CRS_T_SR’, ‘CRS_SI_AV’, ‘CRS_SI_SR’, ‘GA_BATHY’, ‘GA_SLOPE’, ‘GA_ASPECT’, ‘GBR_BATHY’, ‘GA_CRBNT’, ‘GA_GRAVEL’, ‘GA_SAND’, ‘GA_MUD’, ‘GMCS_STRESS_TMN’, ‘GMCS_STRESS_IQR’, ‘SW_CHLA_AV’, ‘SW_CHLA_SR’, ‘SW_K490_AV’, ‘SW_K490_SR’, ‘SW_BIR_AV’, ‘SW_BIR_SR’, ‘MT_SST_AV’, ‘MT_SST_SR’, ‘MT_SST_MIN’, ‘Primary’, ‘Secondary’, ‘Tertiary’, ‘Latitude’, ‘Longitude’, ‘DHW_max’, ‘mindistbar’, ‘mindist-coa’

Because many of the variables are highly correlated (*e.g.,* latitude and temperature), we performed a PCA. In the analysis reported in the main paper, we used the top four PCs, which together account for $> 85\%$ of the total variance, to include as covariates (see **7.7** for results of a GWAS instead using six or eight environmental PCs). We note further that the environmental variables are not significantly correlated with the genetic PCs (see Table S8).

| | PC1 | PC2 | PC3 | PC4 | PC5 | PC6 | PC7 | PC8 |
|---|---|---|---|---|---|---|---|---|
| Standard Deviation | 4.03 | 3.04 | 2.13 | 1.74 | 1.44 | 1.39 | 0.97 | 0.77 |
| Proportion of Variance | 0.41 | 0.24 | 0.12 | 0.08 | 0.05 | 0.05 | 0.02 | 0.01 |
| Cumulative Proportion | 0.41 | 0.65 | 0.77 | 0.85 | 0.90 | 0.95 | 0.97 | 0.99 |

##### 7.1.2 Composition of Symbiont Types

We mapped the final set of paired-end reads from each sample to a modified reference genome containing our *A. millepora* assembly and four draft genomes for *Symbiodinium microadriaticum*, *Breviolum minutum*, *Cladocopium goreau* and *Durusdinium trenchii*, formerly *Symbiodinium* clades A, B, C1 and D, respectively. These draft genomes were available from:

- *Symbiodinium microadriaticum* - `http://reefgenomics.org/`

- *Breviolum minutum* - Shoguchi *et al.* (2013)[52]
- *Cladocopium goreau* - Liu *et al.* (2017)[53]
- *Durusdinium trenchii* - Katherine Dougan/M. Rodriguez-Lanetty (personal communication)

Because of the size of the symbiont sequences, each genome was split into contigs of length 10 Mb. Paired-end reads from each sample were mapped and the alignments sorted, duplicates marked and read groups added, using the same workflow as in **3.2**. The counts of non-duplicate reads mapping to each chromosome, scaffold and contig were tabulated using `samtools` with the following command:

```
samtools view -f 2 -F 3176 $BAM|cut -f3|sort|uniq -c|sort -k1nr|\\
        awk 'OFS="\t"{print $1,$2$}' > ${BAM}.seq
```

The read counts were summed across the split symbiont genome contigs to obtain the total read counts $r$ per genome $k$ for each sample. Following the measure of genome relative abundance in Xia et al (2011)[54], the relative abundance ($a$) of each symbiont taking into account genome length $l$ for each sample $i$ was estimated as:

$$a_i = \frac{\frac{\pi_i}{l_i}}{\sum_{k=1}^{n} \frac{\pi_k}{l_k}}$$

where

$$\pi_i = \frac{\frac{r_i}{l_i}}{\sum_{k=1}^{n} \frac{r_k}{l_k}}$$

The relative symbiont abundance $a_{Sym}$ per sample was found by summing $a_i$ for the four symbiont genomes. The relative composition of each symbiont type (*i.e.,* clade) per sample was then estimated by dividing $a_i$ for each symbiont genome by this total $a_{Sym}$. A script containing functions to calculate read counts and estimate $a_i$ and $a_{Sym}$ can be found at `https://github.com/zfuller5280/CoralGenomes/symbiont_genomes.py`. The relative symbiont abundance $a_{Sym}$ per sample was highly correlated with all three measures of the bleaching phenotype. Plots showing this relationship for total chlorophyll content and symbiont cell density are shown in Figure S22, while the symbiont read abundance across quartiles of visual scores is shown in Figure 5A in the main text.

#### 7.2 Estimating the SNP Heritability ($h^2$)

We estimated the proportion of phenotypic variance explained by common genome-wide SNPs (*i.e.,* the SNP heritability $h^2$) for quantile-normalized visual bleaching scores using the software package GCTA[55]. First, the genetic relationship matrix (GRM) was estimated for all pairs of individuals from autosomal SNPs with MAF $> 0.05$. To this end, we ignored sites containing more than 10% missing individuals, and filtered out SNPs with Hardy-Weinberg $p$ values below $10^{-7}$. Next, a restricted maximum likelihood analysis was performed using the GRM and a set of covariates composed of the sequencing batch, collection date, collection depth, the top four environmental PCs, the top two genetic PCs and the relative proportion of Clade D symbiont reads. The discrete and quantitative covariates were input as separate tables as required by GCTA.

```
./gcta -bfile Amil.gcta_samps --autosome --maf 0.05 --make-grm --out gcta_grm \\
        --autosome-num 14
./gcta --pheno amil.vis_score.norm.gcta.txt --covar amil_covars.gcta.tsv \\
        --qcovar amil_qcovars.gcta.tsv --out Amil.gcta \\
        --reml --reml-alg --grm gcta_grm
```

The estimate of $h^2$ output from GCTA was 0.497. However, the standard error (SE) of this estimate is 1.21, meaning that the 95% confidence interval (calculated as $h^2 \pm 1.96\times$ SE) spans the entire interval 0 to 1, and no meaningful inference can be made from this estimate of $h^2$. Similar results were obtained for total chlorophyll content and symbiont cell density. This result is not surprising, as the individuals are very distantly related, so that even for a trait that is in fact highly heritable, tens of thousands of individuals may be required for a precise estimate of SNP heritability[56].

#### 7.3 Linear Mixed Model

We performed a GWAS for visual score using a linear mixed model (LMM) implemented in the software GEMMA[57,58]. Although we detected little to no population structure among the sequenced individuals and across reefs, we used an LMM for the GWAS to account for subtle effects of population stratification and genetic relatedness between samples when testing for an association between phenotype and genotype. GEMMA fits a univariate LMM in the following form:

$$\mathbf{y} = \mathbf{W}\alpha + \mathbf{x}\beta + \mathbf{u} + \epsilon; \mathbf{u} \sim MNV_n(0, \lambda\tau^{-1}\mathbf{K}), \epsilon \sim MNV_n(0, \tau^{-1}\mathbf{I}_n),$$

where $\mathbf{y}$ is a vector of length $n$ containing phenotypes for $n$ samples; $\mathbf{W} = (\mathbf{w}_1 ..., \mathbf{w}_c)$ is an $n \times c$ matrix of fixed effects (*i.e.,* covariates) for $c$ covariates and including a column of 1s; $\alpha$ is a $c$-vector of the corresponding coefficients for the covariates including the intercept; $\mathbf{x}$ contains the genotypes in an $n$-vector; $\beta$ is the effect size at the tested site; $\mathbf{u}$ is an $n$-vector of random effects; $\epsilon$ in an $n$-vector containing the errors; $\tau^{-1}$ is the variance of the residual errors; $\lambda$ is the ratio between the two variance components; $\mathbf{K}$ is a known $n \times n$ genetic relatedness matrix and $\mathbf{I}_n$ is an $n \times n$ identity matrix. The $n$-dimensional multivariate normal distribution is denoted by $MNV_n$. At each SNP the alternative hypothesis $H_1 : \beta \neq 0$ was tested against the null hypothesis $H_0 : \beta = 0$ using the likelihood ratio score (as recommended in the GEMMA manual).

For the input genotypes, we used imputed genotype dosages in BIMBAM format. We ignored sites containing more than 10% missing individuals. Additionally, we filtered out SNPs with Hardy-Weinberg $p$ values below $10^{-7}$ and with a minor allele frequency (MAF) less than 0.05. We first estimated the relatedness matrix with GEMMA using a set of SNPs in approximate linkage equilibrium after LD pruning in PLINK. This matrix $\mathbf{K} = G_c$ was calculated as the centered relatedness matrix in the following form:

$$G_c = \frac{1}{p}\sum_{i=1}^{p}(\mathbf{x}_i - \mathbf{1}_n\bar{x}_i)(\mathbf{x}_i - \mathbf{1}_n\bar{x}_i)^T,$$

where $\mathbf{X}$ is defined as the $n \times p$ matrix of genotypes and $\mathbf{x}_i$ as the $i$th column of genotypes at the $i$th SNP; $\bar{x}_i$ is the sample mean and $\mathbf{1}_n$ is is an $n$-vector of 1s. Before performing the GWAS, we used quantile-normalization to transform the phenotypes.

```
#LD Prune SNPs
./plink --allow-extra-chr --geno 0.10 --indep-pairwise 200 20 .2 \\
      --maf 0.05 --out pruned --remove outliers_missing_rmv \\
      --set-missing-var-ids @:#[Amil] --vcf ${INPUT}
#Run GEMMA
gemma -bfile gemma.plink_nonmissing.pruned -gk 1 \\
      -o Amil.relatedness -maf 0.05 -hwe 1e-7 -miss 0.10
gemma --g Amil.bimbam.nonmissing.filled.geno -p Amil.vis_score.gemma.pheno \\
-a Amil.bimbam.snps  -k output/Amil.relatedness.cXX.txt \\
-lmm 4 -o Amil.lmm -c gemma.prop_symb.vis_score.4PCs_covars -maf 0.05 -miss 0.10
```

We also performed a GWAS using imputed genotype dosages for standardized total chlorophyll content and standardized symbiont cell density. For each, we used the same covariates and parameters for GEMMA as in the GWAS for visual score and similarly quantile-normalized the phenotypes. Individuals with missing phenotype data were removed, bringing the sample size to $n = 190$ for total chlorophyll content and $n = 172$ for symbiont cell density. Manhattan plots for these GWAS are shown in Figure S23A and Figure S23B. While variants within the peak observed on chromosome 14 in the GWAS for visual score had high $p$-values ($> 0.01$) for either measurement of bleaching, SNPs within the peak observed on chromosome 13 were among the top signals for both, with minimum $p$- values of $2.21 \times 10^{-5}$ and $2.78 \times 10^{-5}$ in the region for total chlorophyll content and symbiont cell density, respectively. However, similar to the GWAS for visual score, no variants for either measurement had $p$-values less than the cutoff we determined for genome-wide significance (see section **7.6**).

#### 7.4 General Linear Model

In addition to an LMM, we performed a GWAS using standard linear regression as implemented in PLINK2[30]. Here, we used the same covariates as the LMM and similarly quantile-normalized the phenotypes. The rank-ordering of $p$ values was highly correlated between the standard linear GWAS and the LMM GWAS for visual score, with a Spearman correlation coefficient = 0.848 (Figure S24).

To further test the effects of latitudinal population structure, we separated individuals into eight northern (Arlington, Fitzroy, Russell, Coates, Feather, North Barnard, Dunk, Taylor) and four southern (Rib, John Brewer, Pandora, Havannah) reefs, performed a GWAS and compared the estimated effect sizes. For covariates, separate genetic PCs were calculated in the southern and northern reefs (Figure S25). For SNPs with $-\log_{10}(p) > 4$ in the GWAS using only samples from northern reefs, there was a correlation of $r^2 = 0.903$ with $\beta$s estimated at those same sites in the GWAS using only individuals from the southern reefs. Similarly, for SNPs with $-\log_{10}(p) > 4$ in the GWAS using only samples from southern reefs, there was a correlation of $r^2 = 0.895$ with $\beta$s estimated at those same sites in the GWAS using only individuals from the northern reefs.

We note that the dominant symbiont type may itself be partly heritable (*i.e.,* under genetic control). If so, and if some of the variants that impact bleaching also influence the symbiont type, then it should not be included as a covariate in the model. In practice, removing it had little effect on the GWAS results. The top peaks on chromosome 13 and chromosome 14 observed in the GWAS for visual score were among the top peaks when the symbiont type was removed as a covariate and Spearman's correlation coefficient between effect size $\beta$ estimates in both GWAS models was 0.71 (Figure S26). We additionally performed a GWAS using the dominant symbiont type as a phenotype (and accordingly removed it as a covariate). Only one SNP genome-wide had a $-\log_{10}(p) > 4$ (Figure S27).

#### 7.5 Bayesian Association Test

We also performed an association test for SNPs with visual score in a Bayesian framework using the software BIMBAM[59]. Here, Bayes Factors are computed for a linear regression of quantitative phenotype values on genotype under the following model

$$Y_i = \mu + aX_i + dI(X_i = 1) + \epsilon_i$$

where $Y_i$ indicates the phenotype value for individual $i$, $X_i$ is the genotype for indiviual $i$, $a$ indicates the additive effect, $d$ is the dominance effect and $\epsilon_i$ represents the error term, which is assumed to be iid normal. We performed the association test using the default parameters of BIMBAM, with the prior D2 from Servin and Stephens (2007)[59] for additive and dominance effects by averaging over $\sigma_a = 0.05, 0.1, 0.2, 0.4$ and $\sigma_d = \sigma_a/4$. Imputed genotype dosages in BIMBAM format were used as input. BIMBAM does not accept covariates as input. Thus, we first regressed the visual scores on the covariates used in the LLM and standard linear GWAS, and used the resulting quantile normalized residuals as the input phenotype for BIMBAM (as recommended in the software manual). Sites with MAF $< 0.05$ and with more than 10% missing genotypes were excluded from the association test.

```
#R
quantNorm =function(x){qnorm(rank(x,ties.method = "average")/(length(x)+1))}
model.residual = lm(Bleaching_score ~ X.D + PC1 + PC2 + PC3 + PC4
                + as.factor(HC_batch2) + as.factor(LC_batch) + EV1 + EV2
                + Date + Depth, data=Amil_vs_covars)
residuals<-resid(model.residual)
qnorm_residuals<-quantNorm(residuals)
write.table(qnorm_residuals[gemma_samp_order],"residualized_phenos.vs.4PCs.pheno",
          quote=F,row.names=F,col.names=F)

#Run bimbam
bimbam-lin -p visual_scores.4PCs.resid.pheno -gmode 1 -g ${GENO} \\
         -pos ${SNPS} -o bimbam_out -exclude-maf 0.05 -exclude-miss 0.10
```

A Manhattan plot showing the Bayes Factors at each SNP for an association with the normalized visual score is presented in Figure S28.

#### 7.6 Determining Genome-wide Significance

The large number of tests performed in a GWAS needs to be taken into account when assessing statistical significance. The common approach is to set a threshold of "genome-wide significance." In human genetics, most GWAS commonly set a cutoff of genome-wide significance at $5 \times 10^{-8}$, which is equivalent to setting a Bonferroni correction of 1 million independent tests (*i.e.,* SNPs) at $\alpha = 0.05$, and is a cut-off beyond which associations are observed to reliably replicate. This value depends on the number of SNPs tested and patterns of linkage disequilibrium along the genome, so will differ across species. Here, we set this threshold by performing a permutation test. Specifically, we shuffled the phenotype values of visual scores across the 213 individuals 10,000 times; for each permutation, we performed a standard linear GWAS using the same covariates and assessed the minimal $p$-value. We then set the genome-wide significance threshold as the $95^{th}$ percentile of the distribution of these minimal permuted $p$-values, equal to $4.293 \times 10^{-8}$. One interpretation is that we expect that approximately 5% of random assignments of phenotypes would yield a $p$-value that low or lower than observed in reality somewhere in the genome[60]. The distribution of minimal $p$-values from the permutation test is shown in Figure S29. We also performed similar permutation tests for total chlorophyll content and symbiont cell density, obtaining genome-wide significance thresholds of $4.631 \times 10^{-8}$ and $5.380 \times 10^{-8}$, respectively.

#### 7.7 Construction of the Polygenic Score (PGS) & Assessment of Prediction Accuracy

In our sample of 213 sequenced individuals phenotyped for visual score, we lacked a true out-of-sample validation set to assess the accuracy of polygenic scores constructed from GWAS. Thus, we used a jackknife cross-validation (CV) procedure to subset the samples into 100 partitions of training and test sets. In each partition, we randomly withheld 15% of individuals as the test set and selected the other 85% of individuals as the training set. We re-calculated the genetic PCs for the training set, and used the loadings from this PCA for the genetic PCs in the test set. A script to randomly separate samples into train and test sets and to recalcualte covariates in each is available at (`https://github.com/zfuller5280/CoralGenomes/make_covars_cv.R`)

For each jackknife partition, a standard linear GWAS for quantile-normalized visual score was performed on the training set, using the top four PCs from environmental and spatial variables, the first two genetic PCs, the collection date, collection depth, sequencing batch and relative proportion of *Symbiodinium* Clade D reads as covariates. Only common variants (MAF $\geq 0.05$) were considered and sites with more than 0.10 of genotypes missing were ignored. To build the PGS, we used $p$-value thresholding followed by LD-clumping to choose sets of approximately independent SNPs. For each set of selected SNPs, the PGS was calculated by summing the allelic dosages weighted by their estimated effect size. To assess prediction accuracy, a PGS was then estimated in the test set. We first determined the $p$-value threshold that maximized the average prediction accuracy of the PGS (measured as $R^2$)[61,62] in the test across the 100 jackknife partitions by considering a logarithmically spaced range of $p$-value thresholds: $\{1 \times 10^{-1}, 1 \times 10^{-2}, 1 \times 10^{-3}, 1 \times 10^{-4}, 1 \times 10^{-5}\}$. $R^2$ was maximized for a $p$-value threshold of $1 \times 10^{-5}$ (Figure S30).

Next, using a $p$-value threshold of $1 \times 10^{-5}$ we tested whether individuals in the higher end of the PGS distribution tended to have a higher actual visual score than those individuals in the low end of the PGS distribution. Within the test set of each partition, we divided the samples into quartiles based on their PGS and calculated the average visual score for each. We then tested for a significant difference between the distributions of average visual scores across the jackknife partitions for the highest and lowest quartile with a Mann-Whitney U test. These distributions of average visual scores are depicted in Figure 6B of the main text. A script to calculate the average visual score in quartiles and plot their distribution can be found at `https://github.com/zfuller5280/CoralGenomes/pgs_accuracy.R`.

We then sought to assess the prediction accuracy of the PGS along with other known influences of bleaching, by examining the change in $R^2$ in the test set for linear models including different combinations of predictors. We considered five different models:

1. sequencing batch + collection date
2. sequencing batch + collection date + top 2 genetic PCs

3. sequencing batch + collection date + top 2 genetic PCs + collection depth + top 4 environmental PCs
4. sequencing batch + collection date + top 2 genetic PCs + collection depth + top 4 environmental PCs + relative proportion of Clade D reads
5. sequencing batch + collection date + top 2 genetic PCs + collection depth + top 4 environmental PCs + relative proportion of Clade D reads + PGS

For each jackknife partition, we estimated $R^2$ in the test set for linear models using the combinations of predictors above. We tested for differences in the distribution of $R^2$ across partitions with Mann-Whitney U tests. A script to fit the linear models listed above, to test for differences in the average $R^2$ and to plot Figure 6C in the main text can be found at `https://github.com/zfuller5280/CoralGenomes/pgs_accuracy.R`.

The PGS provided a significant increase in the average $R^2$ across partitions for training-test splits of 90/10% and 80/10% (Figure S31). The lowest $p$-value (0.010) was obtained for a training-test split of 90/10%, while a higher $p$-value (0.033) was observed for a training-test split of 80/20%.

Lastly, to examine the influence of the choice of environmental and spatial variables on the prediction accuracy of the PGS, we re-ran the same analysis as above but used either six or eight top environmental PCs instead of four. The PGS provided a significant increase in the average $R^2$ across partitions when using either 6 ($p = 0.026$) or 8 ($p = 0.029$) environmental PCs (Figure S32).

### 8 Supplemental Figures

**Figure S1.** Schematic of assembly workflow. **(A)** The construction of a PacBio-only assembly with Canu and a consensus PacBio assembly scaffolded with 10X Chromium barcodes with Falcon. Arrows show the input and output of each step in the assembly, with the program/software name indicated in the dark blue box. Colors of the arrows denote whether the input is derived from PacBio (cyan) or 10X Chromium/Illumina (green) reads. **(B)** The workflow used to map pooled larval reads to the assemblies and filter endosymbiont/contaminant sequences. The filtered assemblies were then merged and a final error correction step performed to create a set of contigs. These contigs were then scaffolded into chromosomes using published linkage maps.

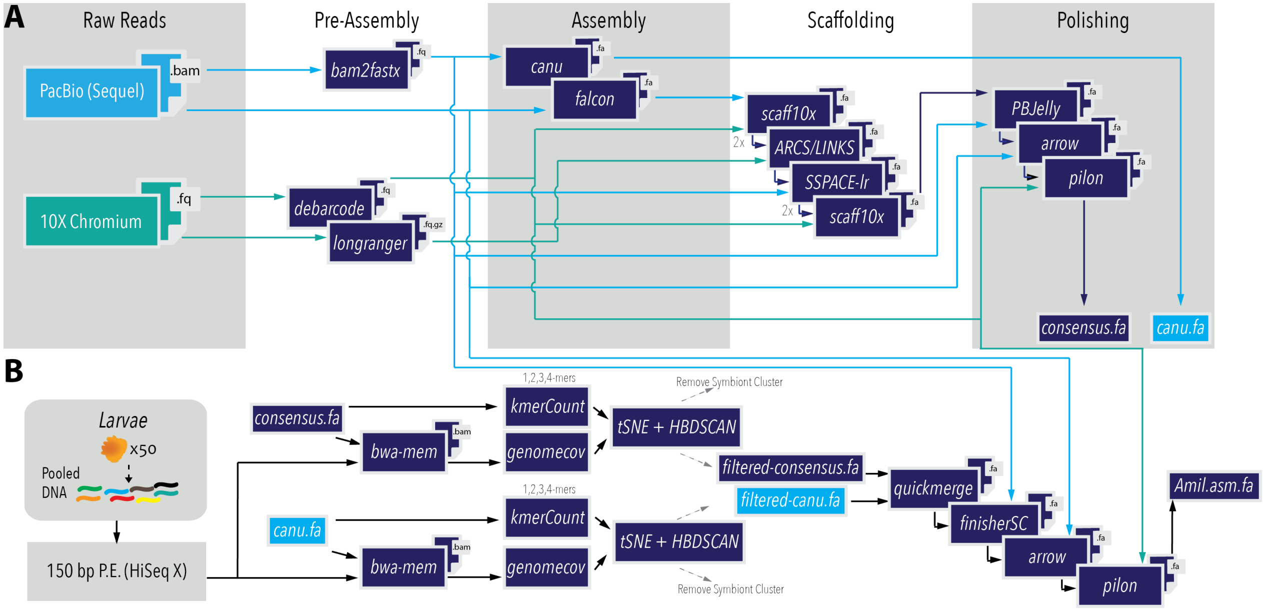

**Figure S2.** $k$-mer profile analyzed with GenomeScope. A total genome length of 447 Mb was estimated with a duplicated/repeat content of 30.4%.

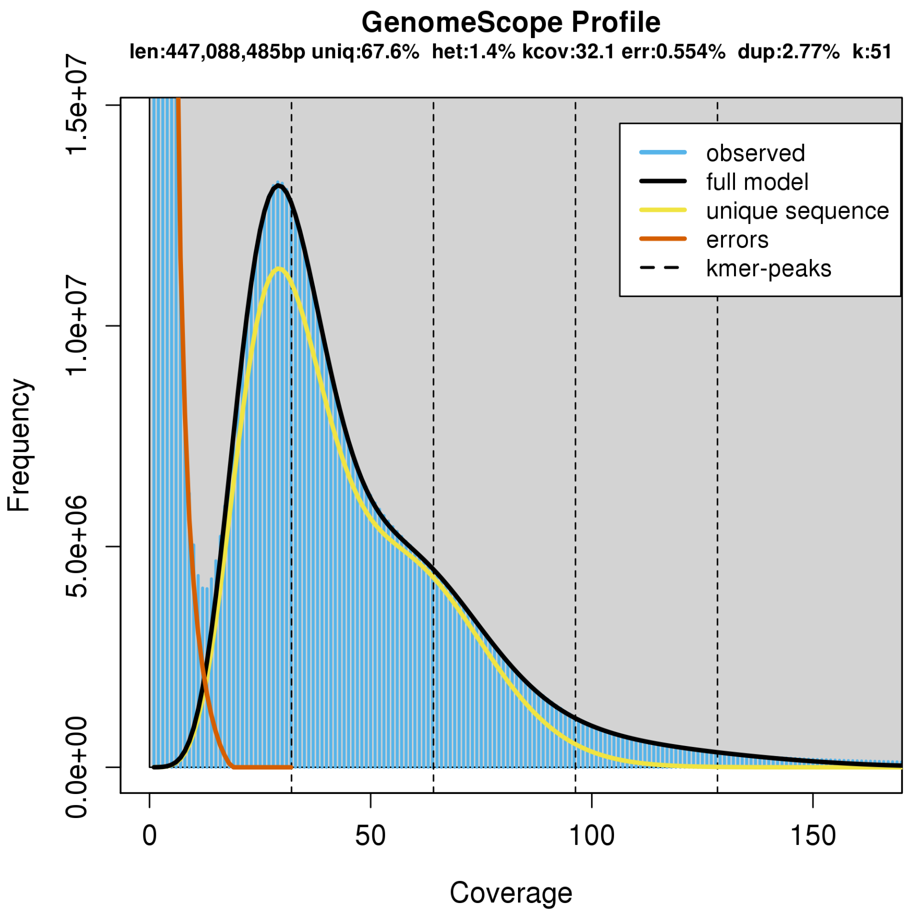

**Figure S3.** t-SNE dimensionality reduction of sequence composition and coverage features to separate putative coral and endosymbiont contigs. Dimensionality reduction was performed for contigs in the "consensus" (left) and Canu (right) assemblies independently. The size of the circles in each plot represent the size of each contigs and colors represent the labels assigned using HBDSCAN clustering. Those circles colored blue are inferred to belong to the cluster of coral contigs, as confirmed using `blastn`; and those colored in green indicate contigs likely assembled from reads of *Symbiodinium spp.*

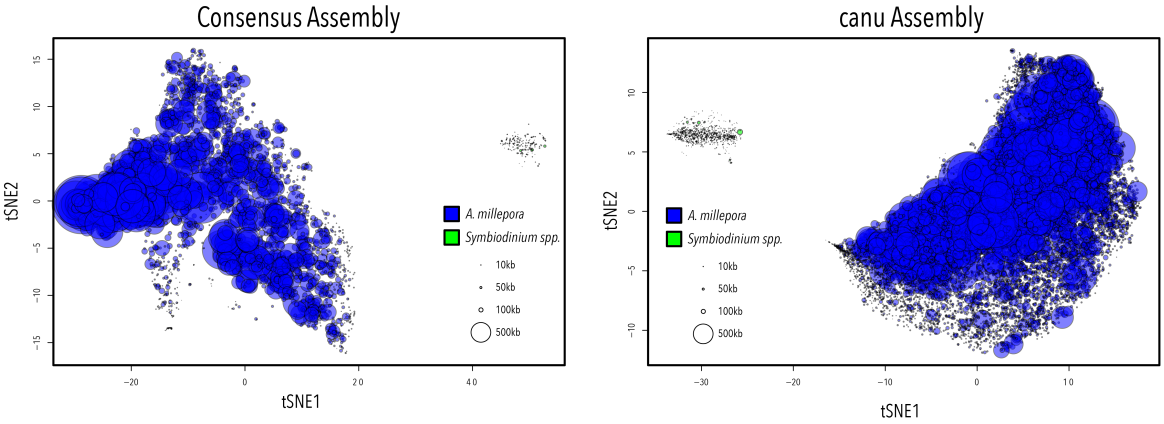

**Figure S4.** Schematic showing the alignment of scaffolds from the published Ying *et al.* assembly[23] (in black) and the 14 chromosome-scale scaffolds constructed in the assembly presented in this article (in gray). On the bottom, the alignments for chromosome 1 and chromosome 14 are shown in detail, with the scaffold names in the Ying *et al.* assembly shown above each.

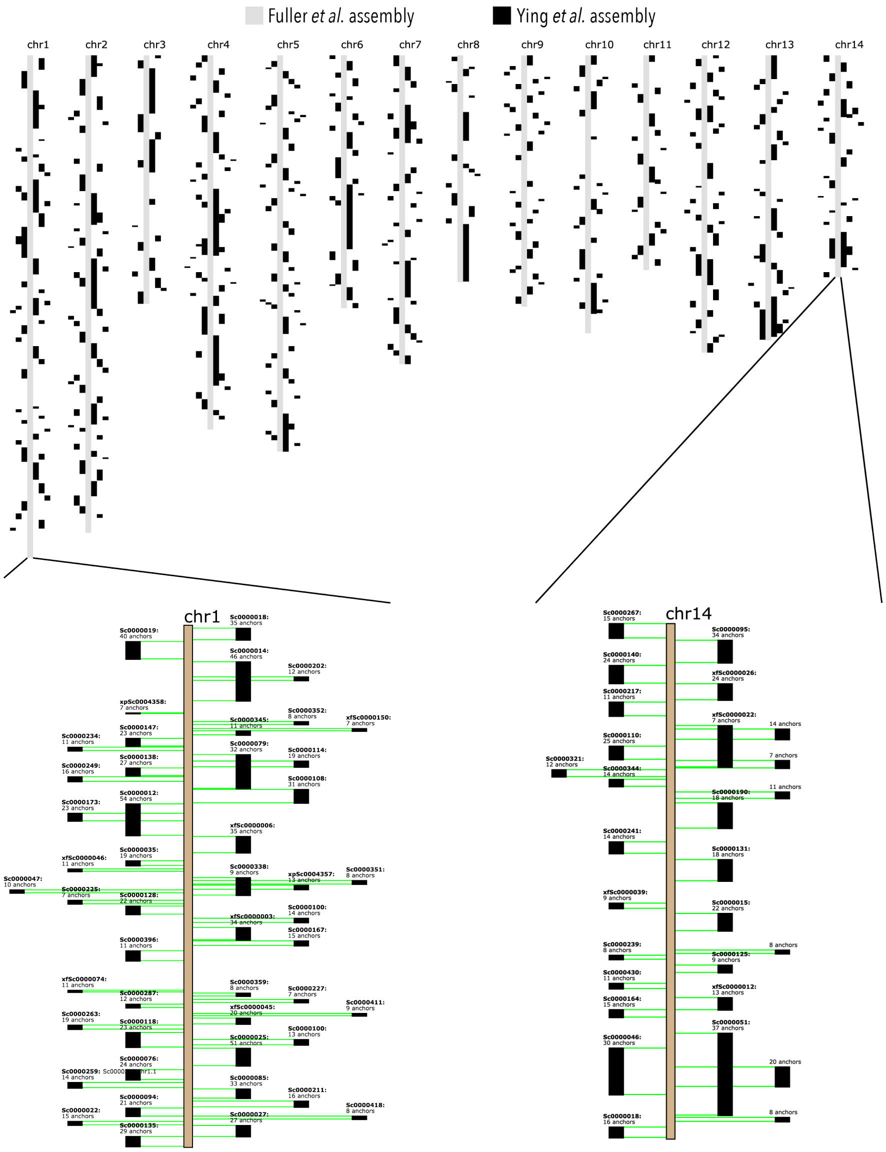

**Figure S5.** Circular plot showing the alignment of scaffolds from the published Ying *et al.* assembly[23] (top half) and the 14 chromosome-scale scaffolds constructed in the assembly presented in this article (bottom half). Each ribbon connects the corresponding position of each scaffold in the alignment. This visualization shows the high level of fragmentation present in the Ying *et al.* assembly[23] as well as multiple regions in the chromosome-scale assembly presented here which are not contained in the Ying *et al.* assembly[23].

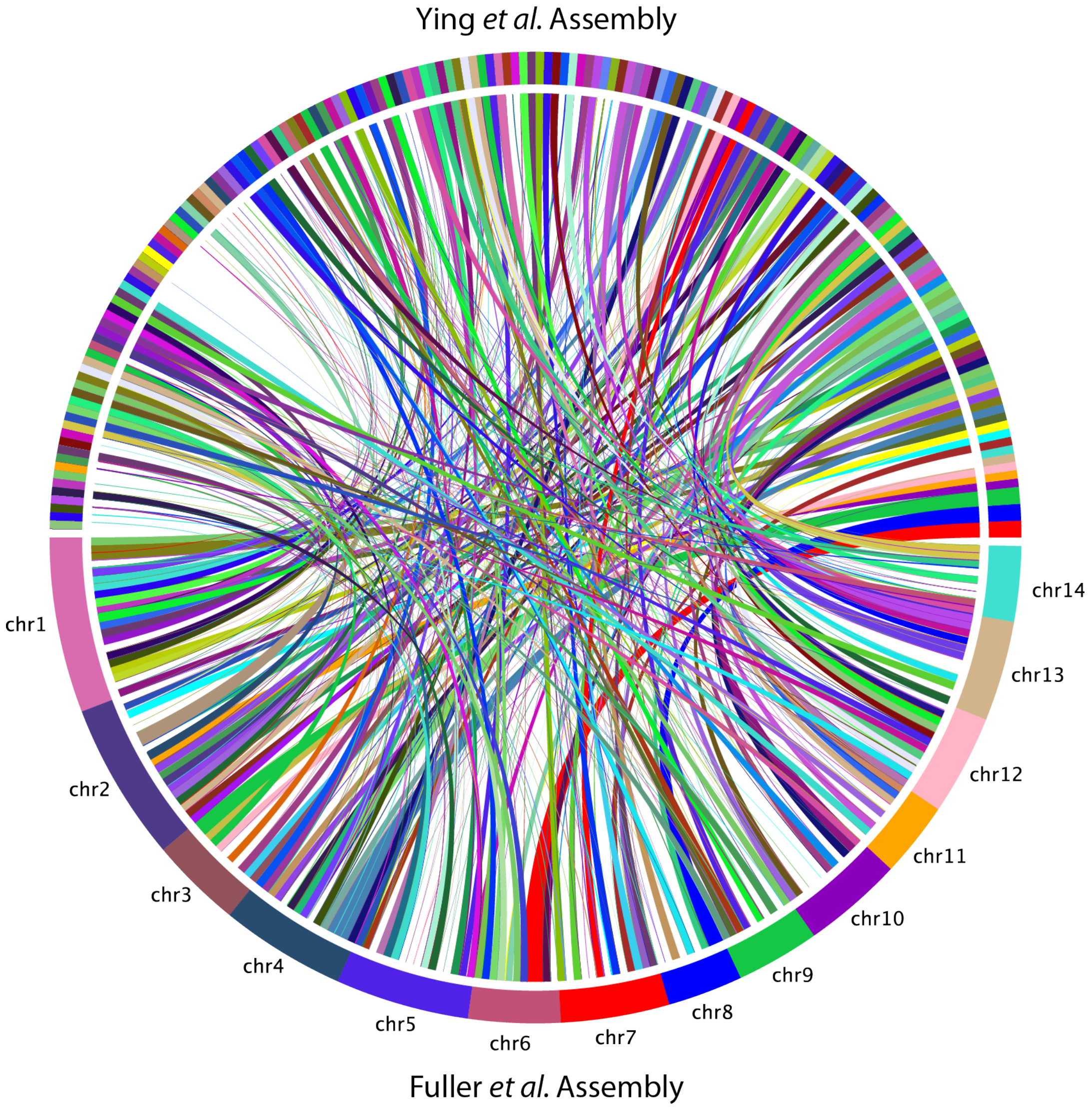

**Figure S6.** Distribution of genome-wide coverage for the 48 samples sequenced at high coverage. Dashed lines represent samples sequenced in Batch 1, and solid lines represent samples sequenced in Batch 2. The 36 samples sequenced in Batch 2 have substantially higher coverage due to an error at the sequencing facility.

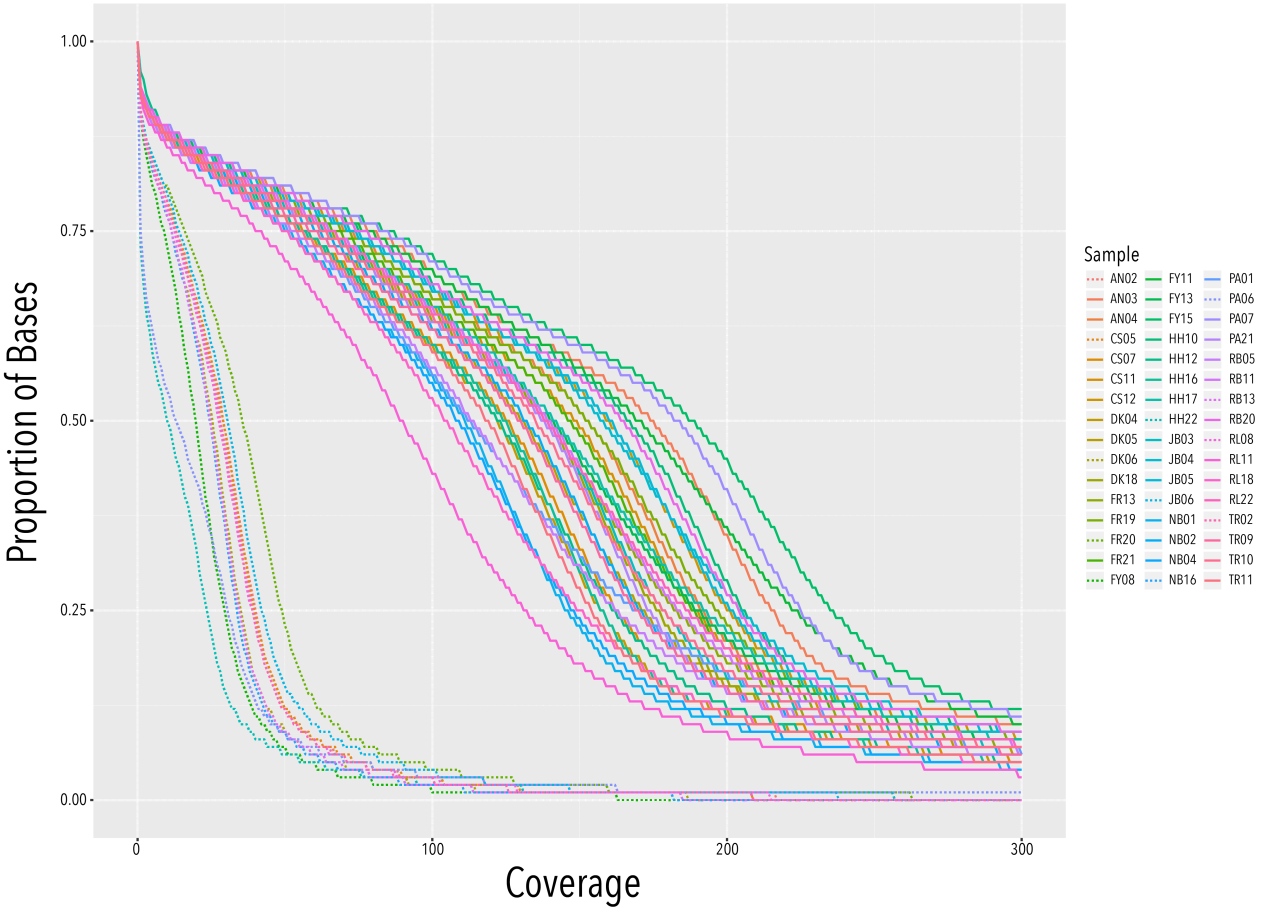

**Figure S7.** Principal components analysis (PCA) on genome-wide SNPs for samples sequenced at high coverage. **(A)** A PCA for all 48 samples. As shown on the first PC, HH22 and PA06 account for 54% of the total variation and are separated from all other samples. Both samples were removed from subsequent analyses as outliers. **(B)** A PCA for 46 samples, with HH22 and PA06 removed. Here, FY08 and JB06 account for 21% of the total variation and are identified as further outliers. Moreover, all four samples were identified as outliers using PLINK with Z-scores > 4 for a test of IBS distance.

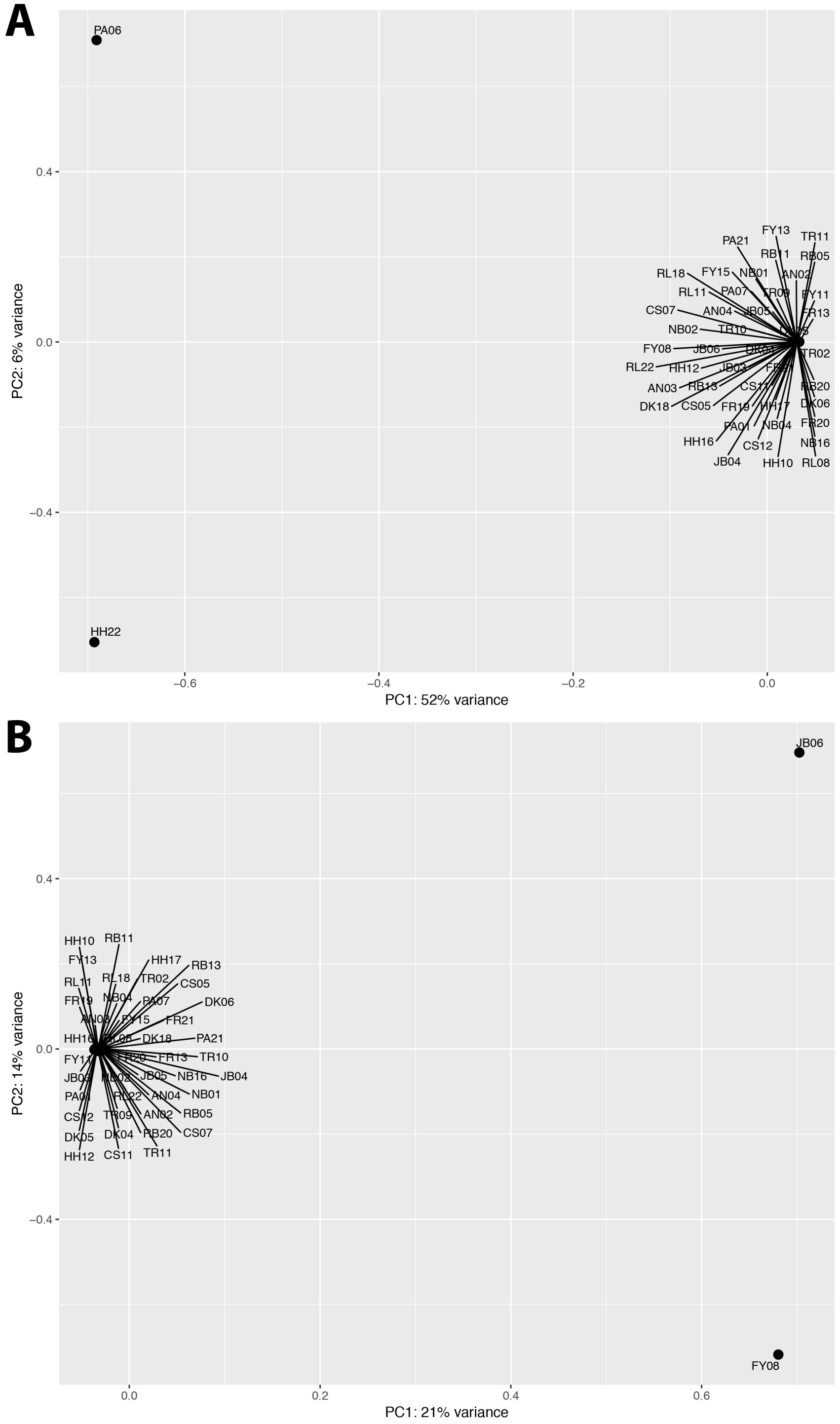

**Figure S8.** PCR amplification success for validation regions. Each region is represented as a row, while samples are represented as columns. Cells in dark green indicate regions that were successfully amplified, while those in light green indicate regions that successfully amplified yet had a mean coverage $< 100\times$ and were therefore not considered. PCR amplifications that failed are marked with blue "Neg".

| **Coral** | 1 | 2 | 3 | 4 | 5 | 6 | 7 | 8 |
|---|---|---|---|---|---|---|---|---|
| # | 313 | 366 | 522 | 744 | 958 | 1189 | 1297 | 1462 |
| SampleID | RB13 | DK06 | TR02 | NB16 | FR20 | CS05 | RL08 | AN02 |
| Region1 | | | | | | | | |
| Region2 | | | | Neg | Neg | Neg | Neg | Neg |
| Region3 | Neg | | | | | | | |
| Region4 | | | | | | | | |
| Region5 | | | | | | | | |
| Region6 | | Neg | Neg | | | | Neg | |
| Region7 | Neg | Neg | Neg | Neg | Neg | | Neg | Neg |
| Region8 | | Neg | | | | | | |
| Region9 | Neg | Neg | Neg | Neg | Neg | Neg | Neg | Neg |
| Region10 | | | | | | | | |

**Figure S9.** Estimation of relatedness in 44 resequenced genomes. The kinship coefficient is plotted against the proportion of IBS0. The dashed horizontal lines represent thresholds for different relationships, as indicated in the KING manual.

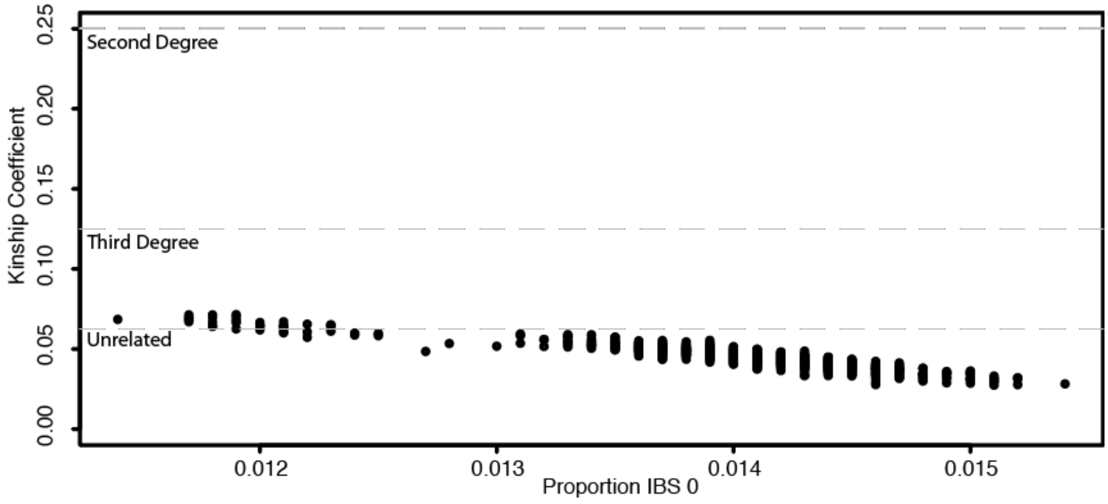

**Figure S10.** MSMC analysis for high coverage genomes. Each line represents a single diploid individual randomly selected from each reef. The changes in effective population size $N_e$ are inferred for all samples jointly and are qualitatively similar to the results generated with the PSMC model presented in the main text.

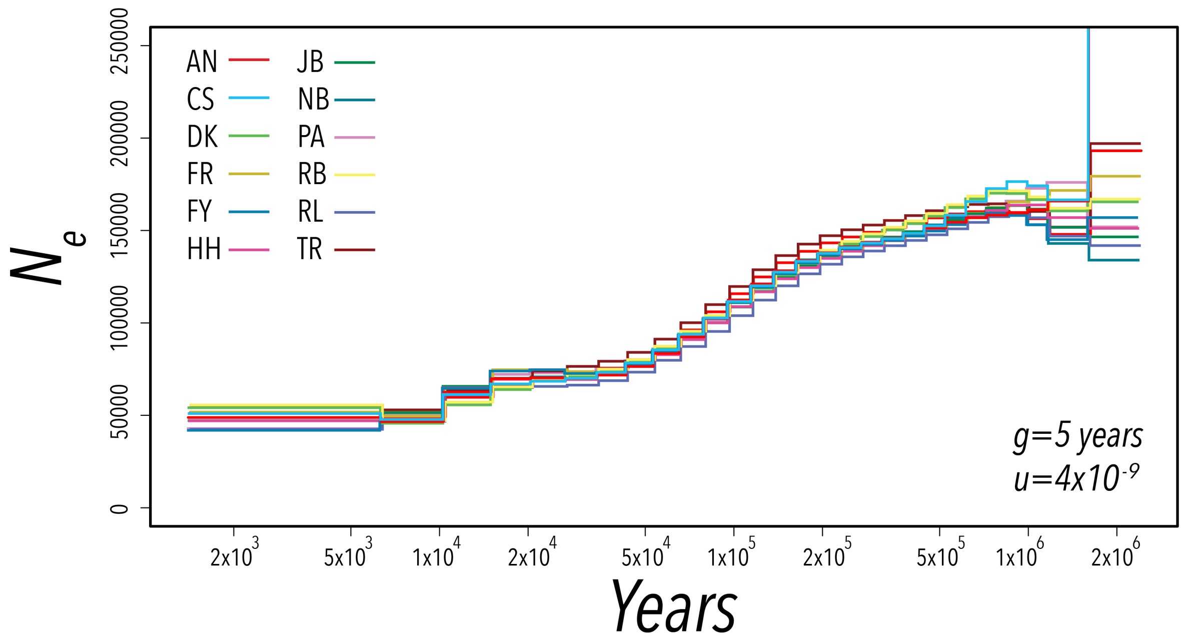

**Figure S11.** Local PCA along the genome, obtained using *lostruct*. The plot on the left shows the first two MDS coordinates used to visualize similarities in relatedness patters among windows, as captured by PCA. Each point representes a 1 kb window. Colors in the corners of the MDS coordinate plot correspond to the colors depicted for windows across the genome. On the right, the midpoint of each window is shown against the first (top) and second (bottom) MDS coordinates. Dotted lines indicate chromosomes. The highest peak on chromosome 7 falls over the *sacsin* gene region.

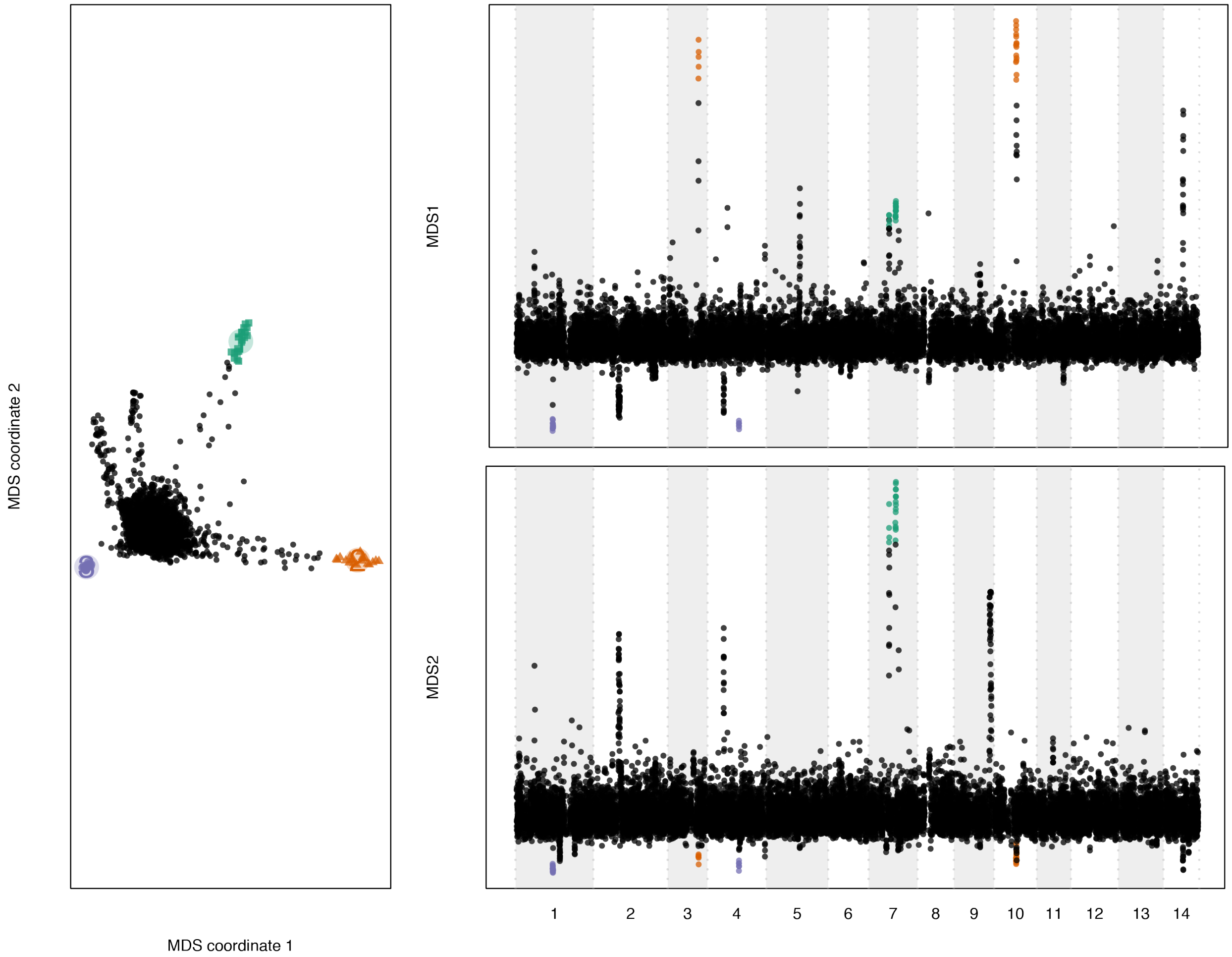

**Figure S12.** The per-base pair coverage distribution in *sacsin* (± 1 kb upstream and downtream of the coding sequence) relative to the average coverage on chromosome 7 for 44 resequenced genomes. The y-axis shows the distribution of coverage represented as a $\log_2$ fold-change.

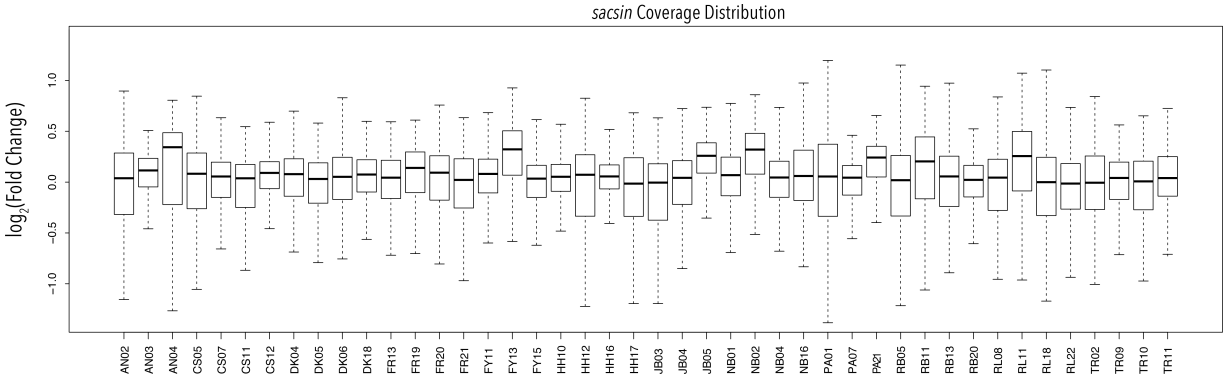

**Figure S13.** Primer design for overlapping amplicons over the *sacsin* region. **(A)** A map showing the location of each segment that was amplified, with the numbers reflecting the primer pair. The coding sequence of *sacsin* is shown in gray. **(B-F)** Gel images showing the results of PCR amplification for all primers pairs in samples PA07, DK06, NB16 and RL08. Primers that successfully amplified for each sample are indicated with a red asterisk above the lane. For regions that successfully amplified, single bands were at the expected length of approximately 2 kb.

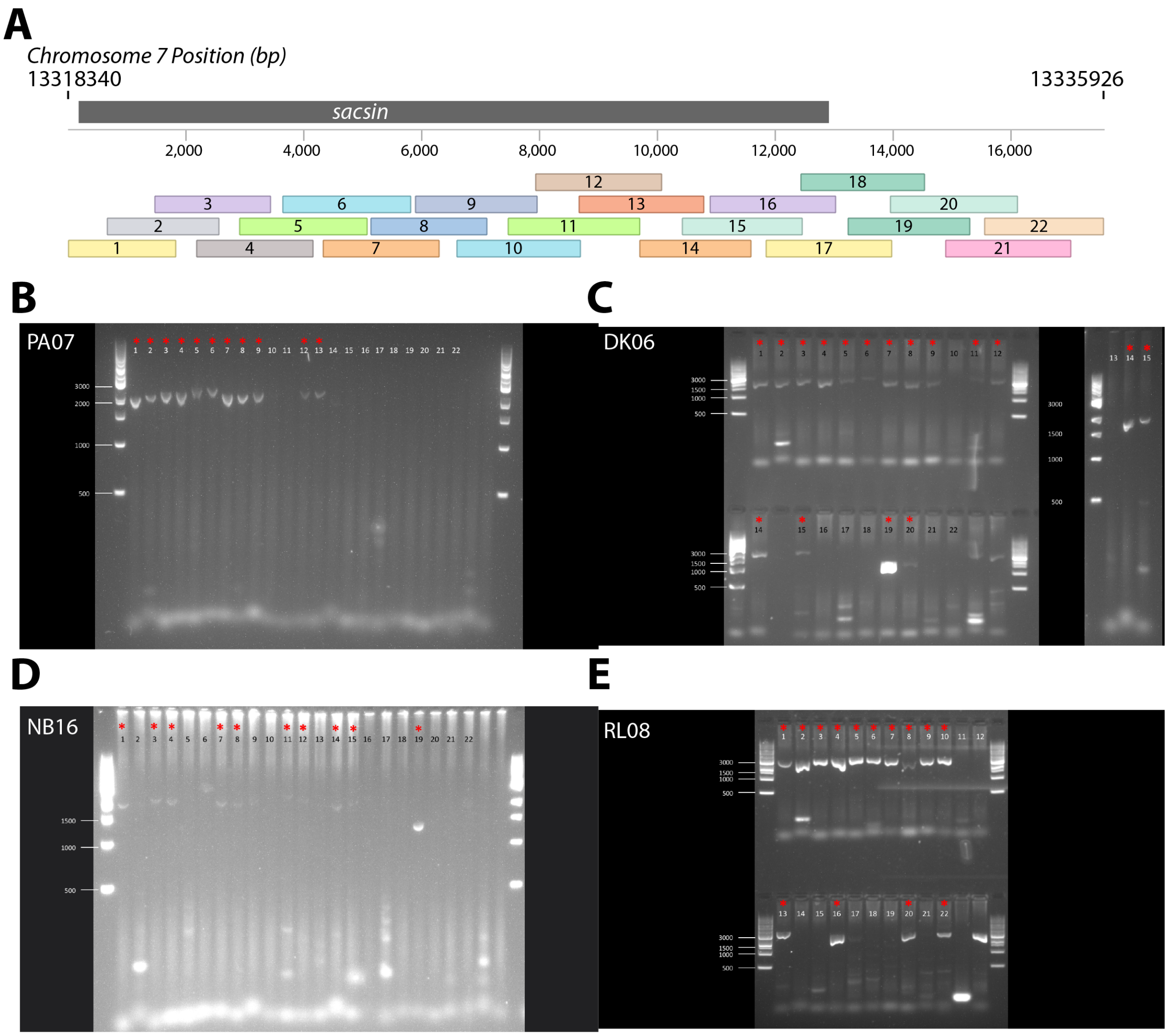

**Figure S14.** Successfully amplified and aligned amplicon sequences for RL08 **(A)** and DK06 **(B)**. Each assembled amplicon sequence for the samples is shown in blue. Small deletions (in red) were observed in both samples downstream of the coding region. Regions of the assembled amplicons that were masked for failing one of our analysis filters are shown in gray.

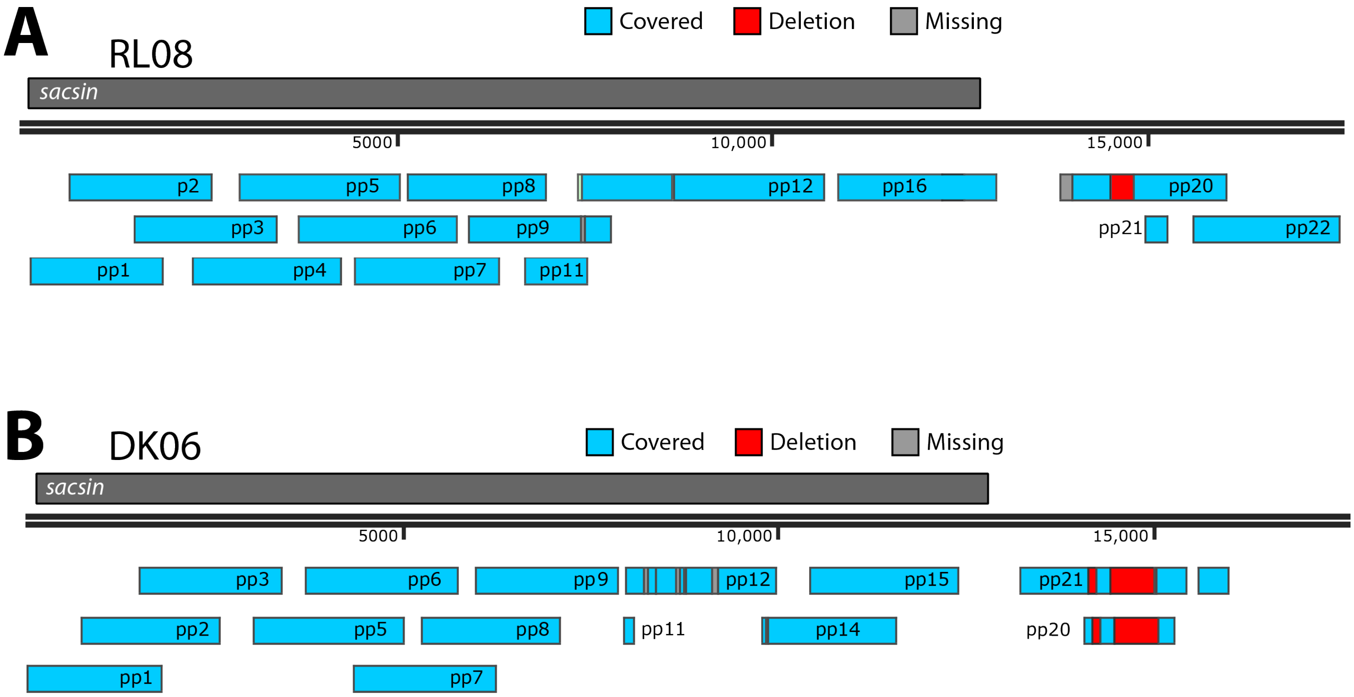

**Figure S15.** The accuracy of imputation approaches. **(A)** The relationship between coverage and minor allele discrepancy (see **5.2**) is shown for two imputation algorithms — Beagle (solid lines) and *loimpute* (dashed lines). For each, minor allele discrepancy was estimated for a range of minor allele frequency (MAF) thresholds, which is represented by the line color. **(B)** The squared correlation coefficient ($r^2$) between imputed and actual genotypes for loimpute (blue) and beagle (green).

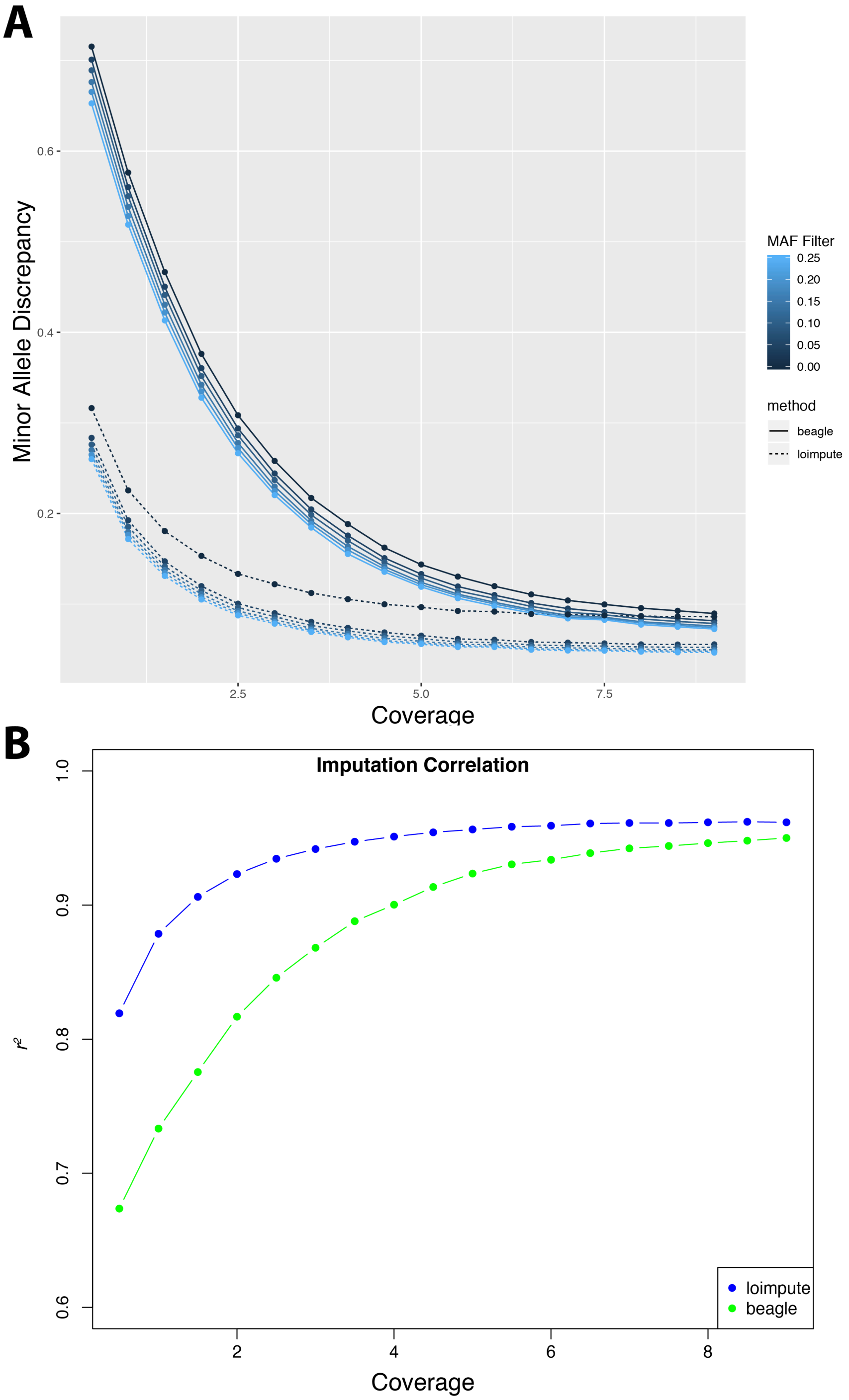

**Figure S16.** Metrics for low-coverage libraries. Histograms showing the proportion of aligned bases that were excluded due to being marked as duplicates, overlapping reads and poor map quality for **(A)** the non-size selected library, **(B)** the size selected library and **(C)** the final combined library. **(D)** The coverage distribution for all samples. Those samples with a mean coverage lower than 0.5× were removed.

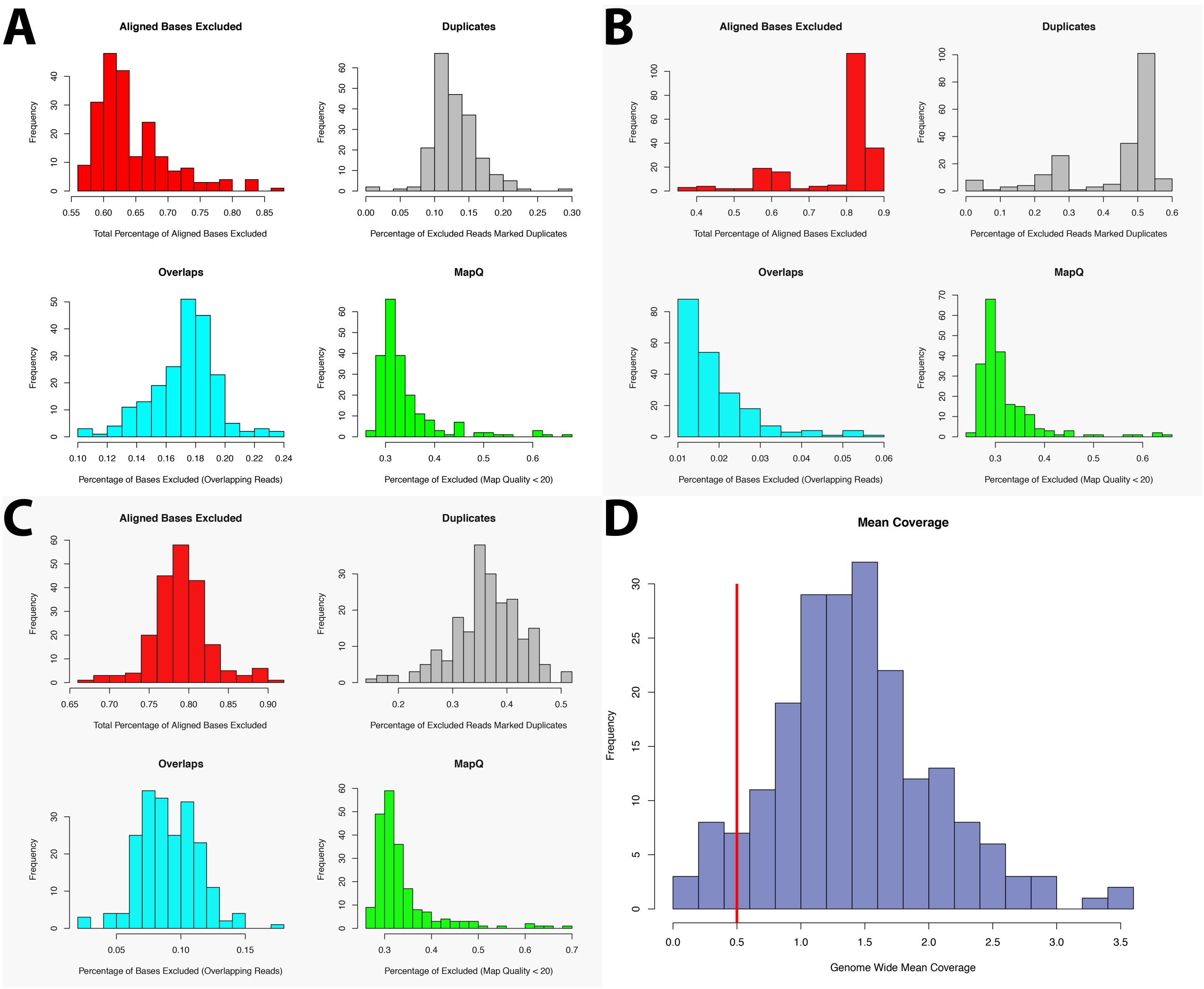

**Figure S17.** Genotype imputation accuracies for 36 samples using *loimpute*. **(A)** Imputation accuracy (measured as $r^2$ between sequenced and imputed genotypes) is plotted against the mean coverage for each sample. Each point represents a sample and solid lines are the best fit, with shaded regions indicating 95% confidence intervals. Imputation accuracy was measured at sites considering three different genotype probability cutoffs, represented by the different colors. Sites with genotype probabilities falling below these thresholds were ignored. **(B)** The distribution of the total proportion of variants excluded across samples for the three different genotype probability cutoffs.

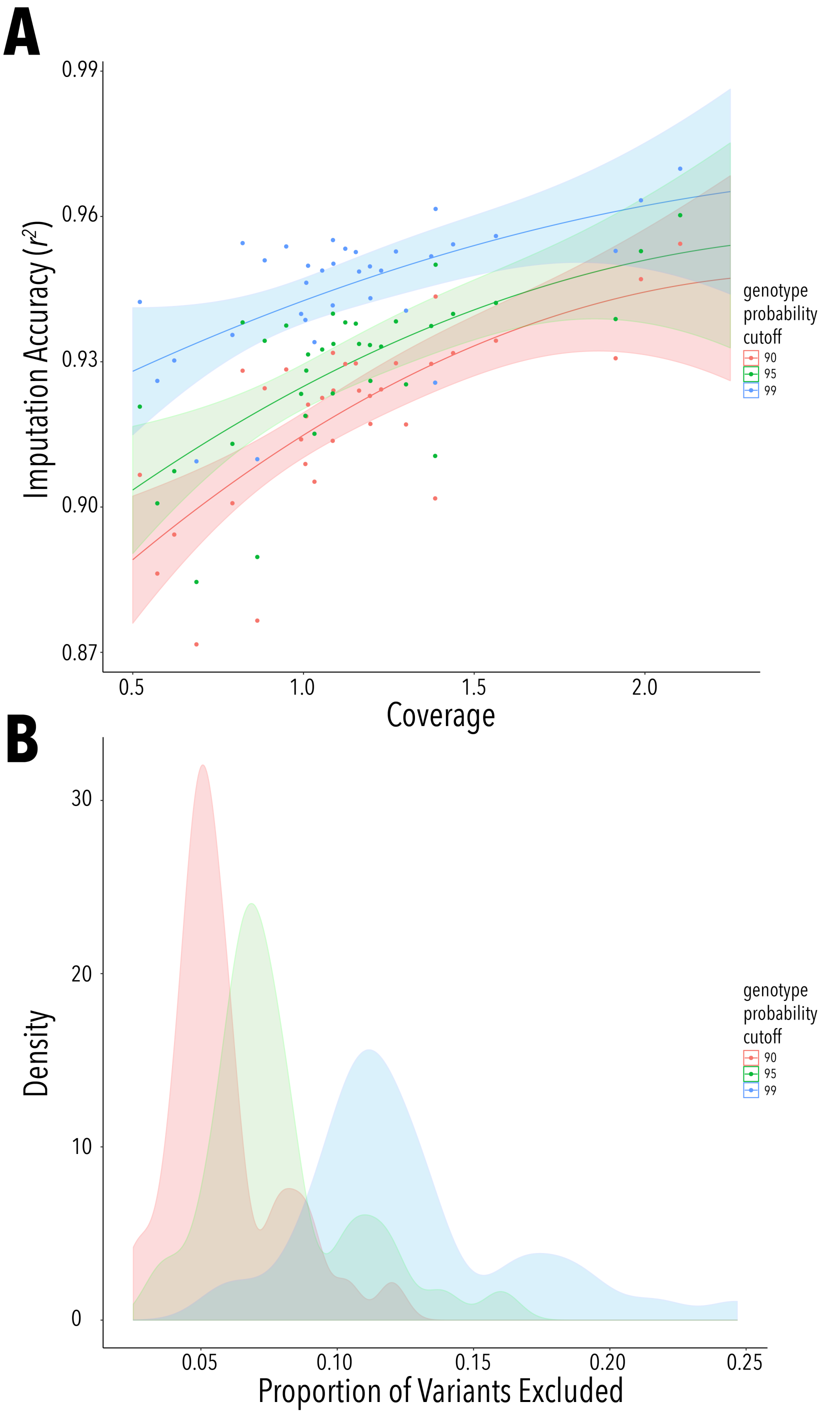

**Figure S18.** Identification of outliers and clonal groups by identity-by-state (IBS) analysis. The dendrogram represents a hierarchical clustering of sequenced corals by genetic distance (1-IBS, shown on the x-axis). Those marked in red have been removed from the dataset as either outliers or clone mates. All samples with a clustering beneath the blue line are retained in the full dataset.

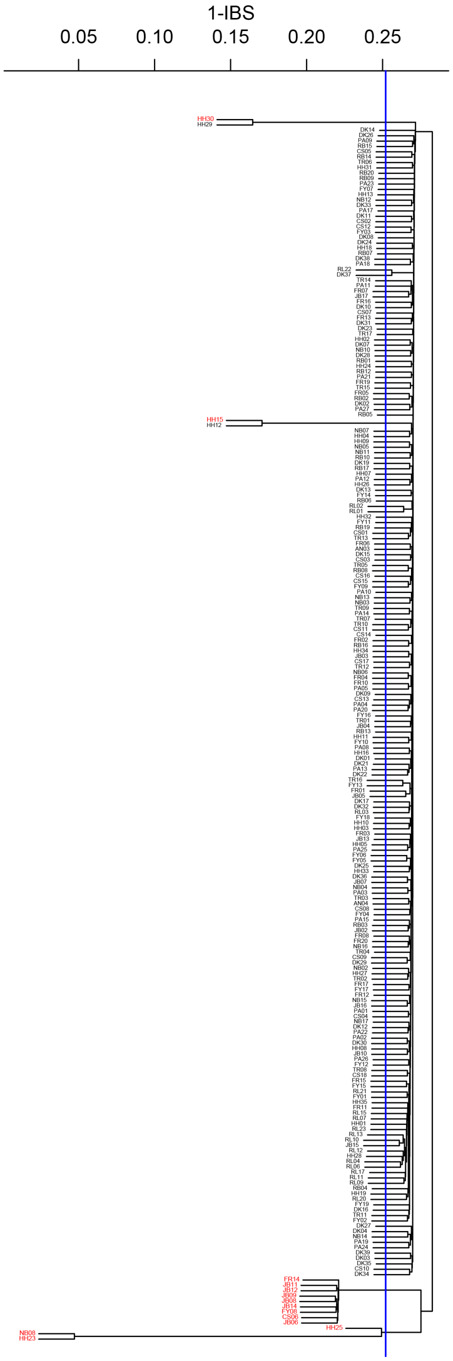

**Figure S19.** PCA for 234 samples, with those identified as outliers by pairwise IBS sharing labeled.

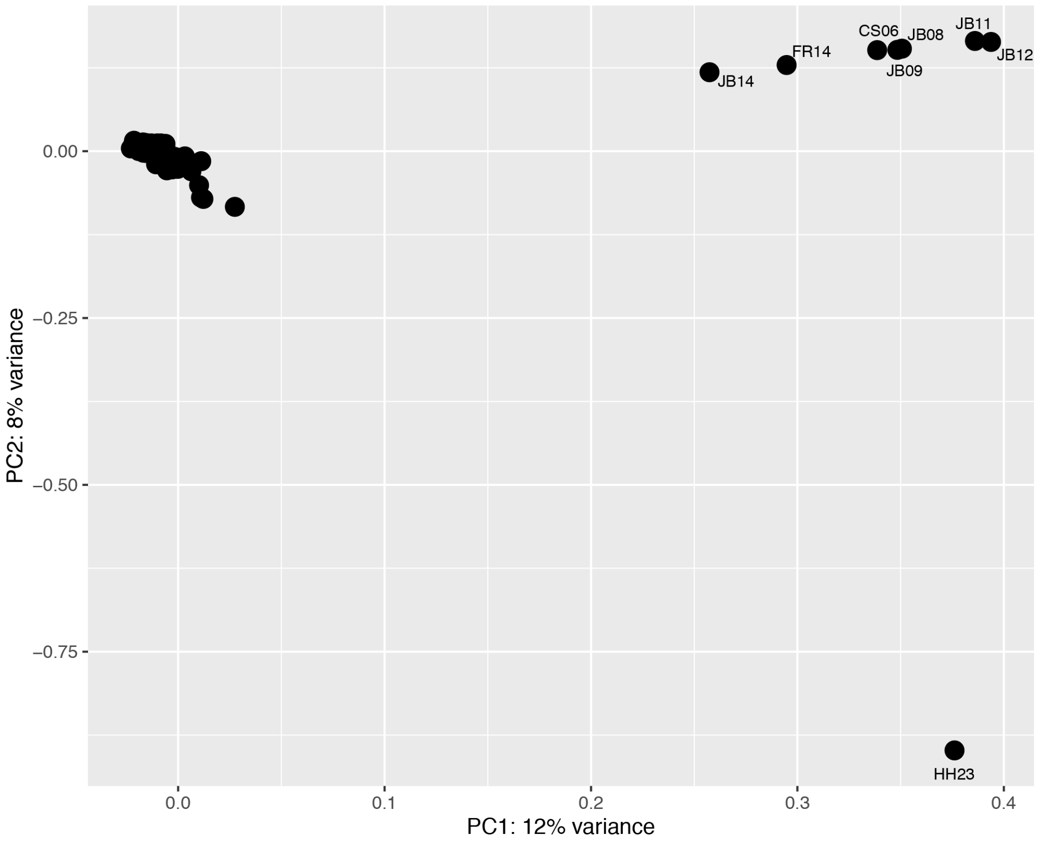

**Figure S20.** Re-analysis in the full set of 213 samples: **(A)** PCA. **(B)** relative effective migration rates. **(C)** Nucleotide diversity levels in 1 kb windows. Points in red represent windows in the top 0.001% genome-wide, and again fall over a single gene *sacsin*. See Figure 4A for the analogous analysis in the 44 high-coverage samples

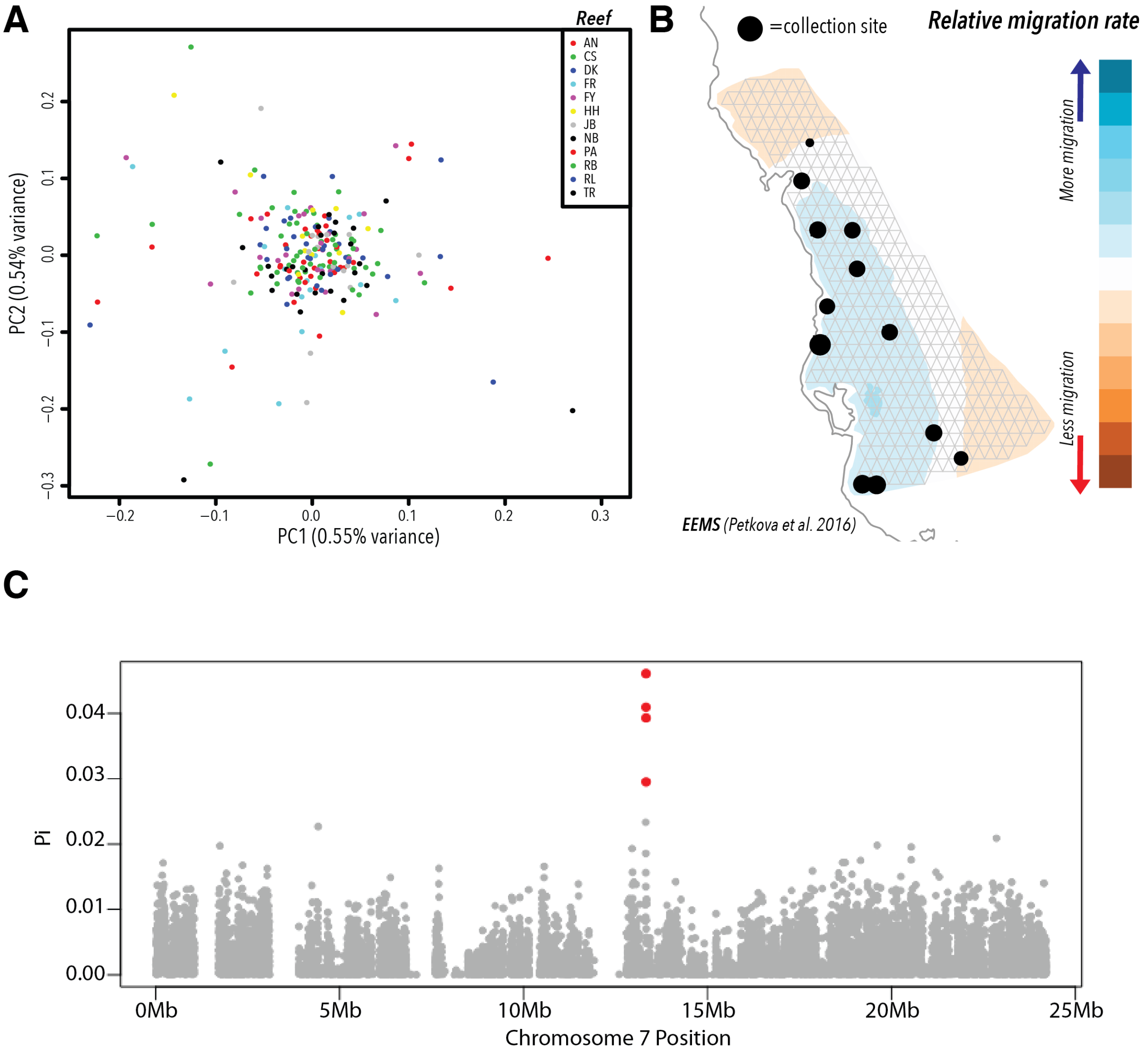

**Figure S21.** Relationship between pairs of phenotype measurements. The Spearman's correlation coefficient for each comparison is shown in the upper-left corner. Visual score vs. total chlorophyll content is shown at the top, visual score vs. symbiont cell density is in the center and total chlorophyll content vs. symbiont cell density is shown at the bottom. For the top two plots, blue lines represent the mean value for that measurement across each visual score increment. In the bottom plot, the blue line represents the line of best of fit.

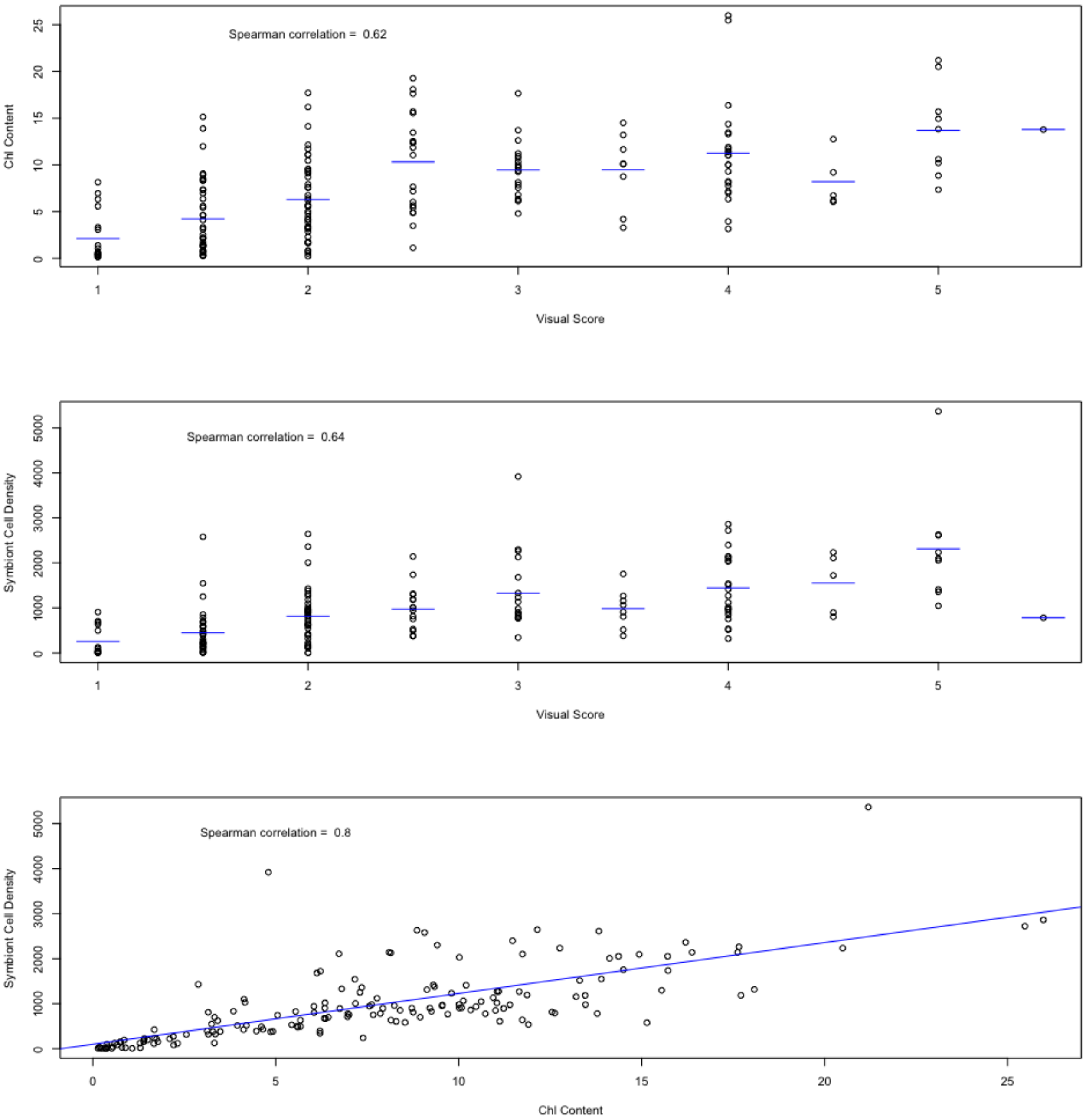

**Figure S22.** Relationship between the abundance of symbiont reads $a_{Symb}$ and measures of bleaching. **(A)** The abundance of symbiont reads $a_{Symb}$ on a $\log_{10}$ scale is on the x-axis against total chlorophyll content standardized by host protein levels on a $\log_{10}$ scale shown on the y-axis. **(B)** The abundance of symbiont reads $a_{Symb}$ on a $\log_{10}$ scale is on the x-axis against standardized symbiont cell density on a $\log_{10}$ scale shown on the y-axis. Squared correlation coefficients are in the upper-left corner of each plot.

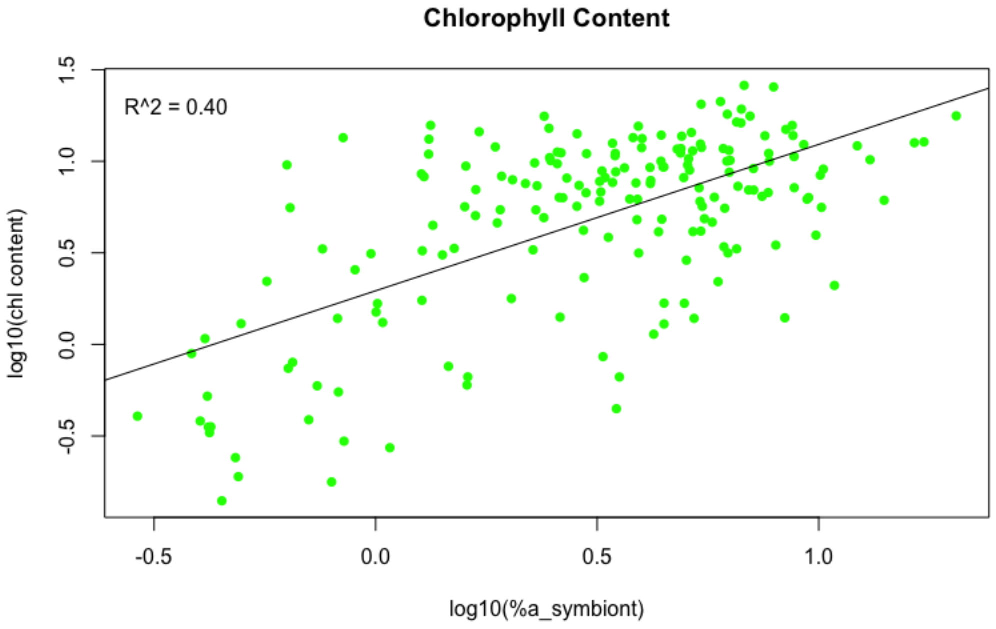

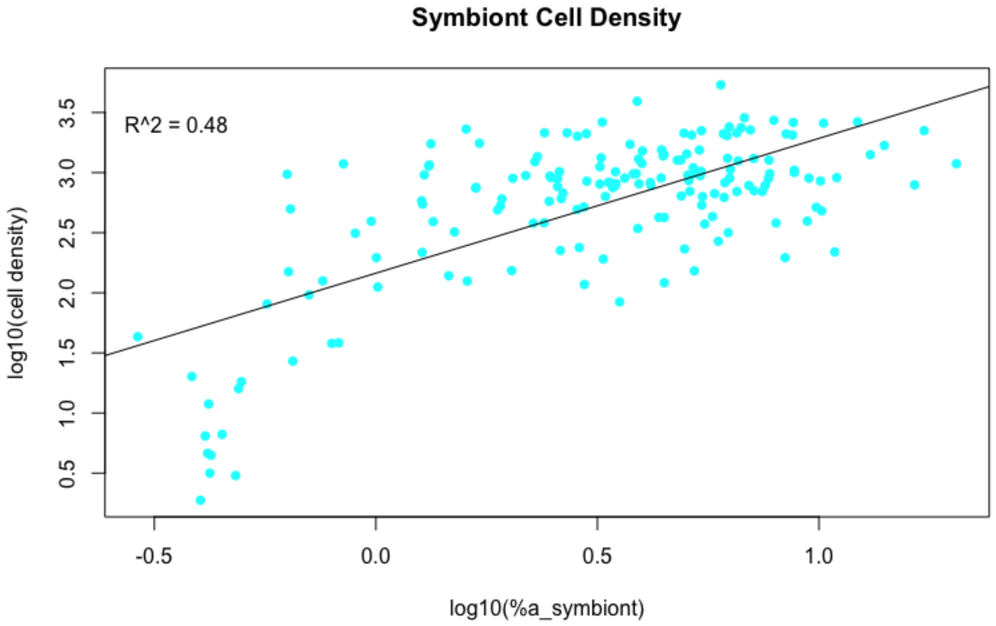

**Figure S23.** Manhattan plots for a GWAS performed using an LMM for total chlorophyll content **(A)** and symbiont cell density **(B)**. Both phenotypes were quantile normalized. Red lines represent the thresholds for genome-wide significance obtained from permutation tests (see **7.6**).

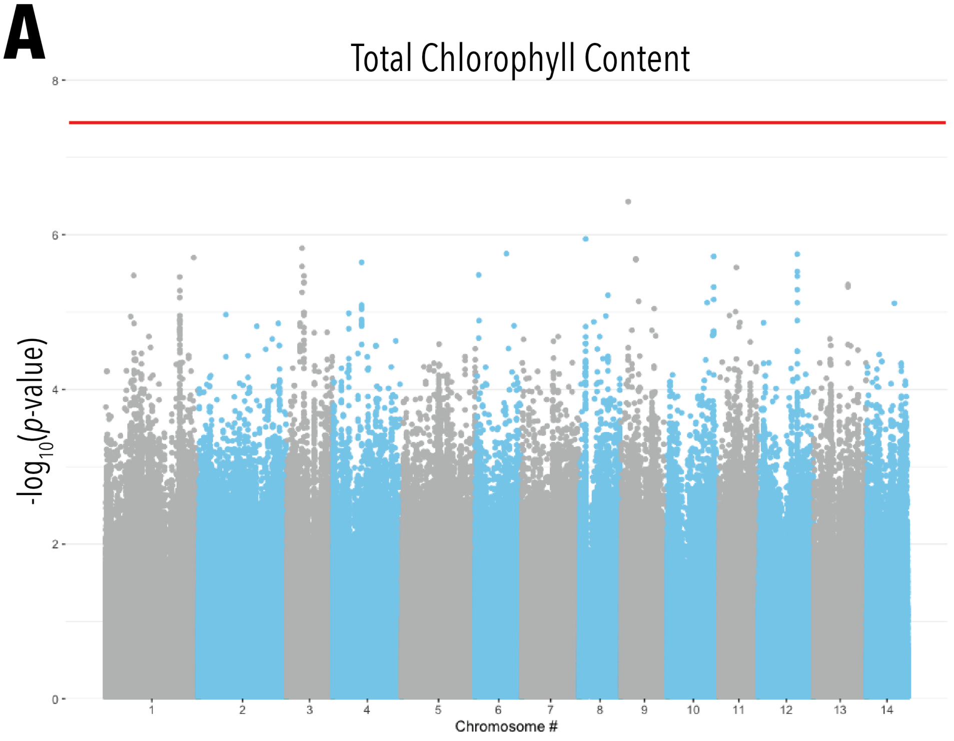

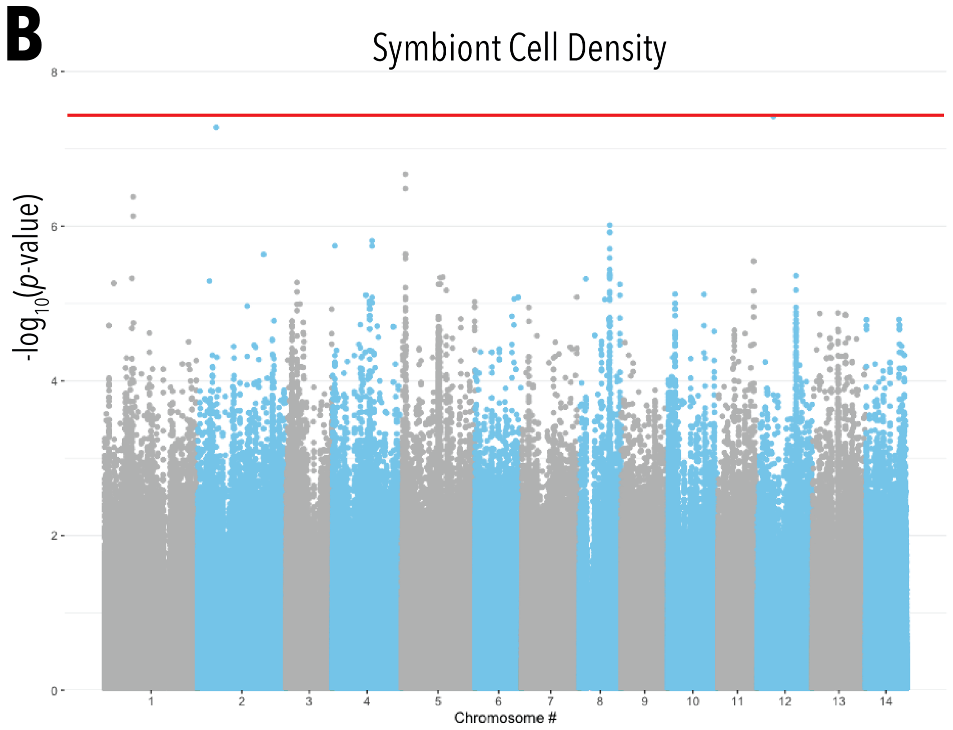

**Figure S24.** Comparison of $p$-values between a GWAS performed using a linear regression implemented in PLINK (with the first two genetic PCs included as covariates) and a GWAS performed using an LMM implemented in GEMMA.

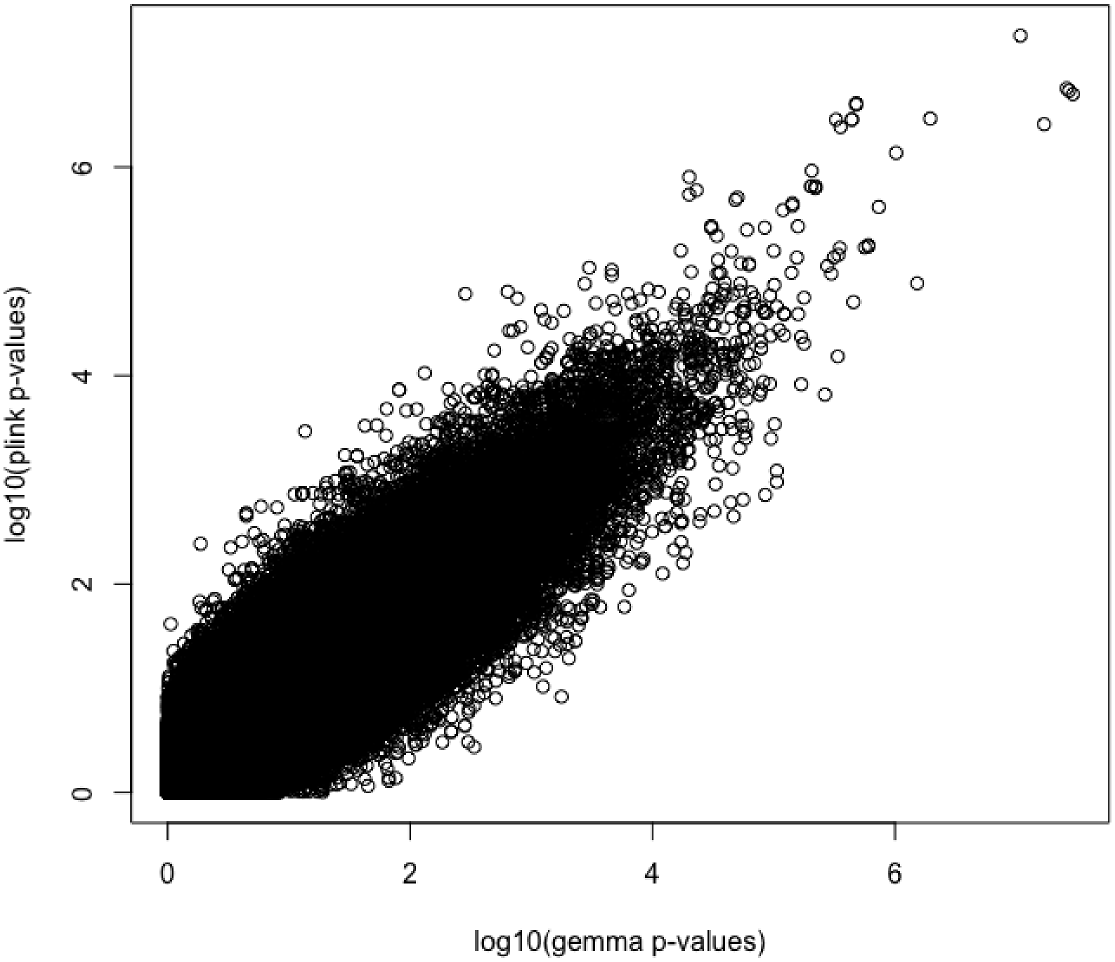

**Figure S25.** Comparison of effect sizes $\beta$ estimated at SNPs with $p < 10^{-4}$ in a GWAS using only individuals from northern reefs (left) and southern reefs (right). In each case, the correlation between effects sizes at SNPs in individuals from the two groupings are highly correlated, approximately 0.90 (with $r^2$).

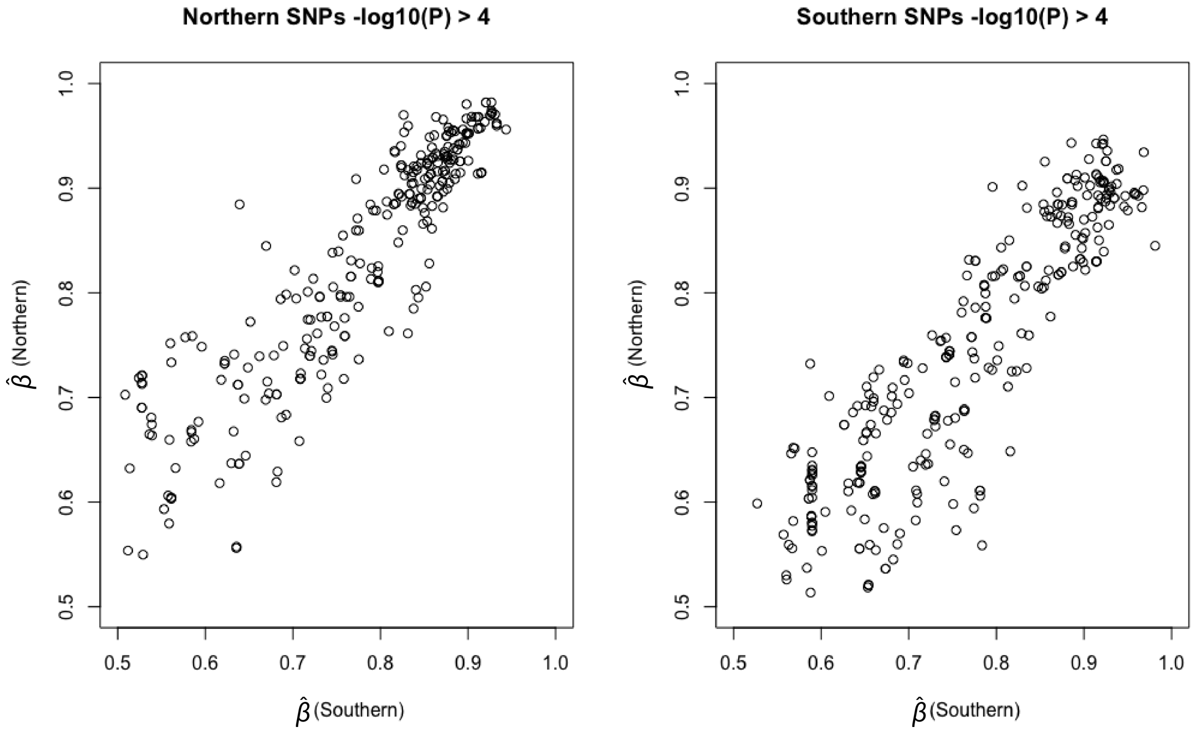

**Figure S26.** GWAS results when removing the symbiont type as a covariate. **(A)** A Manhattan plot for GWAS for quantile-normalized visual socre when removing the dominant symbiont type as a covariate. All other covariates remain the same as the model presented in the main text. **(B)** The relationship between estimated effects sizes ($\beta$) between GWAS models including or removing the symbiont type as covariate.

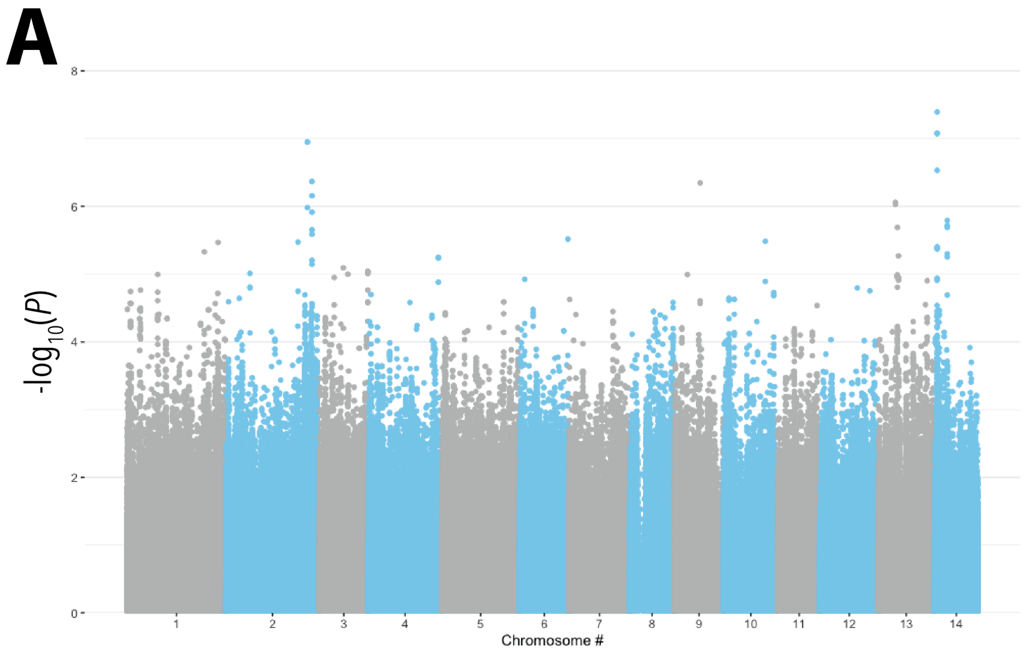

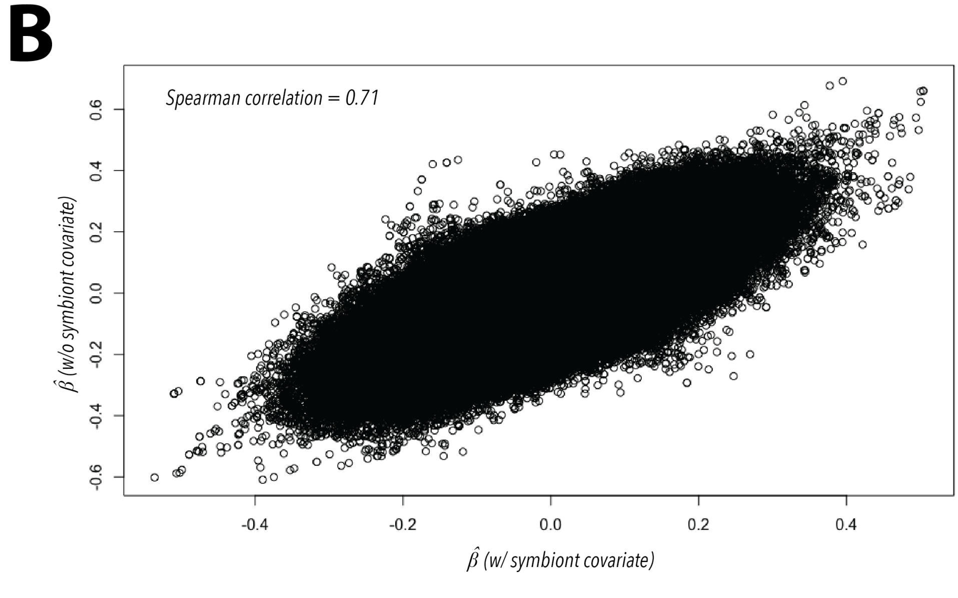

**Figure S27.** Manhattan plot for GWAS for the dominant symbiont type (see **7.1.2**) as the phenotype. Only one SNP has a $p$−value $< 10^{-4}$

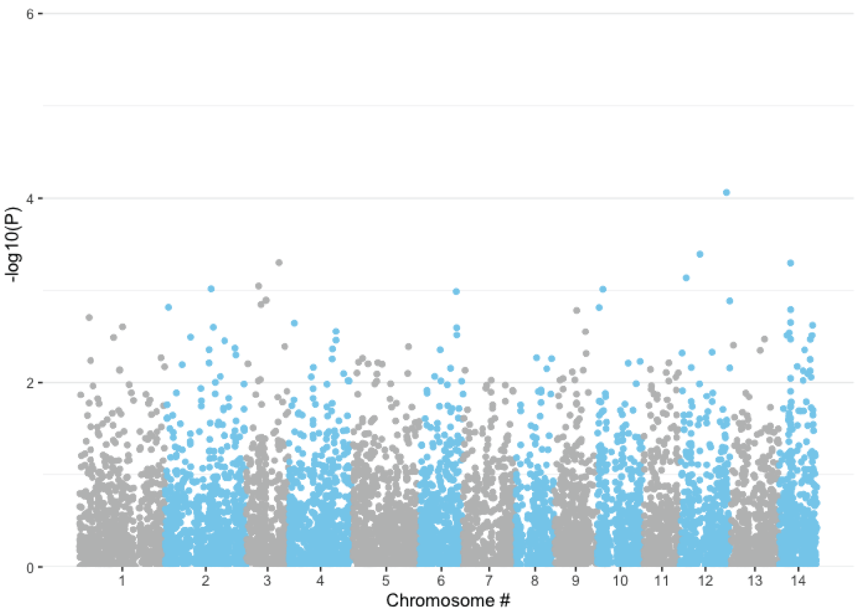

**Figure S28.** Manhattan plot for GWAS of quantile-normalized visual score using BIMBAM[59]. Bayes Factors on a $\log_{10}$ scale are shown on the y-axis.

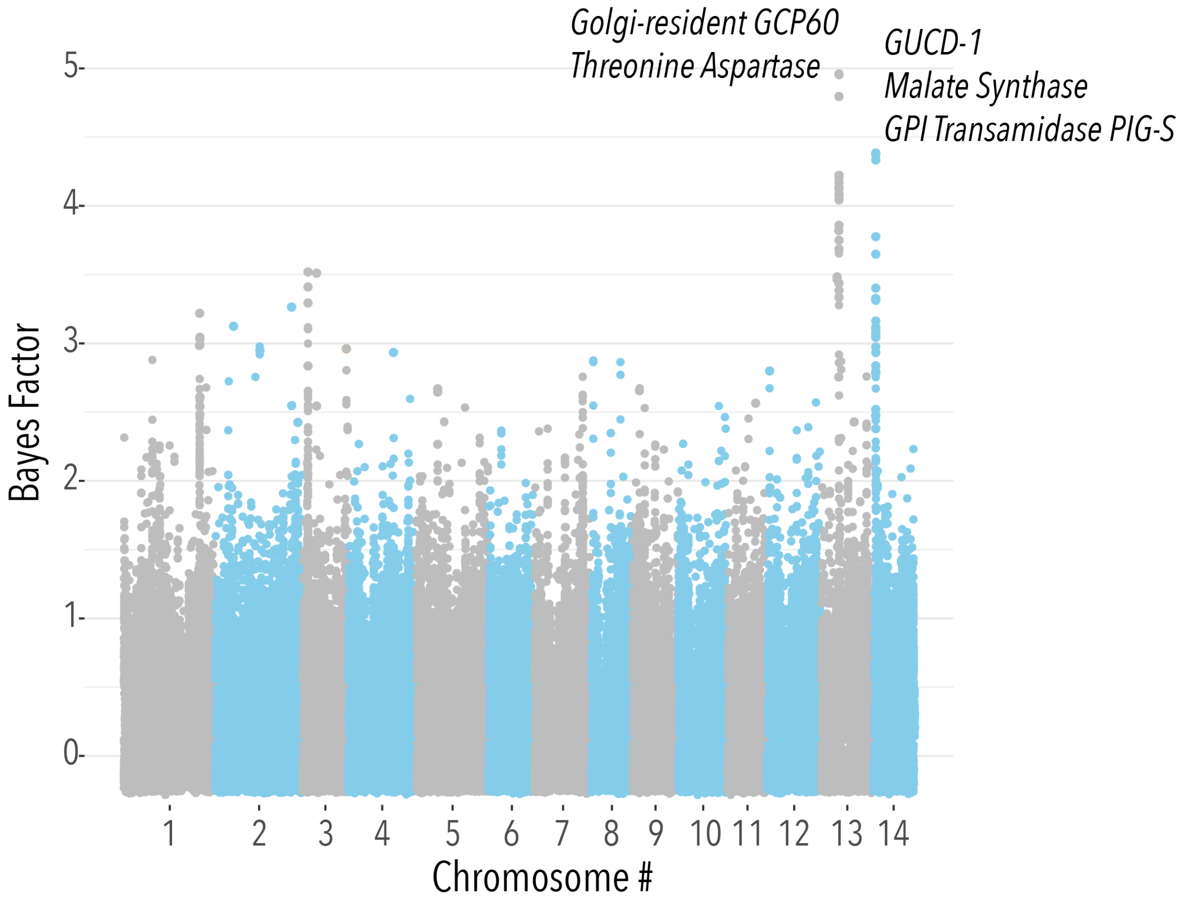

**Figure S29.** Distribution of minimal p-values determined from a permutation test to estimate a threshold of genome-wide significance for the three measures of bleaching (see **7.6**). The $95^{th}$ percentile for each is indicated with a vertical line, and was used as the genome-wide significance threshold.

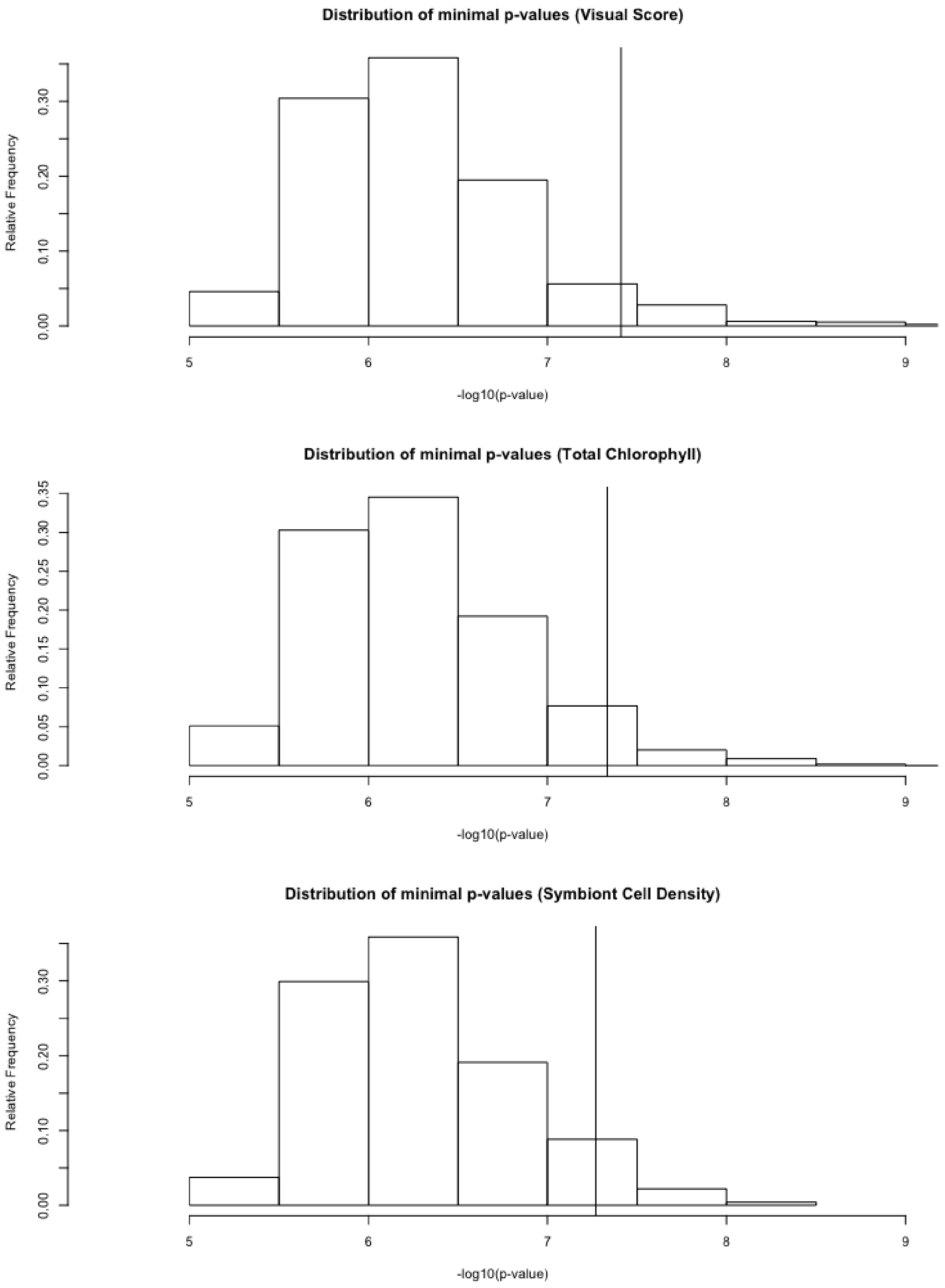

**Figure S30.** Prediction accuracy of the PGS, measured as $R^2$ between the PGS and actual phenotype values in the test set. The dots show the mean and lines show the standard deviation from 100 jackknife partitions of training and test sets for SNPs with a $p$-value less than the cutoff shown on the x-axis. $R^2$ is maximized for a $p$-value cutoff of $10^{-5}$.

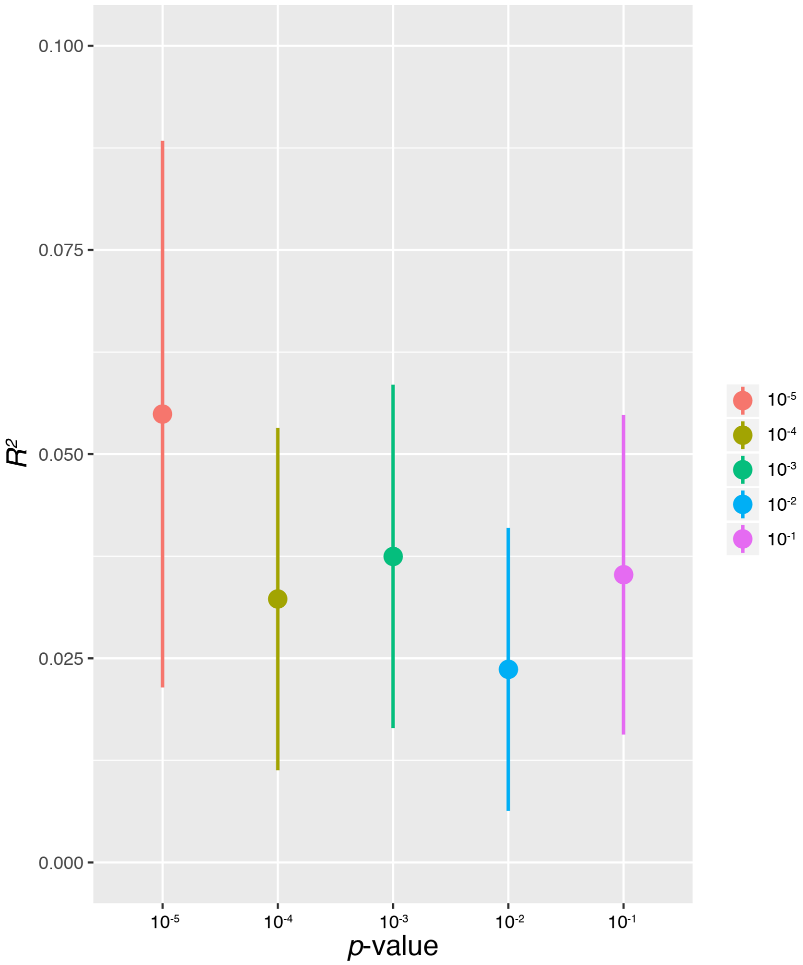

**Figure S31.** Prediction accuracy, measured as $R^2$ in the test set in a jackknife cross validation procedure. **(A)** A training-test split of 90/10%. **(B)** A training-test split of 80/20%. In both cases, the same covariates are used as the model described and presented in the main text.

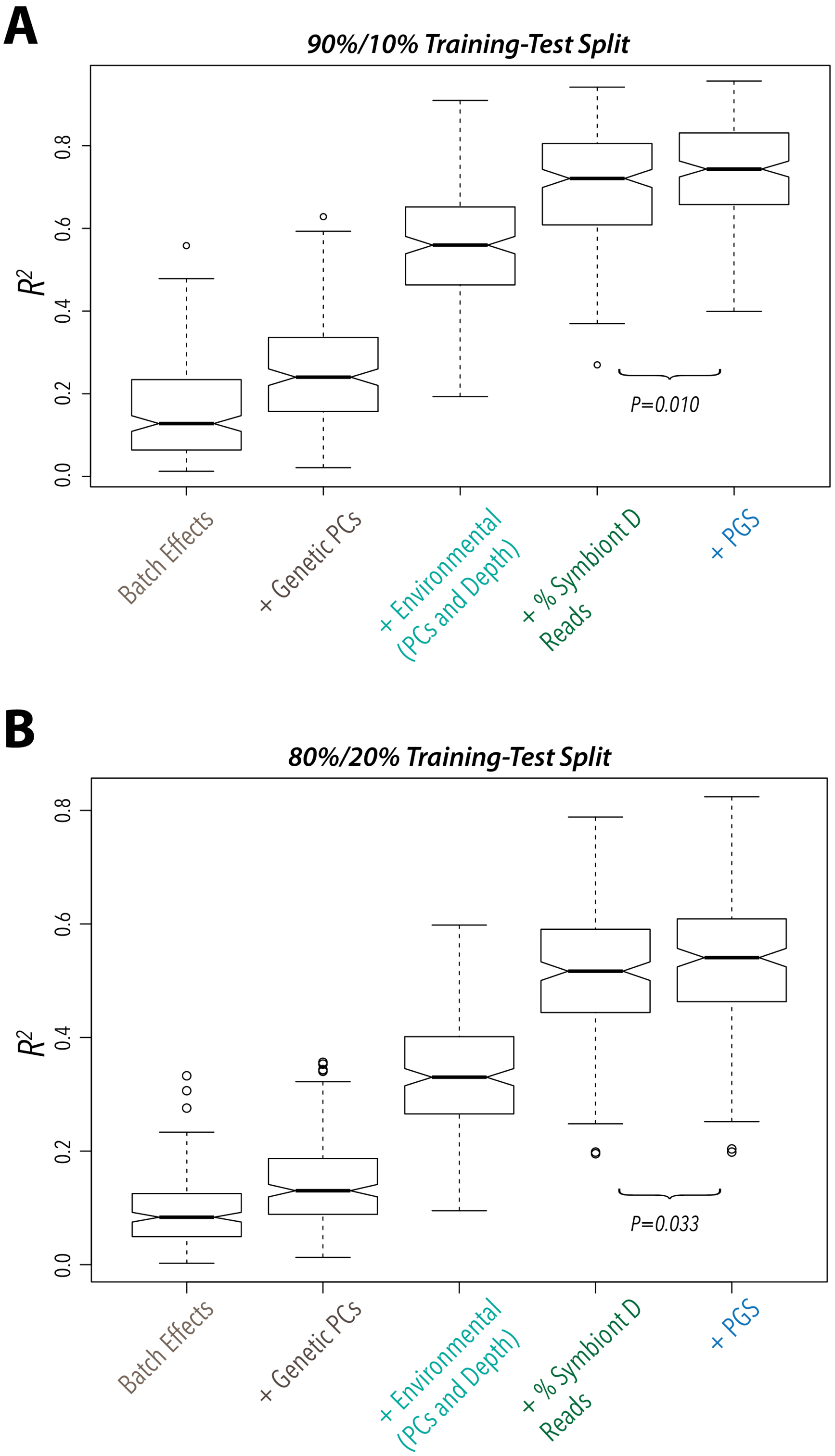

**Figure S32.** Prediction accuracy, measured as $R^2$ in the test set in a jackknife cross validation procedure. **(A)** The environmental PCs include the top six PCs. **(B)** The environmental PCs include the top eight PCs.

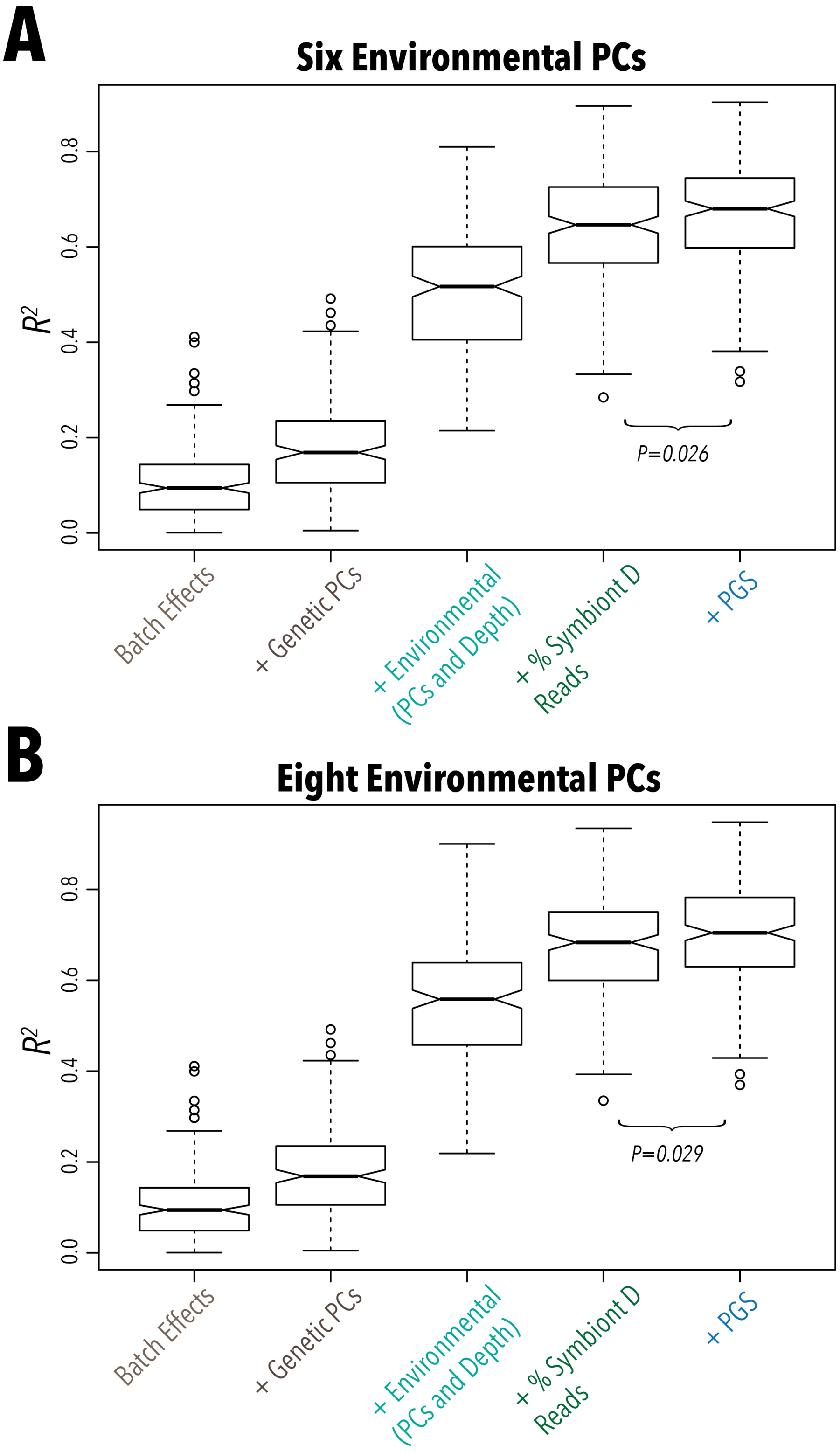

### 9 Supplemental Tables

**Table S1.** Comparison of genome assemblies. Note that the BUSCO results here refer to the core metazoan set, which is reported in Ying *et al.* (2019)[23].

| Statistic | Fuller *et al.* assembly | Ying *et al.* assembly |
|---|---|---|
| Length (Mb) | 475.4 | 386.60 |
| GC-content | 39.06 | 38.85 |
| Repeat % | 42.00 | 34.55 |
| Scaffold # | 854 | 3,876 |
| Longest Scaffold | 39.4 Mb | 3.8Mb |
| N50 (Mb) | 19.8 | 0.49 |
| Annotated Genes | 28,186 | 26,615 |
| BUSCO % | 91.8 | 92.9 |

**Table S2.** Collection sites, locations and dates

| Site | Abbreviation | Latitude | Longitude | Date |
|---|---|---|---|---|
| Arlington Reef | AN | 16°42'16.75"S | 146°2'50.10"E | 24/3/17 |
| Fitzroy Island | FY | 16°55'31.09.75"S | 145°59'20.79"E | 25/3/17 |
| Coates Reef | CS | 17°11'19.79"S | 146°22'11.36"E | 22/3/17 |
| Russell Island | RL | 17°13'24.97"S | 146°5'23.07"E | 23/3/17 |
| Feather Reef | FR | 17°31'43.00"S | 146°23'12.86"E | 21/3/17 |
| North Barnard Islands | NB | 17°44'23.92"S | 146°9'22.48"E | 20/3/17 |
| Taylor Reef | TR | 17°48'52.87"S | 146°34'4.25"E | 18/3/17 |
| Dunk Island | DK | 17°57'20.70"S | 146°8'53.86"E | 17/3/17 |
| Rib Reef | RB | 18°28'51.80"S | 146°52'16.62"E | 16/3/17 |
| Pandora Reef | PA | 18°48'49.01"S | 146°25'56.46"E | 15/3/17 |
| John Brewer Reef | JB | 18°38'23.85"S | 147°2'29.22"E | 1/4/17 |
| Havannah Island | HH | 18°50'22.08"S | 146°32'4.35"E | 14/3/17 |

**Table S3.** Samples used for high coverage sequencing and coverage statistics after alignment. The sequencing batch is the same as in Figure S6. Those samples marked with an asterisk were removed as potential outliers.

| Sample ID | Reef | Batch | Mean Coverage per bp | Standard Deviation per bp |
|---|---|---|---|---|
| AN02 | Arlington | 1 | 29.04 | 22.88 |
| AN03 | Arlington | 2 | 152.49 | 98.93 |
| AN04 | Arlington | 2 | 135.59 | 92.03 |
| CS05 | Coates | 1 | 28.82 | 22.49 |
| CS07 | Coates | 2 | 114.66 | 79.05 |
| CS11 | Coates | 2 | 133.29 | 89.26 |
| CS12 | Coates | 2 | 141.45 | 92.85 |
| DK04 | Dunk | 2 | 111.67 | 76.89 |
| DK05 | Dunk | 2 | 120.88 | 83.22 |
| DK06 | Dunk | 1 | 25.49 | 20.36 |
| DK18 | Dunk | 2 | 125.13 | 84.61 |
| FR13 | Feather | 2 | 128.20 | 87.02 |
| FR19 | Feather | 2 | 137.89 | 91.71 |
| FR20 | Feather | 1 | 35.49 | 27.28 |
| FR21 | Feather | 2 | 132.52 | 89.67 |
| FY08* | Fitzroy | 1 | 21.30 | 18.30 |
| FY11 | Fitzroy | 2 | 154.04 | 101.25 |
| FY13 | Fitzroy | 2 | 131.75 | 90.48 |
| FY15 | Fitzroy | 2 | 164.76 | 105.82 |
| HH10 | Havannah | 2 | 145.29 | 94.58 |
| HH12 | Havannah | 2 | 108.56 | 79.46 |
| HH16 | Havannah | 2 | 111.92 | 76.53 |
| HH17 | Havannah | 2 | 133.78 | 91.10 |
| HH22* | Havannah | 1 | 20.15 | 29.13 |
| JB03 | John Brewer | 2 | 125.99 | 85.93 |
| JB04 | John Brewer | 2 | 142.15 | 96.16 |
| JB05 | John Brewer | 2 | 142.83 | 93.89 |
| JB06* | John Brewer | 1 | 30.97 | 25.16 |
| NB01 | North Barnard | 2 | 103.13 | 72.02 |
| NB02 | North Barnard | 2 | 120.64 | 82.78 |
| NB04 | North Barnard | 2 | 103.65 | 73.23 |
| NB16 | North Barnard | 1 | 24.67 | 19.57 |
| PA01 | Pandora | 2 | 114.41 | 85.44 |
| PA06* | Pandora | 1 | 25.23 | 33.87 |
| PA07 | Pandora | 2 | 159.16 | 103.08 |
| PA21 | Pandora | 2 | 123.54 | 83.85 |
| RB05 | Rib | 2 | 112.36 | 80.98 |
| RB11 | Rib | 2 | 128.75 | 87.71 |
| RB13 | Rib | 1 | 25.07 | 20.16 |
| RB20 | Rib | 2 | 144.52 | 95.40 |
| RL08 | Russell | 1 | 28.28 | 22.76 |
| RL11 | Russell | 2 | 91.24 | 68.43 |
| RL18 | Russell | 1 | 102.25 | 76.37 |
| RL22 | Russell | 1 | 129.78 | 88.62 |
| TR02 | Taylor | 2 | 28.32 | 22.02 |
| TR09 | Taylor | 1 | 120.05 | 82.62 |
| TR10 | Taylor | 1 | 125.41 | 85.61 |
| TR11 | Taylor | 1 | 109.52 | 75.93 |

**Table S4.** Regions used for PCR amplification and MiSeq sequencing to validate SNP calls. Each region was randomly selected, with the primer pairs used to amplify each shown below. Primers were designed to have a minimum annealing temperature of 51°C.

| Region | Scaffold | Start | End | Size | Forward Primer | Reverse Primer |
|---|---|---|---|---|---|---|
| 1 | chr12 | 22825856 | 22828200 | 2344 | TGCAACACTGACCGATGGTT | ACGTGGTTGGCAGAATAGCA |
| 2 | chr1 | 24445742 | 24447829 | 2087 | TCGAAATCACGCCACAGGAA | TCTCTGGGCATTTCCCGTTC |
| 3 | chr7 | 14707604 | 14710055 | 2451 | GGAGAGCGCAGTAACACAGT | TAGTGAAAAGCCCGCTCTGG |
| 4 | chr6 | 1193840 | 1196073 | 2233 | ATACGCTCATTTTTCGCCGC | CGTCGAGAAATCCTCCTCGG |
| 5 | chr4 | 26088722 | 26090722 | 2000 | ACTGCACGACTCTTGCTTCA | CGGATTCGCCATCTCTCCAA |
| 6 | Sc0000066 | 686480 | 688687 | 2207 | GTACCTTTGTCGCCTGACCA | ACGGTTGGCTCGATCGTATG |
| 7 | chr3 | 16299377 | 16301515 | 2138 | AGTCGGCACTCTGCATGAAA | GCTTGAATTTCACCGGCGTT |
| 8 | chr3 | 1658661 | 1660763 | 2102 | CAGTCTCCGCCTCACTTGTT | GCTCGAGCAGCTTTCTGGTA |
| 9 | Sc0000126 | 24480 | 26617 | 2137 | GGGCGGGTATTCCTTCACAA | AAGTACTGAGCCGCACTTCC |
| 10 | chr1 | 32314547 | 32316572 | 2025 | AACCGCAAAAGGTCTCGTCT | ATGACTCCCGGAGGAAAACG |

**Table S5.** Correlation ($r^2$) and $p$-values for a linear regression of frequencies of the two most common haplotypes in *sacsin* and genetic and environmental PCs.

| Variable | $r^2$ | $p$-value |
|---|---|---|
| Environmental PC1 | 0.004 | 0.355 |
| Environmental PC2 | 0.002 | 0.496 |
| Environmental PC3 | 0.001 | 0.862 |
| Environmental PC4 | 0.002 | 0.841 |
| Environmental PC5 | 0.004 | 0.396 |
| Environmental PC6 | 0.010 | 0.166 |
| Environmental PC7 | 0.001 | 0.975 |
| Environmental PC8 | 0.002 | 0.515 |
| Genetic PC1 | 0.004 | 0.402 |
| Genetic PC2 | 0.007 | 0.266 |

**Table S6.** Regions used for PCR amplification and MiSeq sequencing to validate the *sacsin* region. Primers were designed to have a minimum annealing temperature of 51°C.

| Region | Start | End | Size | Forward Primer | Reverse Primer |
|---|---|---|---|---|---|
| 1 | 8 | 1826 | 1818 | TCACTGATGTTGACCACACCTGGA | GGGACAAAGGCCAGATCTCTCA |
| 2 | 666 | 2559 | 1893 | TGCCCTACCTTACGAGCGGTGG | GGGAGCCGAAAAGCCATCTCCA |
| 3 | 1474 | 3435 | 1961 | AGCTCGCTCCCACTTTTTGAGGA | TCTGCTAACCAACATGGGGACA |
| 4 | 2183 | 4156 | 1973 | CGCCTGACAGGTTTCCCAAGAGC | TTGCTTCACCAGTGCCCATGGA |
| 5 | 2912 | 4770 | 1858 | ATTGTGAACGCGAGTGGCTGCA | CGAGTCAAGCACTTCAAGCG |
| 6 | 3645 | 5522 | 1877 | CCCCCTTCGAACCAAGGAACAAGC | CTCTTGGGTGCCATGCGGGAAC |
| 7 | 4330 | 6294 | 1964 | AATGGCGCGTTTGCAGTAGCCT | ACGCCAACTGCTTTCAAAAAGTTCC |
| 8 | 5140 | 7105 | 1965 | TGGCTGGAAATTGCAGAAAGGTCGG | ACCCCAGCTGCTGAACAATGAACC |
| 9 | 5896 | 7958 | 2062 | CAGTCGCTTCATGTCGCTGTTC | GGCGTATTGCTGACATCAGTTGGT |
| 10 | 6599 | 8692 | 2093 | ACGCCAACCCTCAAGAACCAGG | TCGTGCCAGAGACGCAAGCTG |
| 11 | 7469 | 9699 | 2230 | CAGAGCCTCTGAACAGCCCCGA | GCGCGGGAGAAGATGTGCTACT |
| 12 | 7935 | 10071 | 2136 | ACCAACTGATGTCAGCAATACGCC | ACGCCCCAAACTCGAAAGGCAC |
| 13 | 8672 | 10785 | 2113 | CAGCTTGCGTCTCTGGCACGA | TGCCACTTCCGTCTGGCTTCCT |
| 14 | 9702 | 11586 | 1884 | GGCAGACCCGAGACCTCACAGT | CGCTTCGGAGCCTGTTCGTTGT |
| 15 | 10423 | 12457 | 2034 | GGACCGGATTGCAGAGCACGAC | TCCGGCCAAGTGACGCAATGAC |
| 16 | 10916 | 13023 | 2107 | TAGCCGAGGCATGTGAAGGGCT | GCGAGGGGATGAAGGCAAGAGG |
| 17 | 11845 | 13974 | 2129 | TCCTACGAGTGGGCGTGCTTCA | TGCAAATTGTCCGCAGCCCCAA |
| 18 | 12436 | 14531 | 2095 | GTCATTGCGTCACTTGGCCGGA | ACGTCAGCTTTCCAGTTCCACT |
| 19 | 13236 | 15300 | 2064 | ACAGCGACGAACCAGTGACAGT | ACCAAGTGCGAGTTCGTTGCAA |
| 20 | 13954 | 16109 | 2155 | TGGGGCTGCGGACAATTTGCAA | CGGCTGACCATAGCAAGCCTCG |
| 21 | 14892 | 17013 | 2121 | AGGTCAGTGGAGTCCTGTTGCA | CCGTCCACCAGTTCACTCAACCG |
| 22 | 15552 | 17569 | 2017 | AGGACCACACAAGGGACGTTCT | TCGGTCTAGCATGGCAACCCGA |

**Table S7.** Genotype concordance matrix of original SNP calls and MiSeq sequenced amplicons in *sacsin* for DK06 (left) and RL08 (right). In both, the original genotype calls are on the top and the MiSeq validation calls are on the side.

| **DK06** | 0/0 | 0/1 | 1/1 |
|---|---|---|---|
| 0/0 | 855 | 0 | 0 |
| 0/1 | 0 | 1 | 0 |
| 0/1 | 0 | 0 | 20 |

| **RL08** | 0/0 | 0/1 | 1/1 |
|---|---|---|---|
| 0/0 | 598 | 0 | 0 |
| 0/1 | 0 | 13 | 0 |
| 0/1 | 0 | 0 | 324 |

**Table S8.** Correlation ($r^2$) and $p$-value between the top two genetic PCs and the top eight environmental PCs obtained from linear regressions. There are no significant relationships at the 5% level.

| Genetic PC | Environmental PC | $r^2$ | $p$-value |
|---|---|---|---|
| PC1 | PC1 | 0.004 | 0.350 |
| PC1 | PC2 | 0.004 | 0.341 |
| PC1 | PC3 | 0.001 | 0.668 |
| PC1 | PC4 | 0.001 | 0.728 |
| PC1 | PC5 | 0.002 | 0.556 |
| PC1 | PC6 | 0.002 | 0.636 |
| PC1 | PC7 | 0.014 | 0.089 |
| PC1 | PC8 | 0.032 | 0.084 |
| PC2 | PC1 | 0.001 | 0.852 |
| PC2 | PC2 | 0.005 | 0.315 |
| PC2 | PC3 | 0.002 | 0.574 |
| PC2 | PC4 | 0.012 | 0.112 |
| PC2 | PC5 | 0.007 | 0.208 |
| PC2 | PC6 | 0.001 | 0.924 |
| PC2 | PC7 | 0.006 | 0.254 |
| PC2 | PC8 | 0.011 | 0.123 |

### 10 References

#### References

[1] S. Koren, B. P. Walenz, K. Berlin, J. R. Miller, N. H. Bergman, and A. M. Phillippy, Canu: scalable and accurate long-read assembly via adaptive k-mer weighting and repeat separation, *Genome Research* **27**, 722 (2017), ISSN 1088-9051, 1549-5469.

[2] C.-S. Chin, P. Peluso, F. J. Sedlazeck, M. Nattestad, G. T. Concepcion, A. Clum, C. Dunn, R. O'Malley, R. Figueroa-Balderas, A. Morales-Cruz, et al., Phased Diploid Genome Assembly with Single Molecule Real-Time Sequencing, *Nature methods* **13**, 1050 (2016), ISSN 1548-7091.

[3] C. Shinzato, E. Shoguchi, T. Kawashima, M. Hamada, K. Hisata, M. Tanaka, M. Fujie, M. Fujiwara, R. Koyanagi, T. Ikuta, et al., Using the Acropora digitifera genome to understand coral responses to environmental change, *Nature* **476**, 320 (2011), ISSN 0028-0836.

[4] S. Yeo, L. Coombe, R. L. Warren, J. Chu, I. Birol, and C. Sahinalp, ARCS: scaffolding genome drafts with linked reads, *Bioinformatics* **34**, 725 (2018), ISSN 1367-4803.

[5] S. Yeo, L. Coombe, J. Chu, R. L. Warren, and I. Birol, ARCS: Assembly Roundup by Chromium Scaffolding, *bioRxiv* 100750 (2017).

[6] M. Boetzer and W. Pirovano, SSPACE-LongRead: scaffolding bacterial draft genomes using long read sequence information, *BMC Bioinformatics* **15**, 211 (2014), ISSN 1471-2105.

[7] A. C. English, S. Richards, Y. Han, M. Wang, V. Vee, J. Qu, X. Qin, D. M. Muzny, J. G. Reid, K. C. Worley, et al., Mind the Gap: Upgrading Genomes with Pacific Biosciences RS Long-Read Sequencing Technology, *PLOS ONE* **7**, e47768 (2012), ISSN 1932-6203.

[8] C.-S. Chin, D. H. Alexander, P. Marks, A. A. Klammer, J. Drake, C. Heiner, A. Clum, A. Copeland, J. Huddleston, E. E. Eichler, et al., Nonhybrid, finished microbial genome assemblies from long-read SMRT sequencing data, *Nature Methods* **10**, 563 (2013), ISSN 1548-7105.

[9] B. J. Walker, T. Abeel, T. Shea, M. Priest, A. Abouelliel, S. Sakthikumar, C. A. Cuomo, Q. Zeng, J. Wortman, S. K. Young, et al., Pilon: An Integrated Tool for Comprehensive Microbial Variant Detection and Genome Assembly Improvement, *PLOS ONE* **9**, e112963 (2014), ISSN 1932-6203.

[10] H. Li and R. Durbin, Fast and accurate long-read alignment with Burrows–Wheeler transform, *Bioinformatics* **26**, 589 (2010), ISSN 1367-4803.

[11] J. C. Mieog, M. J. H. VAN Oppen, R. Berkelmans, W. T. Stam, and J. L. Olsen, Quantification of algal endosymbionts (Symbiodinium) in coral tissue using real-time PCR, *Molecular Ecology Resources* **9**, 74 (2009), ISSN 1755-098X.

[12] L. v. d. Maaten and G. Hinton, Visualizing Data using t-SNE, *Journal of Machine Learning Research* **9**, 2579 (2008), ISSN ISSN 1533-7928.

[13] S. Huang, M. Kang, and A. Xu, HaploMerger2: rebuilding both haploid sub-assemblies from high-heterozygosity diploid genome assembly, *Bioinformatics (Oxford, England)* **33**, 2577 (2017), ISSN 1367-4811.

[14] J. C. Kenyon, Models of reticulate evolution in the coral genus Acropora based on chromosome numbers: parallels with plants, *Evolution; International Journal of Organic Evolution* **51**, 756 (1997), ISSN 1558-5646.

[15] S. Wang, L. Zhang, E. Meyer, and M. V. Matz, Construction of a high-resolution genetic linkage map and comparative genome analysis for the reef-building coral Acropora millepora, *Genome Biology* **10**, R126 (2009), ISSN 1474-760X.

[16] G. B. Dixon, S. W. Davies, G. V. Aglyamova, E. Meyer, L. K. Bay, and M. V. Matz, Genomic determinants of coral heat tolerance across latitudes, *Science* **348**, 1460 (2015), ISSN 0036-8075, 1095-9203.

[17] H. Tang, X. Zhang, C. Miao, J. Zhang, R. Ming, J. C. Schnable, P. S. Schnable, E. Lyons, and J. Lu, ALLMAPS: robust scaffold ordering based on multiple maps, *Genome Biology* **16**, 3 (2015), ISSN 1465-6906.

[18] M. S. Campbell, M. Law, C. Holt, J. C. Stein, G. D. Moghe, D. E. Hufnagel, J. Lei, R. Achawanantakun, D. Jiao, C. J. Lawrence, et al., MAKER-P: a tool kit for the rapid creation, management, and quality control of plant genome annotations, *Plant Physiology* **164**, 513 (2014), ISSN 1532-2548.

[19] A. Moya, L. Huisman, E. E. Ball, D. C. Hayward, L. C. Grasso, C. M. Chua, H. N. Woo, J.-P. Gattuso,

S. Forêt, and D. J. Miller, Whole Transcriptome Analysis of the Coral Acropora millepora Reveals Complex Responses to CO2-driven Acidification during the Initiation of Calcification, *Molecular Ecology* **21**, 2440 (2012), ISSN 1365-294X.

[20] D. Bhattacharya, S. Agrawal, M. Aranda, S. Baumgarten, M. Belcaid, J. L. Drake, D. Erwin, S. Foret, R. D. Gates, D. F. Gruber, et al., Comparative genomics explains the evolutionary success of reef-forming corals (2016), `https://elifesciences.org/articles/13288`.

[21] F. A. Simão, R. M. Waterhouse, P. Ioannidis, E. V. Kriventseva, and E. M. Zdobnov, BUSCO: assessing genome assembly and annotation completeness with single-copy orthologs, *Bioinformatics* **31**, 3210 (2015), ISSN 1367-4803.

[22] J. Huerta-Cepas, K. Forslund, L. P. Coelho, D. Szklarczyk, L. J. Jensen, C. von Mering, and P. Bork, Fast Genome-Wide Functional Annotation through Orthology Assignment by eggNOG-Mapper, *Molecular Biology and Evolution* **34**, 2115 (2017), ISSN 1537-1719.

[23] H. Ying, D. C. Hayward, I. Cooke, W. Wang, A. Moya, K. R. Siemering, S. Sprungala, E. E. Ball, S. Forêt, and D. J. Miller, The Whole-Genome Sequence of the Coral Acropora millepora, *Genome Biology and Evolution* **11**, 1374 (2019).

[24] U. E. Siebeck, N. J. Marshall, A. Klüter, and O. Hoegh-Guldberg, Monitoring coral bleaching using a colour reference card, *Coral Reefs* **25**, 453 (2006), ISSN 0722-4028, 1432-0975.

[25] R. J. Ritchie, Universal chlorophyll equations for estimating chlorophylls a, b, c, and d and total chlorophylls in natural assemblages of photosynthetic organisms using acetone, methanol, or ethanol solvents, *Photosynthetica* **46**, 115 (2008), ISSN 1573-9058.

[26] R. Cunning and A. C. Baker, Not just who, but how many: the importance of partner abundance in reef coral symbioses, *Frontiers in Microbiology* **5** (2014), ISSN 1664-302X.

[27] A. M. Bolger, M. Lohse, and B. Usadel, Trimmomatic: a flexible trimmer for Illumina sequence data, *Bioinformatics* **30**, 2114 (2014), ISSN 1367-4803.

[28] H. Li, B. Handsaker, A. Wysoker, T. Fennell, J. Ruan, N. Homer, G. Marth, G. Abecasis, and R. Durbin, The Sequence Alignment/Map format and SAMtools, *Bioinformatics* **25**, 2078 (2009), ISSN 1367-4803.

[29] A. McKenna, M. Hanna, E. Banks, A. Sivachenko, K. Cibulskis, A. Kernytsky, K. Garimella, D. Altshuler, S. Gabriel, M. Daly, et al., The Genome Analysis Toolkit: A MapReduce framework for analyzing next-generation DNA sequencing data, *Genome Research* **20**, 1297 (2010), ISSN 1088-9051.

[30] S. Purcell, B. Neale, K. Todd-Brown, L. Thomas, M. A. R. Ferreira, D. Bender, J. Maller, P. Sklar, P. I. W. de Bakker, M. J. Daly, et al., PLINK: A Tool Set for Whole-Genome Association and Population-Based Linkage Analyses, *The American Journal of Human Genetics* **81**, 559 (2007), ISSN 0002-9297.

[31] S. Picelli, K. Björklund, B. Reinius, S. Sagasser, G. Winberg, and R. Sandberg, Tn5 transposase and tagmentation procedures for massively scaled sequencing projects, *Genome Research* **24**, 2033 (2014), ISSN 1088-9051.

[32] J. D. Wall, L. F. Tang, B. Zerbe, M. N. Kvale, P.-Y. Kwok, C. Schaefer, and N. Risch, Estimating genotype error rates from high-coverage next-generation sequence data, *Genome Research* **24**, 1734 (2014), ISSN 1088-9051.

[33] 1000 Genomes Project Consortium, G. R. Abecasis, D. Altshuler, A. Auton, L. D. Brooks, R. M. Durbin, R. A. Gibbs, M. E. Hurles, and G. A. McVean, A map of human genome variation from population-scale sequencing, *Nature* **467**, 1061 (2010), ISSN 1476-4687.

[34] A. Manichaikul, J. C. Mychaleckyj, S. S. Rich, K. Daly, M. Sale, and W.-M. Chen, Robust relationship inference in genome-wide association studies, *Bioinformatics (Oxford, England)* **26**, 2867 (2010), ISSN 1367-4811.

[35] H. Li and R. Durbin, Inference of human population history from individual whole-genome sequences, *Nature* **475**, 493 (2011), ISSN 0028-0836, 1476-4687.

[36] S. Schiffels and R. Durbin, Inferring human population size and separation history from multiple genome sequences, *Nature genetics* **46**, 919 (2014), ISSN 1061-4036.

[37] D. Petkova, J. Novembre, and M. Stephens, Visualizing spatial population structure with estimated effective migration surfaces, *Nature Genetics* **48**, 94 (2016), ISSN 1546-1718.

[38] J. Novembre and B. M. Peter, Recent advances in the study of fine-scale population structure in humans, *Current Opinion in Genetics & Development* **41**, 98 (2016), ISSN 1879-0380.

[39] O. Delaneau, B. Howie, A. J. Cox, J.-F. Zagury, and J. Marchini, Haplotype estimation using sequencing reads, *American Journal of Human Genetics* **93**, 687 (2013), ISSN 1537-6605.

[40] N. R. Garud, P. W. Messer, E. O. Buzbas, and D. A. Petrov, Recent Selective Sweeps in North American Drosophila melanogaster Show Signatures of Soft Sweeps, *PLOS Genetics* **11**, e1005004 (2015), ISSN 1553-7404.

[41] M. W. Hahn, S. V. Zhang, and L. C. Moyle, Sequencing, assembling, and correcting draft genomes using recombinant populations, *G3 (Bethesda, Md.)* **4**, 669 (2014), ISSN 2160-1836.

[42] A. Bankevich, S. Nurk, D. Antipov, A. A. Gurevich, M. Dvorkin, A. S. Kulikov, V. M. Lesin, S. I. Nikolenko, S. Pham, A. D. Prjibelski, et al., SPAdes: A New Genome Assembly Algorithm and Its Applications to Single-Cell Sequencing, *Journal of Computational Biology* **19**, 455 (2012), ISSN 1066-5277.

[43] N. Li and M. Stephens, Modeling linkage disequilibrium and identifying recombination hotspots using single-nucleotide polymorphism data, *Genetics* **165**, 2213 (2003), ISSN 0016-6731.

[44] B. L. Browning, Y. Zhou, and S. R. Browning, A One-Penny Imputed Genome from Next-Generation Reference Panels, *American Journal of Human Genetics* **103**, 338 (2018), ISSN 1537-6605.

[45] L. Huang, Y. Li, A. B. Singleton, J. A. Hardy, G. Abecasis, N. A. Rosenberg, and P. Scheet, Genotype-imputation accuracy across worldwide human populations, *American Journal of Human Genetics* **84**, 235 (2009), ISSN 1537-6605.

[46] B. Howie, C. Fuchsberger, M. Stephens, J. Marchini, and G. R. Abecasis, Fast and accurate genotype imputation in genome-wide association studies through pre-phasing, *Nature Genetics* **44**, 955 (2012), ISSN 1546-1718.

[47] H.-Y. Tsai, O. Matika, S. M. Edwards, R. Antolín–Sánchez, A. Hamilton, D. R. Guy, A. E. Tinch, K. Gharbi, M. J. Stear, J. B. Taggart, et al., Genotype Imputation To Improve the Cost-Efficiency of Genomic Selection in Farmed Atlantic Salmon, *G3: Genes, Genomes, Genetics* **7**, 1377 (2017), ISSN 2160-1836.

[48] T. Druet, I. M. Macleod, and B. J. Hayes, Toward genomic prediction from whole-genome sequence data: impact of sequencing design on genotype imputation and accuracy of predictions, *Heredity* **112**, 39 (2014), ISSN 1365-2540.

[49] M. Schumer, C. Xu, D. L. Powell, A. Durvasula, L. Skov, C. Holland, J. C. Blazier, S. Sankararaman, P. Andolfatto, G. G. Rosenthal, et al., Natural selection interacts with recombination to shape the evolution of hybrid genomes, *Science* **360**, 656 (2018), ISSN 0036-8075, 1095-9203.

[50] T. S. Korneliussen, A. Albrechtsen, and R. Nielsen, ANGSD: Analysis of Next Generation Sequencing Data, *BMC bioinformatics* **15**, 356 (2014), ISSN 1471-2105.

[51] D. P. Manzello, M. V. Matz, I. C. Enochs, L. Valentino, R. D. Carlton, G. Kolodziej, X. Serrano, E. K. Towle, and M. Jankulak, Role of host genetics and heat-tolerant algal symbionts in sustaining populations of the endangered coral Orbicella faveolata in the Florida Keys with ocean warming, *Global Change Biology* **25**, 1016 (2019), ISSN 1365-2486.

[52] E. Shoguchi, C. Shinzato, T. Kawashima, F. Gyoja, S. Mungpakdee, R. Koyanagi, T. Takeuchi, K. Hisata, M. Tanaka, M. Fujiwara, et al., Draft Assembly of the Symbiodinium minutum Nuclear Genome Reveals Dinoflagellate Gene Structure, *Current Biology* **23**, 1399 (2013), ISSN 0960-9822.

[53] H. Liu, T. G. Stephens, R. González-Pech, V. H. Beltran, B. Lapeyre, P. Bongaerts, I. Cooke, D. G. Bourne, S. Forêt, D. J. Miller, et al., Symbiodinium genomes reveal adaptive evolution of functions related to symbiosis, *bioRxiv* 198762 (2017).

[54] L. C. Xia, J. A. Cram, T. Chen, J. A. Fuhrman, and F. Sun, Accurate Genome Relative Abundance Estimation Based on Shotgun Metagenomic Reads, *PLoS ONE* **6** (2011), ISSN 1932-6203.

[55] J. Yang, S. H. Lee, M. E. Goddard, and P. M. Visscher, GCTA: A Tool for Genome-wide Complex Trait Analysis, *American Journal of Human Genetics* **88**, 76 (2011), ISSN 0002-9297.

[56] P. M. Visscher, G. Hemani, A. A. E. Vinkhuyzen, G.-B. Chen, S. H. Lee, N. R. Wray, M. E. Goddard, and J. Yang, Statistical Power to Detect Genetic (Co)Variance of Complex Traits Using SNP Data in Unrelated Samples, *PLOS Genetics* **10**, e1004269 (2014), ISSN 1553-7404.

[57] X. Zhou, A UNIFIED FRAMEWORK FOR VARIANCE COMPONENT ESTIMATION WITH SUMMARY STATISTICS IN GENOME-WIDE ASSOCIATION STUDIES, *The Annals of Applied Statistics* **11**, 2027 (2017), ISSN 1932-6157.

[58] X. Zhou and M. Stephens, Genome-wide efficient mixed-model analysis for association studies, *Nature Genetics* **44**, 821 (2012), ISSN 1546-1718.

[59] B. Servin and M. Stephens, Imputation-Based Analysis of Association Studies: Candidate Regions and Quantitative Traits, *PLOS Genetics* **3**, e114 (2007), ISSN 1553-7404.

[60] W. K. Meyer, B. Arbeithuber, C. Ober, T. Ebner, I. Tiemann-Boege, R. R. Hudson, and M. Przeworski, Evaluating the Evidence for Transmission Distortion in Human Pedigrees, *Genetics* **191**, 215 (2012), ISSN 0016-6731, 1943-2631.

[61] A. R. Martin, M. Kanai, Y. Kamatani, Y. Okada, B. M. Neale, and M. J. Daly, Clinical use of current polygenic risk scores may exacerbate health disparities, *Nature Genetics* **51**, 584 (2019), ISSN 1546-1718.

[62] F. Dudbridge, Power and Predictive Accuracy of Polygenic Risk Scores, *PLOS Genetics* **9**, e1003348 (2013), ISSN 1553-7404.
